# The AP-1 clathrin adaptor has a conserved role in recycling resident proteins of the *trans*-Golgi network

**DOI:** 10.64898/2026.10.21.760840

**Authors:** Fernando M. Valbuena, Maddison Lessard, Adam H. Krahn, Areti Pantazopoulou, Conor F. Lee-Smith, J. Christopher Fromme, Petia Adarska, Francesca Bottanelli, Benjamin S. Glick

## Abstract

Although the AP-1 clathrin adaptor has been studied for decades, its role in membrane trafficking has been ambiguous. Here, we systematically investigated the localization and function of mammalian AP-1. This adaptor associates with the *trans*-Golgi network (TGN) and with TGN-related recycling endosomes. Disruption of the µ1A subunit of AP-1 has only a mild effect on the Golgi export of secretory cargoes, but it causes depletion from the Golgi of multiple transmembrane TGN proteins. Live-cell imaging and a cell surface labeling assay indicate that when TGN proteins are unable to bind AP-1, they exit the Golgi to the plasma membrane. Our interpretation is that the vesicles generated with AP-1 recycle transmembrane TGN proteins, thereby keeping those proteins in the Golgi during cisternal maturation. The clathrin adaptor epsinR seems to function as a partner for AP-1 in recycling a subset of TGN proteins. These results closely align with studies of AP-1 function in yeast, suggesting that the conserved core role of AP-1 is the intra-Golgi recycling of TGN proteins.

## Introduction

The heterotetrameric AP-1 and AP-2 adaptor complexes aid the formation of clathrin-coated vesicles (Robinson, 2015). These adaptors are peripheral membrane proteins that bind the cytoplasmic tails of cargo proteins while associating with clathrin. AP-2 acts at the plasma membrane to promote the formation of endocytic vesicles. Binding of AP-2 to cargo and to the plasma membrane lipid phosphatidylinositol 4,5-bisphosphate triggers a conformational change that enables recruitment of clathrin (Paczkowski et al., 2015).

Although AP-1 is closely related to AP-2, the roles of AP-1 have been harder to discern (Tan and Gleeson, 2019; Duncan, 2022). A pool of mammalian AP-1 is found at the *trans*-Golgi network (TGN). Recruitment of AP-1 to the TGN requires the GTPase Arf1 together with the lipid phosphatidylinositol 4-phosphate (PI4P) (Stamnes and Rothman, 1993; Wang et al., 2003; Heldwein et al., 2004; Ren et al., 2013). Transmembrane cargo proteins may also play a role in the recruitment and assembly of AP-1 (Deng et al., 2026). The original hypothesis was that AP-1 operates at the TGN to capture the cation-independent and cation-dependent mannose 6-phosphate receptors (CIMPR and CDMPR) and their bound lysosomal hydrolases into clathrin-coated vesicles for delivery to endosomes (Ghosh et al., 2003). However, that model struggled to explain why loss of AP-1 depletes the Golgi of some TGN proteins, including CIMPR and CDMPR (Meyer et al., 2000; Robinson et al., 2010; Hirst et al., 2012). A further challenge to the original hypothesis about AP-1 was the discovery that the GGA clathrin adaptors mediate TGN-to-endosome delivery of mannose 6-phosphate receptors and their bound lysosomal hydrolases (Dell’Angelica et al., 2000; Hirst et al., 2000; Ghosh et al., 2003). The characterization of GGAs raised the question of why AP-1 is needed. A possible interpretation is that AP-1 does not mediate anterograde traffic from the TGN but instead mediates retrograde traffic from post-Golgi compartments back to the TGN (Hinners and Tooze, 2003; Hirst et al., 2012). In support of that idea, mammalian AP-1 is present not only on the TGN but also on recycling endosomes, and AP-1 seems to play a role in traffic from recycling endosomes to the TGN (Guo et al., 2013; Matsudaira et al., 2015; Stockhammer et al., 2024). Robinson and colleagues recently concluded that the primary function of mammalian AP-1 is retrieval of resident TGN proteins from post-Golgi carriers and/or recycling endosomes (Robinson et al., 2024). However, others have proposed that AP-1 acts at early or late endosomes to mediate the recycling of mannose 6-phosphate receptors to the TGN (Meyer et al., 2000; Brown et al., 2011; Buser et al., 2018) or to promote endosome maturation (Candiello et al., 2016). It was reported that an effector of the Rab4 GTPase binds to an AP-1 subunit (Deneka et al., 2003) and that AP-1 is present on Rab4-containing tubular domains of early endosomes (D’Souza et al., 2014). Further complexity comes from the finding that inactivation of mammalian AP-1 causes certain secretory cargoes to accumulate in the Golgi, leading to the idea that AP-1 acts at the TGN to support anterograde traffic to the plasma membrane (Duncan, 2022). The bottom line is that there is no consensus about where mammalian AP-1 operates or which traffic pathways it supports.

Insight into these questions has come from the budding yeast *Saccharomyces cerevisiae*. This yeast contains non-stacked Golgi cisternae that progressively mature, and the TGN phase of maturation is a large fraction of the cisternal lifetime (Losev et al., 2006; Tojima et al., 2019). In a process analogous to the sorting of mammalian lysosomal hydrolases, yeast vacuolar hydrolases bind to the Vps10 receptor in the TGN, and those complexes travel to prevacuolar endosomes in a GGA-dependent pathway (Bowers and Stevens, 2005). Kinetic analysis of individual Golgi cisternae revealed that GGA-dependent departure of Vps10 occurs about halfway through maturation (Casler and Glick, 2020; Glick, 2026). Downstream of the GGAs, yeast AP-1 arrives at maturing Golgi cisternae with the aid of Arf1, additional partner proteins, and phosphatidylinositol 4-phosphate (Daboussi et al., 2012; Zysnarski et al., 2019; Highland and Fromme, 2021). There is no indication that yeast AP-1 mediates the traffic of vacuolar hydrolases (Stepp et al., 1995; Yeung et al., 1999; Casler and Glick, 2020). Instead, yeast AP-1 has been implicated in the retrograde traffic of resident TGN proteins (Valdivia et al., 2002; Liu et al., 2008; Casler et al., 2022; Krahn et al., 2026b). We have shown that yeast AP-1 resides exclusively on maturing TGN cisternae, where it promotes the intra-Golgi recycling of TGN proteins (Day et al., 2018; Casler et al., 2022). Historically, yeast TGN proteins were thought to recycle from prevacuolar endosomes, but recent findings suggest that the primary traffic pathway for many yeast TGN proteins is actually intra-Golgi recycling (Glick, 2026; Krahn et al., 2026b). An attractive idea is that the yeast pathway of AP-1-dependent intra-Golgi retrograde traffic resembles the mammalian pathway of AP-1-dependent retrograde traffic from recycling endosomes to the TGN. This model is bolstered by accumulating evidence that recycling endosomes are closely related to the TGN and may be TGN-derived compartments (Grant and Donaldson, 2009; Fujii et al., 2020; Stockhammer et al., 2024; Satoh, 2026).

In addition to AP-1 and the GGAs, epsinR acts as a clathrin adaptor at the mammalian TGN (Hirst et al., 2015). epsinR binds to and colocalizes with AP-1, suggesting that the functions of these two adaptors are linked (Hirst et al., 2003; Mills et al., 2003). The only established epsinR cargo is the SNARE protein Vti1b, although the paralogous protein Vti1a also requires epsinR for normal localization (Chidambaram et al., 2004; Hirst et al., 2004). There is evidence that epsinR has a broader involvement in clathrin-coated vesicle formation (Hirst et al., 2015), but the specific role of epsinR has been uncertain. Budding yeast has three epsinR homologs: Ent3, Ent4, and Ent5 (Myers and Payne, 2013). Ent3 arrives at maturing Golgi cisternae together with the GGAs and binds Vti1, the yeast ortholog of Vti1a and Vti1b, for delivery to prevacuolar endosomes (Chidambaram et al., 2004). The poorly characterized Ent4 has also been implicated in Golgi-to-prevacuolar endosome traffic (Deng et al., 2009). Therefore, Ent3 and perhaps Ent4 cooperate with the GGAs. Ent5 arrives together with AP-1 and functions with AP-1 (Duncan et al., 2003; Costaguta et al., 2006; Čopič et al., 2007; Casler et al., 2022). A complete view of the trafficking pathways mediated by AP-1 and the GGAs will require elucidating the contributions of epsinR and its homologs.

The main focus of this study is AP-1. Yeast experiments have led to the simple view that AP-1 (in cooperation with Ent5) mediates the continual recycling of TGN proteins, thereby keeping those proteins in place during Golgi cisternal maturation (Pantazopoulou and Glick, 2019; Casler et al., 2022; Glick, 2026). Our working hypothesis is that mammalian AP-1 performs an analogous role by recycling TGN proteins from terminally maturing TGN compartments and from TGN-derived recycling endosomes. Here, we have tested this hypothesis by characterizing mammalian AP-1 with an emphasis on the parallels between mammalian and yeast cells.

## Results

### Mammalian cells with disrupted AP-1 function have mild defects in cargo export from the Golgi

We chose to examine RPE-1 cells, which are derived from human retinal pigment epithelium, because they are stably diploid and amenable to fluorescence microscopy (Bodnar et al., 1998; Tormanen et al., 2019; Volpe et al., 2023). A "Flp-In" version of this cell line was generated to enable Flp recombinase-mediated stable integration of an expression vector into the genome (O’Gorman et al., 1991). Whole-genome sequencing confirmed that a single copy of the Flp recombination target site was present in chromosome 2 (see Methods). The original and Flp-In versions of the RPE-1 cell line are here designated "WT".

The original RPE-1 cell line was then mutated by using CRISPR/Cas9-mediated genome editing to knock out specific genes (Wang and Doudna, 2023). To create a derivative lacking AP-1, we knocked out the AP1M1 gene, which encodes the μ1A subunit, by introducing frameshift mutations (Figure S1A). Loss of µ1A was confirmed by immunoblotting (Figure S1C). It was expected that loss of μ1A would disrupt assembly of the heterotetrameric AP-1 complex (Meyer et al., 2000). Indeed, based on immunofluorescence analysis of RPE-1 cells lacking μ1A, the Golgi signal for AP-1 as detected by visualizing the γ1 subunit (AP1G1) was largely abolished even though the Golgi ribbon as detected by visualizing the *cis*-Golgi marker GM130 (Nakamura et al., 1995) appeared normal (Figure 1A). The loss of Golgi-associated AP-1 did not noticeably impede cell growth. Our RPE-1-derived µ1A knockout cell line is here designated "µ1A KO".

**Figure 1.**
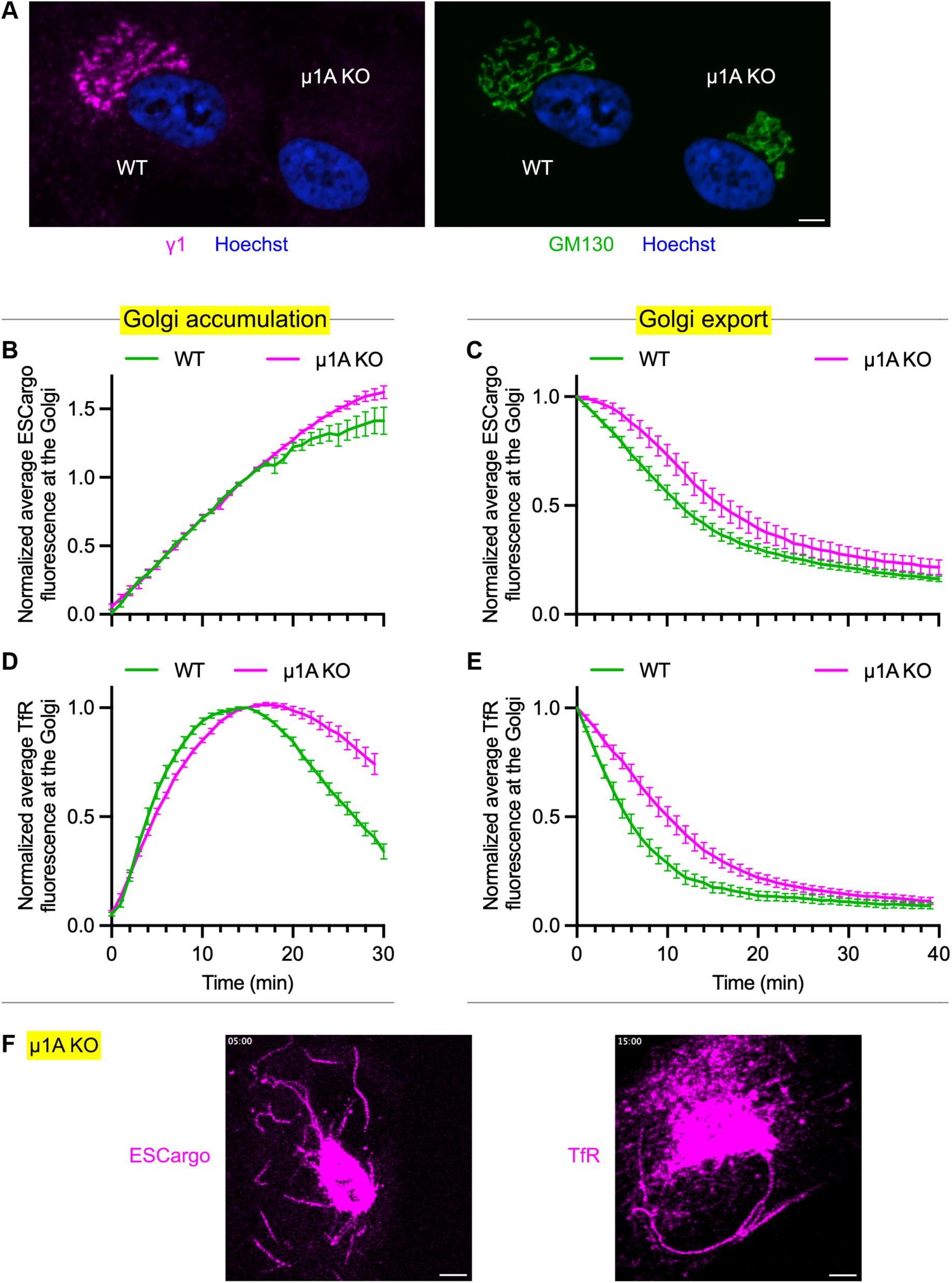
Loss of AP-1 mildly alters Golgi structure and function. **(A)** Confirmation that RPE-1 cells with a disrupted µ1A gene lack detectable Golgi-associated AP-1. Equal numbers of WT and µ1A KO cells were seeded onto a coverslip and grown overnight. The following day, the cells were processed for immunofluorescence to detect the γ1 subunit of AP-1 (magenta) together with GM130 (green). Nuclei were stained with Hoechst dye (blue). In a separate experiment that examined coverslips seeded purely with either WT or µ1A KO cells, essentially all of the WT cells and none of the mutant cells showed Golgi signals for γ1. For the mixed culture used here, approximately half of the cells showed Golgi signals for γ1, indicating that WT cells and mutant cells could be distinguished in this manner. Shown is an example of a presumptive WT cell next to a presumptive µ1A KO cell. Scale bar, 5 µm. **(B)** Effect of AP-1 inactivation on the accumulation of a soluble secretory cargo in the Golgi after release from the ER. Cells were transiently transfected to express GalNAc-T2-HaloTag plus ESCargo, and GalNAc-T2-HaloTag was labeled with JFX_650_ to mark the Golgi. ESCargo was released from the ER by addition of SLF, and accumulation of ESCargo in the Golgi was quantified at 1-min intervals by measuring the red fluorescence that overlapped with the Golgi as marked by GalNAc-T2-HaloTag. This analysis was performed for 9 WT cells (green) on 2 different days from a total of 4 separate dishes, and for 9 µ1A KO cells (magenta) on 2 different days from a total of 4 separate dishes. The plots for the individual cells were normalized to 1.0 at the 15-min time point and then averaged. Shown are mean and SEM values at each time point. Representative movies for the two cell lines are shown in Video 1. **(C)** Effect of AP-1 inactivation on the export of a soluble secretory cargo from the Golgi. Cells were transiently transfected to express GalNAc-T2-HaloTag plus ESCargo, and GalNAc-T2-HaloTag was labeled with JFX_650_ to mark the Golgi. ESCargo was released from the ER by addition of SLF, and the cells were incubated for 10-15 min to accumulate ESCargo in the Golgi. Then non-Golgi ESCargo fluorescence was photobleached, and export of unbleached ESCargo from the Golgi was tracked by measuring the remaining red fluorescence that overlapped with the Golgi as marked by GalNAc-T2-HaloTag. The post-photobleaching quantification of Golgi export began 18-20 min after SLF addition. This analysis was performed for 15 WT cells (green) on 2 different days from a total of 4 separate dishes, and for 11 µ1A KO cells (magenta) on 3 different days from a total of 5 separate dishes. The plots for the individual cells were normalized to 1.0 at time zero and then averaged. Shown are mean and SEM values at each time point. Representative movies for the two cell lines are shown in Video 2. **(D)** Effect of AP-1 inactivation on the accumulation of a transmembrane secretory cargo in the Golgi after release from the ER. The analysis was performed as in (B), except that SBP-mScarlet-tagged TfR was released from the ER by addition of biotin. Accumulation of tagged TfR in the Golgi was analyzed for 15 WT cells (green) on 2 different days from a total of 4 separate dishes, and for 18 µ1A KO cells (magenta) on 2 different days from a total of 4 separate dishes. Representative movies for the two cell lines are shown in Video 3. **(E)** Effect of AP-1 inactivation on the export of a transmembrane secretory cargo from the Golgi. The analysis was performed as in (C), except that Golgi export of SBP-mScarlet-tagged TfR was analyzed for 7 WT cells (green) on 2 different days from a total of 2 separate dishes, and for 11 µ1A KO cells (magenta) on 2 different days from a total of 4 separate dishes. Representative movies for the two cell lines are shown in Video 4. **(F)** Visualization of secretory cargo-containing tubules in µ1A KO cells. Representative frames were taken from the iFRAP movies of µ1A KO cells in Videos 2 and 4, and the indicated cargo signals were enhanced to highlight the Golgi-associated tubules. For comparison, tubules were rarely visible in the iFRAP movies of WT cells. Scale bar, 5 µm.

Humans have an AP1M1-related gene called AP1M2 that encodes the μ1B subunit (Ohno et al., 1999; Fölsch, 2015). µ1B is expressed in many epithelial cell types, but it was reported that only μ1A was expressed in retinal pigment epithelium-derived cells (Diaz et al., 2009). We used reverse transcription quantitative PCR (RT-qPCR) to examine µ1A and µ1B expression in RPE-1 cells. As a control, a parallel analysis was performed with human bronchial epithelial (HBE) cells, which express both µ1A and µ1B (Nokes et al., 2008). Surprisingly, transcripts for both µ1A and µ1B were detected not only in HBE cells, but also in RPE-1 cells (Figure S1B). Immunoblotting confirmed that µ1B was present in HBE cells and, at a lower level, in RPE-1 cells (Figure S1C). When µ1A was knocked out in RPE-1 cells, µ1B was no longer detected (Figure S1C), suggesting that µ1B stability might depend on µ1A. For comparison, no µ1B was detected in wild-type U2OS cells (Pontén and Saksela, 1967) or in a U2OS derivative with knockouts of both µ1A and µ1B (Figure S1C). This result with U2OS cells was expected because µ1B is generally absent from non-epithelial cells (Fölsch, 2015). The role of µ1B and its relationship to µ1A merit further analysis, but for our purposes, the µ1A KO is apparently sufficient to cause extensive and perhaps complete inactivation of AP-1 as judged by the loss of γ1 from the Golgi.

The robust growth of µ1A KO cells suggested that secretion was occurring efficiently. We set out to verify this point using two model secretory cargoes. Traffic of the secretory cargoes was initially visualized in WT cells. The first cargo examined was ESCargo, a red fluorescent secretory protein that can be trapped as aggregates in the ER lumen and then solubilized for rapid delivery to the Golgi by adding the small molecule SLF (Casler et al., 2020). Accumulation of ESCargo in the Golgi after SLF addition was quantified. The data for each cell were normalized to the signal at the 15-min time point, when substantial Golgi accumulation had occurred but Golgi export had not yet begun. During a 30-min time course, ESCargo progressively accumulated in the Golgi (Video 1 and Figure 1B). Golgi export was then examined by a separate method. At 8-12 min after SLF addition, when many of the ESCargo molecules had reached the Golgi, we isolated this Golgi pool by photobleaching ESCargo molecules outside the Golgi, and then we measured the time-dependent loss of fluorescent ESCargo from the Golgi. This procedure is known as inverse fluorescence recovery after photobleaching (iFRAP) (Patterson et al., 2008). It was facilitated by using a version of ESCargo based on DsRed-Max, which can be photobleached efficiently (Strack et al., 2008). After photobleaching the non-Golgi pool, we tracked ESCargo fluorescence in the Golgi. Imaging started 18 min after SLF addition because that time point was approximately the beginning of the export phase. In this case, the data were normalized to the starting signal for the Golgi pool that had been isolated by photobleaching. During a 40-min time course, most of the unbleached ESCargo molecules departed from the Golgi (Video 2 and Figure 1C). The second cargo examined was the transferrin receptor (TfR), modified for use in the RUSH ("Retention Using Selective Hooks") system of regulated cargo exit from the ER (Boncompain et al., 2012; Boncompain and Perez, 2013). In this system, a construct containing a luminal streptavidin-binding peptide (SBP) is retained in the ER in cells expressing streptavidin-KDEL, and then the construct is released by addition of biotin to generate a wave of cargo protein entering the Golgi. TfR was tagged with SBP and with the red fluorescent protein mScarlet-2I, which can be photobleached efficiently (Valbuena et al., 2020). Addition of biotin caused mScarlet-TfR to exit the ER and accumulate in the Golgi (Video 3 and Figure 1D). As with ESCargo, photobleaching was then performed to isolate this Golgi pool, and we tracked the unbleached TfR molecules as they departed from the Golgi (Video 4 and Figure 1E). The results were similar to those seen with ESCargo, except that Golgi export of TfR was significantly faster.

Then we compared WT cells with µ1A KO cells by examining the kinetics of Golgi accumulation after release from the ER as well as the kinetics of Golgi export. For ESCargo, the kinetics of accumulation in the Golgi were similar in WT and µ1A KO cells for the first 15 min, but then the Golgi signal began to level off sooner for WT cells (Figure 1B). A similar but more pronounced effect of µ1A disruption was seen for TfR. After 15 min, the Golgi signal for TfR began to decline in WT cells, but this decline was more gradual in µ1A KO cells (Figure 1D). Thus, when Golgi accumulation after release from the ER was examined, both cargoes reached their maximal Golgi signals later in µ1A KO cells. The reason became apparent when we compared Golgi export kinetics in WT versus µ1A KO cells. For the two cargoes, Golgi export was approximately 1.5-to 2-fold slower in µ1A KO cells (Videos 2 and 4 and Figure 1C,E). Bottanelli and colleagues reported an apparently more severe delay when they quantified how loss of µ1A affected Golgi export (see Figure 7h in Stockhammer et al., 2024), but this discrepancy likely reflects their use of a Pearson correlation coefficient with a standard RUSH protocol versus our measurement of the residual Golgi signal with an iFRAP protocol. Extensive cargo-containing tubules were often visible in µ1A KO cells, as illustrated by boosting the cargo signals in frames from the iFRAP movies in Videos 2 and 4 (Figure 1F). Similar Golgi tubulation was previously seen in cells lacking AP-1 (Stockhammer et al., 2024; Takiguchi et al., 2026). We conclude that the structure and function of the TGN are somewhat altered in µ1A KO cells but that the effects on secretory cargo traffic are mild.

### Mammalian AP-1 localizes to the TGN and recycling endosomes

There had not previously been a quantitative analysis of the localization of endogenous AP-1 in cultured mammalian cells, so we set out to fill that gap. AP-1 was detected by an immunofluorescence method that efficiently preserves cellular structures (Bhattacharyya et al., 2010) using a monoclonal antibody against γ1. The signal obtained with this antibody was specific because it was abolished when the AP-1 complex was disrupted by loss of µ1A (see Figure 1A). As shown in Figure 2A, AP-1 exhibited close association with golgin-97 (GOLGA1), a marker for the TGN (Barr, 1999; Kjer-Nielsen et al., 1999; Munro and Nichols, 1999).

**Figure 2.**
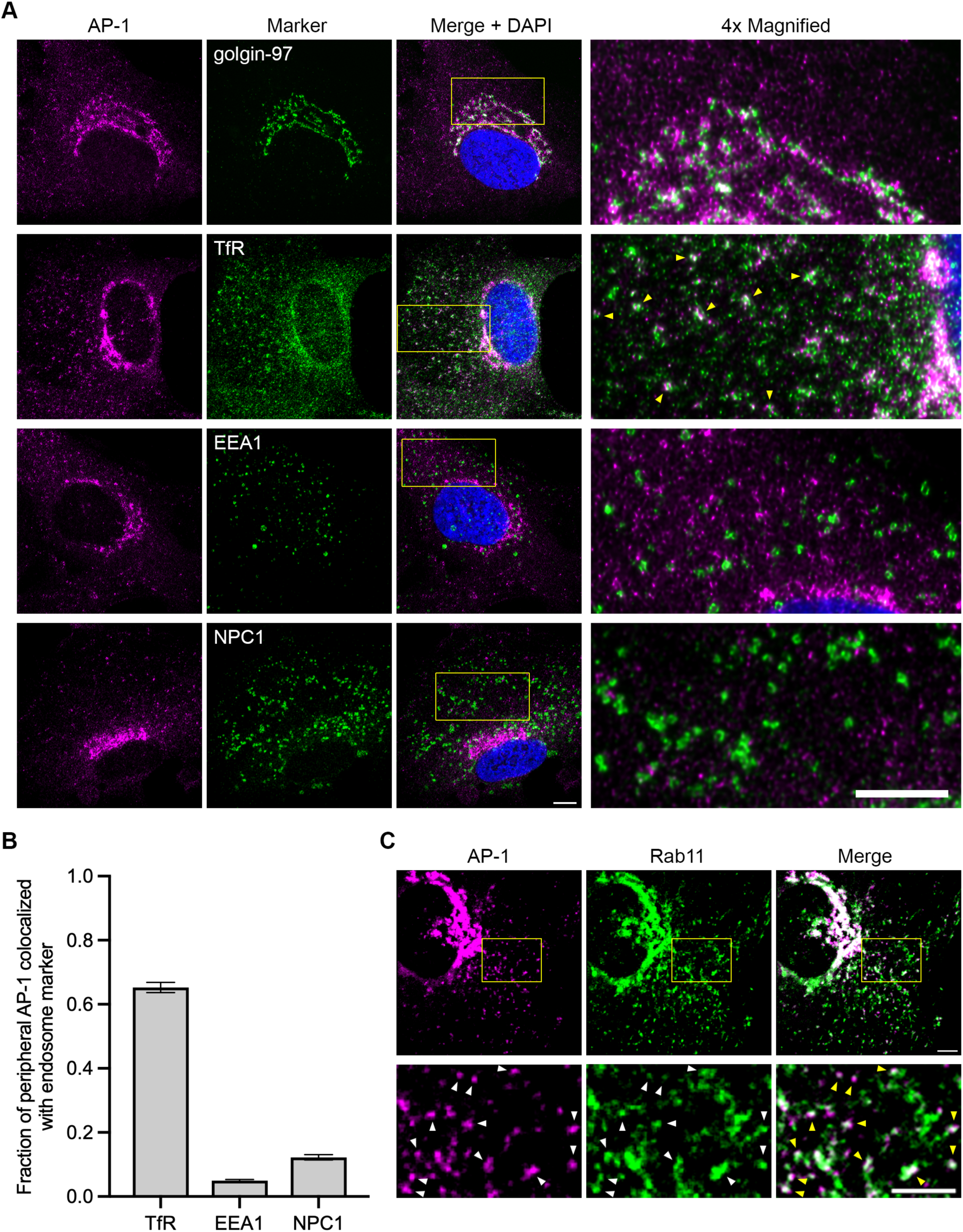
AP-1 localizes to the TGN and recycling endosome. **(A)** Representative immunofluorescence images of AP-1 together with various organelle markers in RPE-1 cells. The γ1 subunit of AP-1 is shown in magenta, the indicated organelle markers are shown in green, and DAPI-labeled nuclei are shown in blue. Golgin-97 marks the TGN, TfR marks recycling endosomes, EEA1 marks early endosomes, and NPC1 marks late endosomes. The areas in the yellow boxes are magnified 4x in the right column, where arrowheads in the second magnified image indicate examples of structures that contain both γ1 and TfR. Scale bars, 5 µm. **(B)** Ǫuantification of the data from (A). Deconvolved z-stacks were average projected. An ROI was drawn around the nucleus and Golgi using the AP-1 γ1 signal as a reference, and the signal from this ROI was erased so that the analysis was restricted to peripheral endosomes. The γ1 signal in the red channel was then thresholded in ImageJ so that only the pixels in the highest 2% of the intensity histogram were preserved, thereby eliminating most of the diffuse background signal while preserving most of the punctate signal. A mask was created from the punctate endosomal structures in the green channel, and the fraction of the thresholded red signal that overlapped with the green mask was measured. Plotted are the mean and SEM values obtained by analyzing cells on 2 different days, from a total of 4 wells on a coverslip for each marker. The total number of cells analyzed was 38 for TfR, 44 for EEA1, and 43 for NPC1. **(C)** Representative images of live cells expressing endogenously tagged µ1A and Rab11. µ1A labeled with SNAP-tag and JFX_554_ SNAP-tag ligand is shown in magenta, and Rab11 labeled with HaloTag and JFX_650_ HaloTag ligand is shown in green as a recycling endosome marker. The area in the yellow box is magnified 3.17-fold in the bottom row, where arrowheads indicate examples of structures that contain both µ1A and Rab11. Scale bars, 5 µm.

In addition to the TGN signal, AP-1 was detected on peripheral punctate structures that were presumably endosomes. This phenomenon has been reported previously, but the identities of the AP-1-positive endosomes were not always clear (Meyer et al., 2000; Waguri et al., 2003; Tan and Gleeson, 2019; Buser and Spang, 2023). Mammalian endosomes can be assigned to distinct types based on characteristic markers (Scott et al., 2014; Naslavsky and Caplan, 2018). Here, we define recycling endosomes as structures marked by the transferrin receptor (TfR), and we define early endosomes as structures marked by EEA1, and we define late endosomes as structures marked by NPC1. In accord with earlier findings (Guo et al., 2013), peripheral AP-1 puncta showed substantial association with the recycling endosome marker TfR (Figure 2A). As shown in the magnified image in the second row of Figure 2A, AP-1 and TfR often appeared to mark adjacent or partially overlapping domains of the same peripheral structures. By contrast, there was no consistent colocalization with the early endosome marker EEA1 or the late endosome marker NPC1 (Figure 2A). To quantify the colocalization data for the AP-1-positive endosomes, each immunofluorescence image was processed as follows. First, the area containing the nucleus and Golgi was excluded. Then the AP-1 channel was subjected to thresholding to remove the background non-punctate signal, leaving only the peripheral punctate AP-1 signal. Finally, a mask was created from a reference endosome marker, either TfR or EEA1 or NPC1, and the fraction of the peripheral punctate AP-1 signal that overlapped with that mask was quantified. As expected, the colocalization of AP-1 was high with the recycling endosome marker TfR and much lower with the early endosome marker EEA1 and the late endosome marker NPC1 (Figure 2B). These results suggest that the peripheral punctate structures labeled with AP-1 are recycling endosomes.

To verify this conclusion, we used an RPE-1 derivative in which the endogenous μ1A gene had been modified to encode a fusion to SNAP-tag (Stockhammer et al., 2024) (Figure S1C). That cell line was further engineered so that the endogenous gene encoding Rab11a, a canonical recycling endosome marker (Goldenring, 2015), was modified to encode a fusion to HaloTag (Stockhammer et al., 2024). Double labeling of live cells with a far-red SNAP-tag ligand and an orange HaloTag ligand (Grimm et al., 2021) confirmed that virtually all of the peripheral punctate μ1A-SNAP-tag-containing structures were also positive for HaloTag-Rab11a (Figure 2C). The shapes of the µ1A-containing domains did not always match those of the Rab11a-containing domains (Figure 2C), suggesting that these two proteins were sometimes concentrated in different regions of the same membrane compartments. The combined results indicate that mammalian AP-1 is present on the TGN and on specific domains of recycling endosomes but is largely or entirely absent from early and late endosomes.

As a further test of this interpretation, we took advantage of the finding that the multi-subunit EARP tether is evidently restricted to recycling endosomes (Schindler et al., 2015). EARP is related to GARP, a multi-subunit tether of the late Golgi (Bonifacino and Hierro, 2011), but EARP contains the VPS50 subunit instead of VPS54 in GARP. Immunofluorescence of VPS50 revealed scattered punctate structures that were presumably recycling endosomes (Figure S2A). The labeling with this antibody was variable between experiments, but when the labeling was successful, there was frequent colocalization or close juxtaposition between peripheral VPS50-containing structures and AP-1-containing puncta (Figure S2A). Thus, peripherally localized AP-1 shows clear association with three different recycling endosome markers.

Our finding that the peripheral AP-1-containing structures in RPE-1 cells are recycling endosomes is at odds with a report that in the simian fibroblast-like cell line COS-7 (Gluzman, 1981), AP-1 was present on Rab4-containing tubular domains of early endosomes (D’Souza et al., 2014). That report was ambiguous because Rab4 also marks recycling endosomes (Deneka et al., 2003; Schindler et al., 2015), but we considered the possibility that AP-1 might localize differently in COS-7 cells than in RPE-1 cells. Therefore, we tested COS-7 cells by immunofluorescence. In COS-7 cells, AP-1 was present in the Golgi region and on domains of TfR-containing recycling endosomes, but AP-1 was not detectably associated with EEA1-containing early endosomes (Figure S2B). Similar results were obtained with the immortalized human fibroblast cell line BJ-5ta (Bodnar et al., 1998), which has unusually large early endosomes (Figure S2C). Thus, data from various cell lines indicate that in the cell periphery, AP-1 localizes mainly or exclusively to recycling endosomes.

Does AP-1 act primarily at the TGN or primarily at recycling endosomes? To address this question, we measured the fraction of the membrane-associated AP-1 signal in the Golgi region versus the fraction in peripheral puncta (see Methods). Approximately 41% of the membrane-associated AP-1 was in the Golgi region while the remaining 59% was in peripheral puncta. Assuming that localization reflects function, AP-1 has roles both at the TGN and at recycling endosomes.

### Loss of AP-1 in RPE-1 cells selectively depletes certain transmembrane proteins from the Golgi

Previous studies reported that inactivation of mammalian AP-1 resulted in depletion from the Golgi of various transmembrane TGN proteins (Meyer et al., 2000; Harasaki et al., 2005; Hirst et al., 2005; Ishizaki et al., 2008; Hirst et al., 2012; Bonnemaison et al., 2014), but this phenomenon has not been investigated systematically or quantitatively. We therefore used immunofluorescence of WT and µ1A KO cells (see Methods) to measure the Golgi localization of the proteins described below.

As a control to confirm that the total amount of TGN material was similar in WT and µ1A KO cells, we examined golgin-97. The early Golgi protein GM130 served as a reference marker for quantifying the levels of golgin-97 in the Golgi region. Based on visual inspection and quantification, golgin-97 levels were similar in WT and µ1A KO cells (Figure 3A,D).

**Figure 3.**
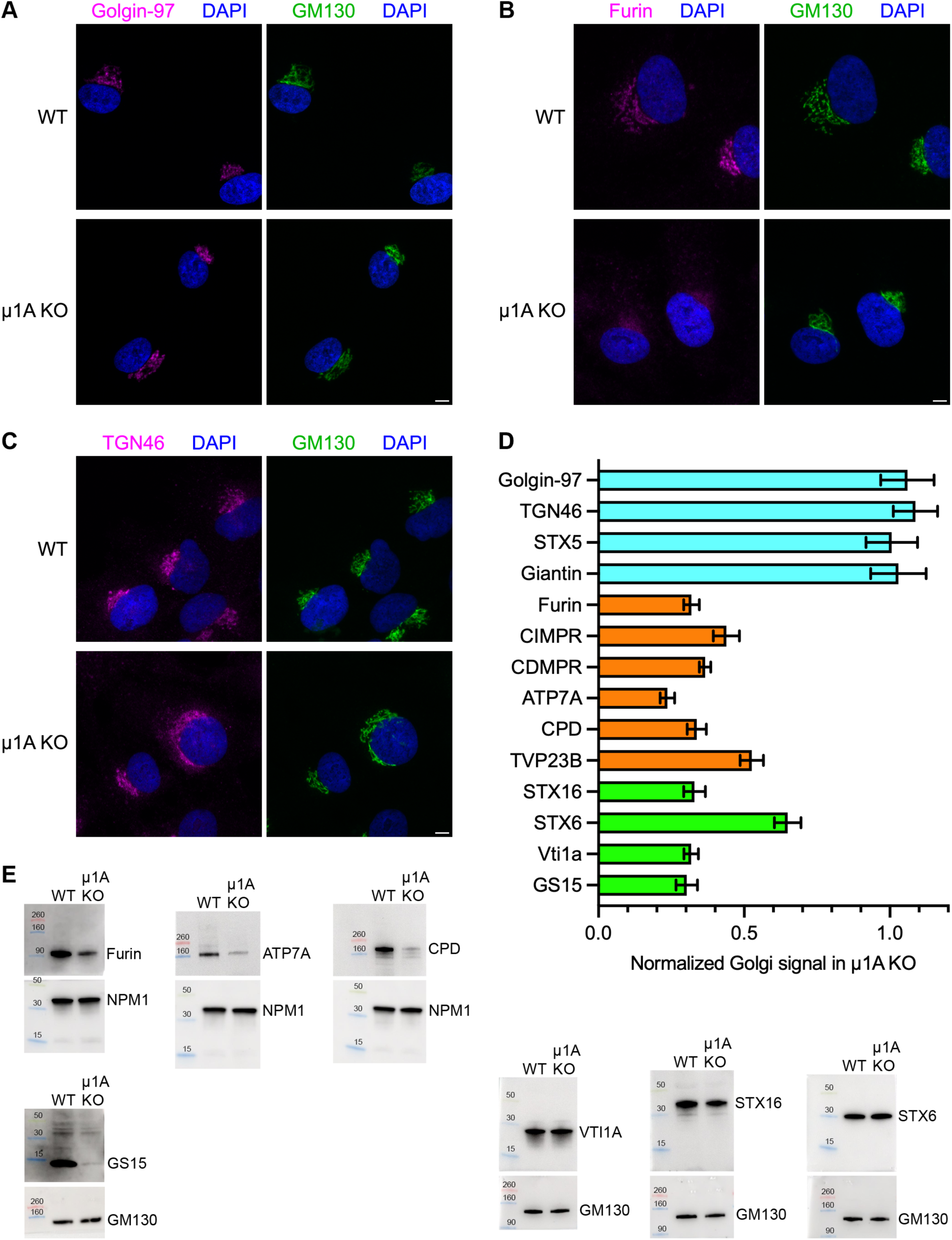
Loss of µ1A in RPE-1 cells results in depletion from the Golgi of multiple resident TGN proteins. **(A-C)** Representative immunofluorescence images of golgin-97, furin, or TGN46 (magenta) together with GM130 (green) in WT and µ1A KO cells. Each comparison shows cells imaged in the same experiment using the same parameters. DAPI-labeled nuclei are shown in blue. Scale bars, 5 µm. **(D)** Ǫuantification of the depletion of various resident proteins from the Golgi in µ1A KO cells. The average signal for each of the indicated proteins in µ1A KO cells was normalized to the corresponding average signal in WT cells. Bars represent SEM. Blue represents Golgi proteins that were unaffected by loss of AP-1, orange represents Golgi non-SNARE proteins that were affected by loss of AP-1, and green represents Golgi SNARE proteins that were affected by loss of AP-1. Each experiment was done 3 times on different days, and the normalization was relative to the WT cell data from the same-day experiment. The total number of cells analyzed was as follows: golgin 97 = 41 WT, 41 µ1A KO; furin = 41 WT, 38 µ1A KO; TGN46 = 36 WT, 42 µ1A KO; CIMPR = 42 WT, 35 µ1A KO; CDMPR = 41 WT, 52 µ1A KO; ATP7A = 39 WT, 31 µ1A KO; CPD = 32 WT, 37 µ1A KO; STX16 = 33 WT, 30 µ1A KO; STX6 = 42 WT, 45 µ1A KO; Vti1a = 35 WT, 36 µ1A KO; TVP23B = 34 WT, 35 µ1A KO; STX5 = 35 WT, 36 µ1A KO; giantin = 34 WT, 36 µ1A KO; GS15 = 33 WT, 31 µ1A KO. **(E)** Immunoblots showing the levels of various resident Golgi proteins in WT and µ1A KO cells. The numbers in the first lanes represent the sizes in kDa of molecular weight marker proteins. Blots at the top show the Golgi proteins. Blots at the bottom show the loading control protein, either NPM1 (nucleophosmin) or GM130.

Mammalian furin is a proprotein convertase that resides in the TGN (Thomas, 2002). Furin is related to yeast Kex2, which recycles in the AP-1/Ent5 pathway (Casler et al., 2022; Glick, 2026; Krahn et al., 2026b). Knockdown of AP-1 reportedly depleted a furin fusion protein from the Golgi (Hirst et al., 2005), and furin has been identified as an AP-1 cargo (Teuchert et al., 1999; Hirst et al., 2012). In accord with those earlier results, we observed depletion of furin from the Golgi in µ1A KO cells (Figure 3B,D). The magnitude of furin depletion was somewhat variable, but on average, the furin signal in the Golgi region of µ1A KO cells was about one-third the signal in the Golgi region of WT cells (Figure 3D). For this quantification, confocal z-stacks were average projected, and the Golgi signal was defined as the fluorescence in the Golgi region minus the background fluorescence from an identically sized area outside the Golgi region. This method may underestimate loss from the Golgi in µ1A KO cells because the juxtanuclear Golgi region of the cell is relatively thick and therefore includes a high background signal. Thus, furin was strongly depleted from the Golgi in µ1A KO cells. All such experiments have the caveat that the effects could be indirect, but a straightforward interpretation is that AP-1 helps to maintain furin in the Golgi.

To evaluate the specificity of this result, we examined a transmembrane TGN protein that was not predicted to show depletion in µ1A KO cells. TGN46 (TGOLN2), also known as TGN38 in rodents, is a secretory cargo receptor that cycles between the TGN and the plasma membrane (Luzio et al., 1990; Banting and Ponnambalam, 1997; Lujan et al., 2024). TGN46 has not been identified as an AP-1 cargo, and indeed, the Golgi signal for TGN46 was similar in WT versus µ1A KO cells (Figure 3C,D).

In a study of µ1A knockout mouse cells, it was reported that CIMPR and CDMPR largely disappeared from the Golgi (Meyer et al., 2000). We found that CIMPR and CDMPR were depleted from the Golgi in µ1A KO cells, although for CIMPR the depletion was not quite as extensive as with furin (Figure S3A,B and Figure 3D). CIMPR and CDMPR have been identified as AP-1 cargoes (Ghosh et al., 2003; Hirst et al., 2012). The combined results support the idea that AP-1 helps to maintain CIMPR and CDMPR in the Golgi.

Two prominent AP-1 cargoes are the copper transporter ATP7A, which resides in the TGN at low copper concentrations, and the processing enzyme carboxypeptidase D (CPD), which also resides in the TGN (Arolas et al., 2007; Hirst et al., 2012; Polishchuk and Lutsenko, 2013). Inactivation of AP-1 was reported to deplete both proteins from the Golgi (Hirst et al., 2012; Bonnemaison et al., 2014). Indeed, we found that ATP7A and CPD levels in the Golgi were strongly reduced in µ1A KO cells (Figure S3C,D and Figure 3D).

In yeast cells, the Ǫa-SNARE Tlg2 and the Ǫc-SNARE Tlg1 recycle together with Kex2 in the AP-1/Ent5 pathway (Casler et al., 2022; A. Pantazopoulou, in preparation). The mammalian orthologs are syntaxin 16 (STX16) and syntaxin 6 (STX6), respectively (Struthers et al., 2009; Pérez-Victoria et al., 2010). In line with previous work (Hirst et al., 2012), both of these mammalian Ǫ-SNAREs showed reduced Golgi localization in µ1A KO cells (Figure S3E,F and Figure 3D). For syntaxin 6, however, the reduction was relatively mild, with significant residual Golgi signal visible in many cells. We also examined Vti1a, which is the Ǫb-SNARE component of the Golgi-localized SNARE complex that contains STX16 and STX6 (Tang et al., 2022). The Golgi signal for Vti1a was strongly reduced in µ1A KO cells (Figure S3G and Figure 3D). All three of the mammalian Ǫ-SNAREs have been identified as AP-1 cargoes (Hirst et al., 2012), and accordingly, they show a mild to strong dependency on AP-1 for their Golgi localization.

We then examined TVP23B, also known as FAM18B. This transmembrane TGN protein is broadly conserved, and it has a poorly understood role in Golgi function (Inadome et al., 2007; Stein et al., 2009; Gendre et al., 2011; Song et al., 2023). TVP23B was identified as a putative AP-1 cargo in a proteomics screen (Hirst et al., 2012) but the effects of inactivating AP-1 on TVP23B have not been reported. We found that TVP23B was depleted from the Golgi in µ1A KO cells, although the depletion was not as extensive as with some of the other TGN proteins (Figure S3H and Figure 3D).

Control experiments confirmed that other Golgi transmembrane proteins retained their normal localizations after loss of AP-1. The yeast Ǫa-SNARE Sed5 recycles through the ER in a COPI-dependent manner (Gao and Banfield, 2020), and the same is likely true for the mammalian ortholog syntaxin 5 (STX5), which is present throughout much of the Golgi stack (Volchuk et al., 2004). The Golgi signal for STX5 was not detectably different in WT versus µ1A KO cells (Figure S3I and Figure 3D). Similarly, the signal for the early Golgi coiled-coil protein giantin (GOLGB1) was not detectably different in wild-type versus µ1A KO cells (Figure S3J and Figure 3D). These data confirm that elimination of AP-1 selectively depletes a subset of transmembrane proteins from the Golgi.

A particularly interesting protein is the Ǫc-SNARE GS15, also known as BET1L (Xu et al., 2002). GS15 is the ortholog of yeast Sft1, which is present in the late Golgi (Wooding and Pelham, 1998; Xu et al., 2002). Because GS15 is concentrated in the medial/*trans* part of the mammalian Golgi (Xu et al., 2002; Volchuk et al., 2004), it might recycle with the aid of AP-1. GS15 has not previously been identified as a component of AP-1 vesicles, but its small size (∼12 kDa) could have hindered detection by proteomics. We found that GS15 was strongly depleted from the Golgi in µ1A KO cells (Figure S3K and Figure 3D). GS15 is therefore a candidate for being a novel AP-1 cargo.

We wondered whether depletion from the Golgi in µ1A KO cells was accompanied by reduced protein levels. In the case of furin, ATP7A, CPD, and GS15, the protein levels were clearly reduced in µ1A KO cells (Figure 3E). However, for Vti1a, STX16, and STX6, little or no reduction was seen (Figure 3E). A plausible interpretation is that loss of µ1A compromises the ability of certain proteins to remain in the Golgi, and that some but not all of those mislocalized proteins undergo accelerated turnover.

### Loss of TBC1D23 does not mimic loss of AP-1 in RPE-1 cells

The WDR11 complex was implicated in the capture of AP-1 vesicles (Navarro Negredo et al., 2018; Deng et al., 2024), and TBC1D23 was identified as a protein linking the WDR11 complex to golgin tethers that capture vesicles at the TGN (Shin et al., 2017). It was reported that TBC1D23 binds the cytosolic tails of CPD and STX16, and that loss of TBC1D23 depletes CPD from the Golgi of insulinoma INS-1 cells (Cattin-Ortolá et al., 2024). We therefore wondered whether TBC1D23 functions in the AP-1-dependent localization of resident TGN proteins, in which case a TBC1D23 knockout might deplete those proteins from the Golgi. Our RPE-1-derived TBC1D23 knockout cell line (Figure S1A) is here designated "TBC1D23 KO". Unlike µ1A KO cells, TBC1D23 KO cells showed an apparently normal Golgi signal for CPD (Figure 4A,D) and a normal or even slightly elevated signal for furin (Figure 4B,D). We infer that TBC1D23 is not crucial for the AP-1-dependent localization of resident TGN proteins, at least in RPE-1 cells.

**Figure 4.**
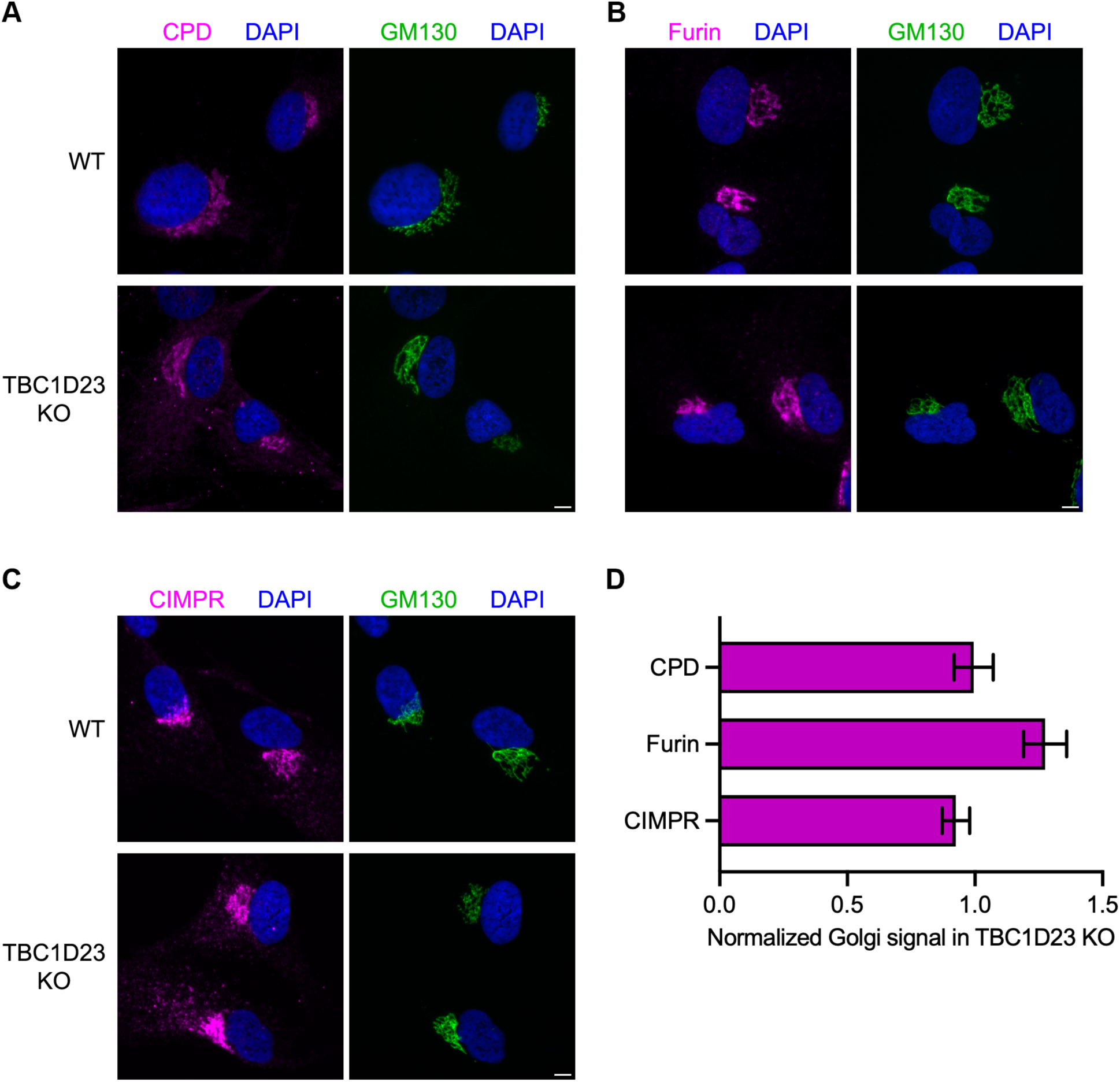
Loss of TBC1D23 does not mimic loss of µ1A. **(A)** Representative immunofluorescence images of CPD (magenta) together with GM130 (green) in WT and TBC1D23 KO cells. Each comparison shows cells imaged in the same experiment using the same parameters. DAPI-labeled nuclei are shown in blue. Scale bars, 5 µm. **(B)** Same as (A) except with furin. **(C)** Same as (A) except with CIMPR. **(D)** Ǫuantification of the Golgi signals for CPD, furin, and CIMPR in WT versus TBC1D23 KO cells. The average signal for each of the indicated proteins in TBC1D23 KO cells was normalized to the corresponding average signal in WT cells. Bars represent SEM. Each experiment was done 3 times on different days, and the normalization was relative to the WT cell data from the same-day experiment. The total number of cells analyzed was as follows: CPD = 32 WT, 36 TBC1D23 KO; furin = 33 WT, 39 TBC1D23 KO; CIMPR = 38 WT, 39 TBC1D23 KO.

Loss of TBC1D23 was also reported to disrupt CIMPR trafficking (Shin et al., 2017). Compared to WT RPE-1 cells, TBC1D23 KO cells showed no significant change in the average CIMPR signal in the Golgi (Figure 4D), but they sometimes showed a more prominent CIMPR signal in peripheral punctate structures that were presumably endosomes (Figure 4C). CIMPR has a complex itinerary that includes frequent traffic to and from endosomes (Ghosh et al., 2003). Thus, a possible interpretation of our findings is that TBC1D23 functions mainly in AP-1-independent recycling from endosomes.

### epsinR assists mammalian AP-1 in maintaining the Golgi localization of several TGN proteins

Based on the close relationship between yeast AP-1 and the epsinR homolog Ent5 (Myers and Payne, 2013; Casler et al., 2022; Krahn et al., 2026b), together with the known physical interaction between mammalian AP-1 and epsinR (Hirst et al., 2003; Mills et al., 2003), we speculated that epsinR might work together with mammalian AP-1 to maintain certain proteins in the TGN. It was reported that epsinR colocalizes with AP-1 (Hirst et al., 2003). Our immunofluorescence analysis confirmed that epsinR shows striking colocalization with the γ1 subunit of AP-1, both at the Golgi and at peripheral recycling endosomes, consistent with a possible functional interaction between epsinR and AP-1 (Figure S3L).

It was previously reported that acute inactivation of epsinR caused depletion from clathrin-coated vesicles of a number of AP-1 cargoes including CIMPR, CDMPR, CPD, STX16, and Vti1a (Hirst et al., 2015), further suggesting that epsinR works together with AP-1. We therefore tested whether loss of epsinR would deplete the Golgi of AP-1 cargoes. This experiment employed an RPE-1-derived cell line in which epsinR had been knocked out (Figure S1A), here designated "epsinR KO". Figure 5A-D shows that loss of epsinR caused strong depletion from the Golgi of CDMPR, CPD, and Vti1a. As controls, there was no depletion of TGN46 or STX5 (Figure 5D). Loss of epsinR caused no detectable depletion from the Golgi of furin, CIMPR, ATP7A, STX16, or STX6 (Figure 5D). These results are partly consistent with the results from acute inactivation of epsinR (Hirst et al., 2015), although for unknown reasons, two of the proteins that were affected in that earlier study (CIMPR and STX16) did not show depletion from the Golgi in our epsinR KO cells. The combined data suggest that epsinR activity is more important for some AP-1 cargoes than for others.

**Figure 5.**
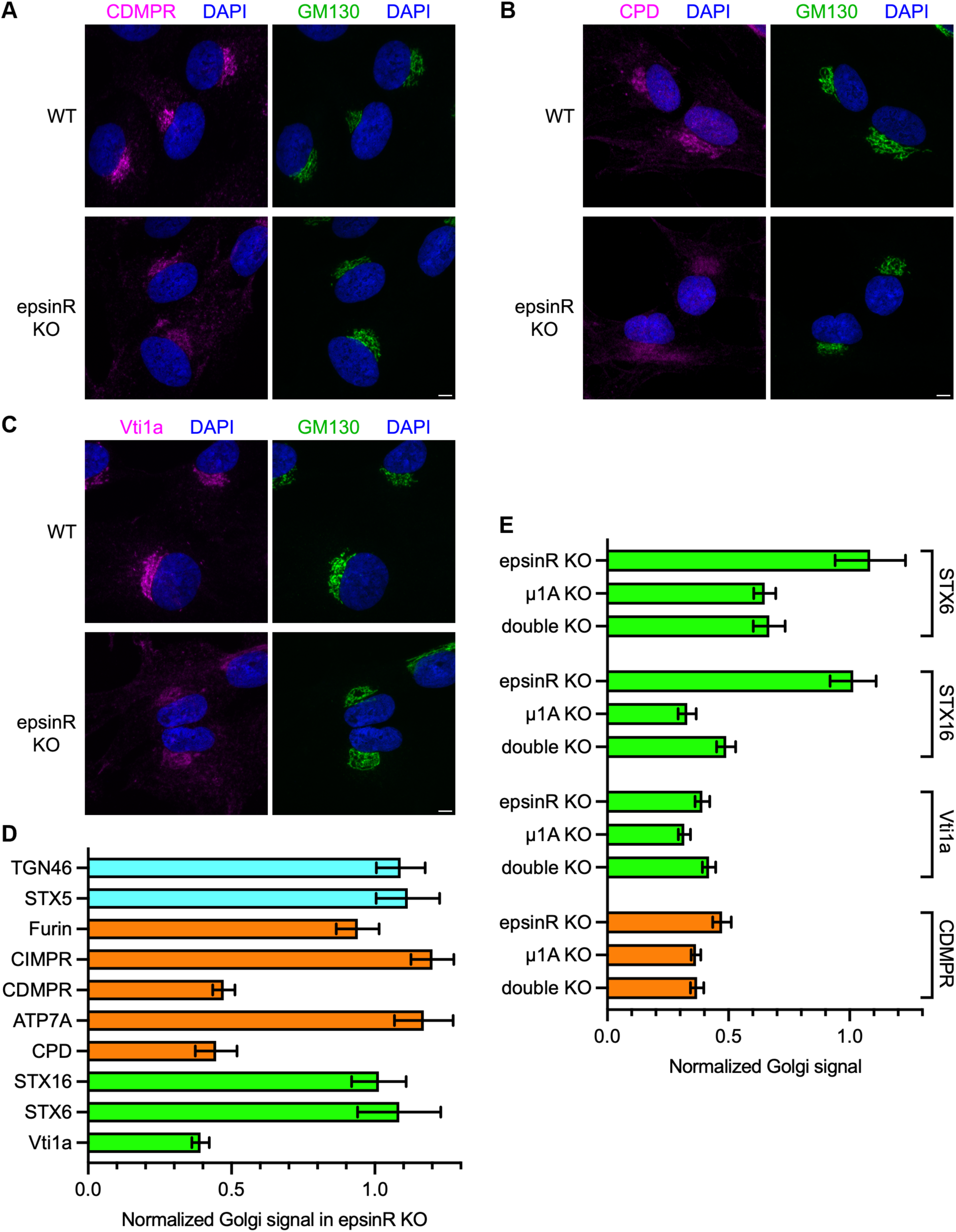
Loss of epsinR in RPE-1 cells results in depletion from the Golgi of a limited set of resident TGN proteins. **(A-C)** Representative immunofluorescence images of CDMPR, CPD, or Vti1a (magenta) together with GM130 (green) in WT and epsinR KO cells. Each comparison shows cells imaged in the same experiment using the same parameters. DAPI-labeled nuclei are shown in blue. Scale bars, 5 µm. **(D)** Ǫuantification of the depletion of various resident proteins from the Golgi in epsinR KO cells. The average signal for each of the indicated proteins in epsinR KO cells was normalized to the corresponding average signal in WT cells. Bars represent SEM. Blue represents control Golgi proteins, orange represents Golgi non-SNARE proteins, and green represents Golgi SNARE proteins. Each experiment was done 3 times on different days, and the normalization was relative to the WT cell data from the same-day experiment. The total number of cells analyzed was as follows: TGN46 = 40 WT, 36 epsinR KO; STX5 = 38 WT, 36 epsinR KO; Furin = 36 WT, 35 epsinR KO; CIMPR = 40 WT, 38 epsinR KO; CDMPR = 48 WT, 53 epsinR KO; ATP7A = 35 WT, 35 epsinR KO; CPD = 39 WT, 42 epsinR KO; STX16 = 35 WT, 37 epsinR KO; STX6 = 39 WT, 35 epsinR KO; VTI1A = 31 WT, 37 epsinR KO. **(E)** Ǫuantification of the depletion of four resident proteins from the Golgi in epsinR KO cells, µ1A KO cells, and epsinR/µ1A double KO cells. The average signal for each of the indicated proteins in KO cells was normalized to the corresponding average signal in WT cells. Bars represent SEM. Green represents Golgi SNARE proteins, and orange represents a Golgi non-SNARE protein. Each experiment was done 3 times on different days, and the normalization was relative to the WT cell data from the same-day experiment. The µ1A KO data are from the experiment of Figure 3, and the epsinR KO data are from the experiment of part (D). For the double KO analysis, the total number of cells analyzed was as follows: STX6 = 47 WT, 35 double KO; STX16 = 39 WT, 37 double KO; Vti1a = 34 WT, 40 double KO; CDMPR = 39 WT, 37 double KO.

Because loss of yeast Ent5 perturbs Golgi localization of the SNAREs Tlg2 and Tlg1 (Hung and Duncan, 2016; Casler et al., 2022; Petersen et al., 2023), we were surprised that loss of epsinR had no effect on Golgi localization of the homologous mammalian SNAREs STX16 and STX6. It seemed possible that AP-1 normally acts to maintain STX16 and STX6 in the Golgi and that epsinR serves as a backup, in which case the contribution from epsinR would only become evident after loss of AP-1. A sensitive test of this hypothesis focused on STX6, which showed only mild depletion from the Golgi in µ1A KO cells (see Figure 3), leading us to predict that additional loss of epsinR might enhance this depletion. We therefore examined STX6 in double-knockout cells lacking both µ1A and epsinR (Figure 5E). Contrary to our prediction, the results from the double-knockout cells resembled the results from the µ1A KO cells for STX6. The same was true for STX16. Moreover, even though epsinR is needed for normal localization of Vti1a and CDMPR, those proteins did not show stronger depletion from the Golgi in the double-knockout cells than in the µ1A KO cells (Figure 5E). Thus, epsinR does not partially compensate for the loss of AP-1, at least for localizing the TGN proteins examined here. Taken together, the data suggest that epsinR assists AP-1 in maintaining the Golgi localization of a subset of TGN proteins but that epsinR probably cannot perform this function independently of AP-1.

### Structure predictions suggest that AP-1 forms similar complexes with mammalian epsinR and yeast Ent5

Based on the indications that both mammalian epsinR and yeast Ent5 interact with AP-1 and work together with AP-1, we speculated that epsinR and Ent5 might show a conserved physical association with AP-1. This idea was explored by generating structure predictions with AlphaFold (Abramson et al., 2024). For epsinR and human AP-1, we included GTP-bound ARF1 in the modeling because two copies of ARF1 were shown to bind to the AP-1 tetramer to stabilize the cargo-binding conformation (Ren et al., 2013). A recent study reported an interaction between ARF1 and epsinR (Uehara et al., 2026), and indeed, AlphaFold generated a high-confidence prediction of an interaction between ARF1 and the N-terminal ENTH domain of epsinR (Figure 6A). Because the predicted interaction interface did not overlap with the established ARF1:AP-1 or epsinR:Vti1b interactions (Miller et al., 2007; Ren et al., 2013), we modeled a complex in which ARF1 is bound to both AP-1 and epsinR while epsinR is bound to the H_abc_ domain of Vti1b (Figure 6B). epsinR has been described as a potential PI4P-binding protein based on *in vitro* data (Hirst et al., 2003; Mills et al., 2003), but PI4P was not included in the model because this ligand did not improve the prediction. The organization of the predicted complex is compatible with membrane association, as the N-terminal amphipathic helices of ARF1 and epsinR lie on the same surface where AP-1 engages with cargo and lipids.

**Figure 6.**
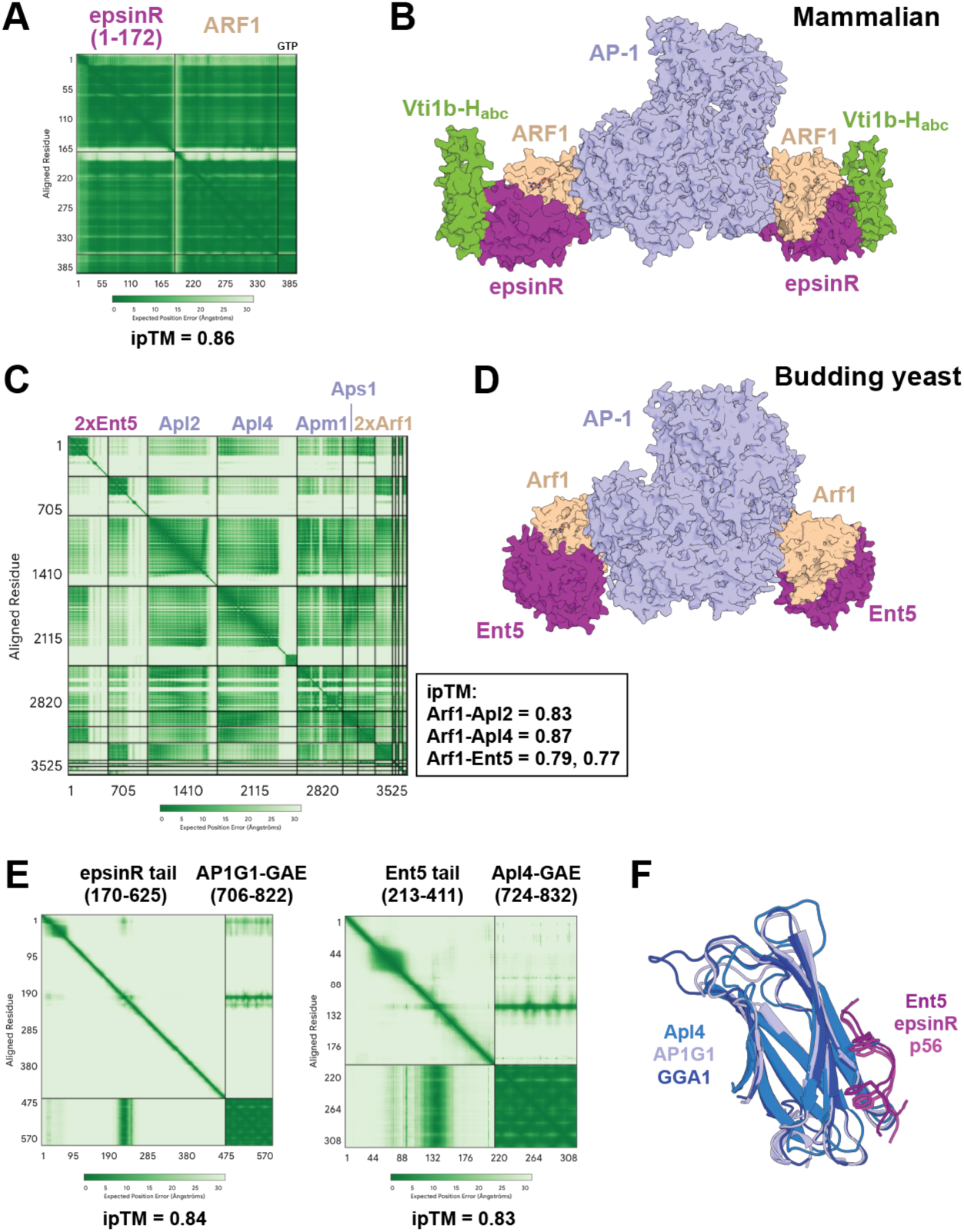
Structural models of the AP-1:ARF1:epsinR and AP-1:Arf1:Ent5 complexes. **(A)** Predicted aligned error (PAE) plot for an AlphaFold3 prediction (Abramson et al., 2024) of the N-terminal domain of human epsinR and human ARF1-GTP. **(B)** Structural model of a mammalian AP-1:ARF1-GTP:epsinR:Vti1b complex. This model was generated by superposition of the cryo-EM structure of mammalian AP-1:ARF1-GTP (PDB: 6DFF; Morris et al., 2018), the crystal structure of epsinR:Vti1b (PDB: 2V8S; Miller et al., 2007), and the epsinR:ARF1-GTP prediction. The N-terminal amphipathic helices of ARF1 and epsinR are not visible, but both lie at the bottom of the model, consistent with membrane engagement. **(C)** PAE plot, and interface predicted template modeling (ipTM) scores for selected chain-chain interactions, for an AlphaFold3 prediction of a yeast AP-1:Arf1-GTP:Ent5:PI(4)P complex. A short-chain version of PI(4)P with 8 carbons in each acyl chain was used. **(D)** Structural model of the predicted AP-1:Arf1-GTP:Ent5:PI(4)P complex with the flexible regions and gamma-adaptin ear (GAE) domains omitted. **(E)** PAE plots of AlphaFold3 predictions of interactions between the GAE domain of human gamma-adaptin (AP1G1) and the epsinR C-terminal tail (***left***) and between the GAE domain of yeast gamma-adaptin (Apl4) and the Ent5 C-terminal tail (***right***). The ipTM scores indicate high-confidence predictions. **(F)** Superposition of the GGA1-GAE:p56 crystal structure (PDB: 1OM9; Collins et al., 2003) with the AP1G1-GAE:epsinR-tail prediction and the Apl4-GAE:Ent5-tail prediction. This analysis suggests similar interactions of epsinR and Ent5 with the GAE domains of AP1G1 and Apl4, respectively. The predicted interactions involve sequences corresponding to GAE interacting motifs (371-DFGDW-375 of epsinR and 330-EFGDF-334 of Ent5). For clarity, only the hydrophobic side chains of the motifs are shown.

We then used AlphaFold to determine whether a similar complex of the homologous yeast proteins is also plausible. The ENTH domain of Ent5 has a PI4P binding pocket (Zhang et al., 2016) and *in vivo* data implicated PI4P in the membrane recruitment of Ent5 (Daboussi et al., 2012), so in this case PI4P was included in the model. The prediction was high-confidence, and it yielded a complex similar to the one obtained with the mammalian proteins (Figure 6C,D).

The flexible C-terminal tails of both epsinR and Ent5 have been reported to interact with the gamma-adaptin ear (GAE) domain (Duncan et al., 2003; Hirst et al., 2003; Mills et al., 2003). Surprisingly, this interaction was not confidently predicted when we modeled the entire human complex. Better results were obtained by using only the tail of epsinR or Ent5 together with the corresponding GAE domain of gamma-adaptin. High confidence predictions (Figure 6E) showed epsinR or Ent5 sequences matching the expected fi-G-[P/D/E]-[fi/L/M] GAE-binding motif (Mattera et al., 2004) bound to the same GAE domain surface of human gamma-adaptin (AP1G1) or yeast gamma-adaptin (Apl4) that was previously reported for the GGA1 GAE domain (Collins et al., 2003) (Figure 6F). Thus, structural predictions suggest that the epsinR:ARF1:AP-1 complex and the Ent5:Arf1:AP-1 complex are similar, and that associations mediated by the GTPase are likely augmented by interactions between the adaptors.

### The yeast CPD-like protein Kex1 is a TGN resident that recycles in the AP-1/Ent5 pathway

The observation that AP-1 plays a conserved role in the localization of certain classes of TGN proteins made us wonder whether we could predict novel yeast AP-1 cargoes by their similarity to mammalian AP-1 cargoes. Mammalian CPD is a carboxypeptidase that acts after furin during proprotein processing, and similarly, yeast Kex1 is a carboxypeptidase that acts after Kex2 during proprotein processing (Fuller et al., 1988; Cooper and Bussey, 1989). Although Kex1 is not evolutionarily related to CPD, the two proteins have the same type I transmembrane topology and the same kind of enzymatic activity (Cooper and Bussey, 1992). Kex1 is reportedly present in the TGN (Bryant and Boyd, 1993; Lussier et al., 1995) but its trafficking had not been characterized. By analogy to CPD, we predicted that Kex1 trafficking would involve AP-1, in which case Kex1 would behave similarly to Kex2, which is a TGN protein that recycles in the AP-1/Ent5 pathway (Casler et al., 2022; Glick, 2026; Krahn et al., 2026b).

To test this idea, gene replacement was used to tag Kex1 at its C-terminus with two copies of a monomeric superfolder GFP (Valbuena et al., 2020). Fluorescence microscopy revealed puncta that varied in size and intensity (Figure S4A and Video 5). Such puncta represent Golgi cisternae, which are optically resolvable because the *S. cerevisiae* Golgi is not stacked (Wooding and Pelham, 1998; Papanikou and Glick, 2009).

As a first step in characterizing the trafficking of Kex1, we performed kinetic mapping by 4D confocal microscopy (Losev et al., 2006; Johnson and Glick, 2019). A strain was engineered to express GFP-tagged Kex1 together with HaloTag-modified Vrg4, which is an early Golgi protein that recycles in a COPI-dependent pathway (Abe et al., 2004; Losev et al., 2006; Papanikou et al., 2015). HaloTag was then labeled with the far-red dye JFX_650_ (Grimm et al., 2021). The arrival and departure of each tagged protein was quantified for individual cisternae, and the resulting fluorescence traces were smoothed, aligned, and combined to generate averaged traces (Figure S4A, Figure 7A, and Video 5) (Casler et al., 2022). Kex1 began to arrive as Vrg4 was departing, as expected if Kex1 recycles within the TGN. Imaging was then performed with a strain engineered to express GFP-tagged Kex1 together with HaloTag-modified Kex2. Kex1 arrived and departed synchronously with Kex2 (Figure S4A, Figure 7A, and Video 5), suggesting that Kex1 and Kex2 recycle in the same vesicle population.

To test directly whether Kex1 and Kex2 travel in the same vesicles, we employed a vesicle capture assay (Krahn et al., 2026b). A bud-neck-localized septin was tagged with a red fluorescent protein and with FK506-binding protein (FKBP), and Kex2 was tagged at its cytosolic C-terminus with the FKBP-rapamycin-binding domain (FRB). Treatment with rapamycin for 5 min caused Kex2-containing vesicles to be captured at the bud neck (Krahn et al., 2026b). In the same strain, Kex1 had been tagged with GFP, and we asked whether some of the fluorescent Kex1 molecules accumulated at the bud neck after addition of rapamycin. Indeed, an average of 3.4% of the Kex1 signal was detected at the bud neck in the presence of rapamycin versus only 0.3% in the absence of rapamycin (Figure 7B). As controls, Figure S4B is a reproduction of previous results (Krahn et al., 2026b) showing that the AP-1/Ent5 cargo Stv1 was robustly co-captured with Kex2 while the COPI cargo Vrg4 was not detectably co-captured. These data indicate that Kex1 localizes to the TGN by recycling together with Kex2 in AP-1/Ent5 vesicles.

We previously found that when AP-1 and Ent5 are deleted, resident TGN proteins that follow the AP-1/Ent5 pathway escape to the plasma membrane and then return to the TGN by endocytosis (Day et al., 2018; Casler et al., 2022). This phenomenon can be observed by knocking out the AP-1 subunit Apl4 together with Ent5 (Casler et al., 2022). In *apl4Δ ent5Δ* cells, Kex2 was still found in the Golgi but the signal was reduced, presumably because a fraction of the Kex2 molecules were now at the plasma membrane or traveling between the Golgi and the plasma membrane (Casler et al., 2022) (Figure 7C). For Kex1, the Golgi signal in *apl4Δ ent5Δ* cells was similarly reduced (Figure 7C). A more rigorous test involves blocking endocytosis in the mutant cells with the Arp2/3 inhibitor CK-666 (Hetrick et al., 2013; Casler et al., 2022; Krahn et al., 2026b). CK-666 treatment of *apl4Δ ent5Δ* cells caused Kex2 to accumulate in secretory vesicles at sites of polarized growth (Casler et al., 2022; Krahn et al., 2026b) (Figure 7C). As a negative control, the COPI cargo Vrg4 showed no such accumulation (Casler et al., 2022; Krahn et al., 2026b) (Figure 7C). Like Kex2, Kex1 accumulated at sites of polarized growth in CK-666-treated *apl4Δ ent5Δ* cells (Figure 7C). This result confirms that in wild-type cells, the intra-Golgi recycling of Kex1 employs the AP-1/Ent5 pathway. Thus, the trafficking of yeast Kex1 is similar to that of mammalian CPD.

### Mammalian AP-1 keeps proteins in the TGN by preventing their escape to the plasma membrane

The similarities between yeast and mammalian AP-1 cargoes, together with the documented role of yeast AP-1 (Casler et al., 2022; Glick, 2026; Krahn et al., 2026b), led to a hypothesis about the role of mammalian AP-1 at the Golgi. A plausible idea is that mammalian AP-1 recycles resident TGN proteins from maturing TGN compartments to young TGN compartments, thereby keeping those proteins from escaping to the plasma membrane. If so, then inhibition of AP-1-dependent recycling should enable TGN proteins to enter secretory carriers destined for the plasma membrane. We tested this hypothesis as follows.

As a model TGN protein, we fused luminal mScarlet to the transmembrane domain and cytosolic tail of furin (Figure 8A). The furin cytosolic tail contains TGN localization information in the form of two AP-1 interaction signals (Schäfer et al., 1995; Teuchert et al., 1999; Thomas, 2002; Navarro Negredo et al., 2017). Therefore, in addition to the wild-type furin construct, we created a mutant construct in which key residues in the AP-1 interaction signals were replaced by alanines (Tie et al., 2016) (Figure 8A). Each construct also contained an N-terminal (luminal) SBP tag for use in a RUSH protocol (Figure 8A). At 12-15 min after biotin addition, a large fraction of the furin fusion protein molecules had reached the Golgi, and we took advantage of the efficient photobleaching of mScarlet to perform iFRAP. The pool of unbleached Golgi-localized molecules was tracked by live-cell imaging. Our prediction was that the wild-type furin construct would remain in the Golgi, whereas the mutant furin construct would be unable to bind AP-1 and would therefore rapidly escape to the plasma membrane. If the cells were mutated to inactivate AP-1, then the wild-type and mutant furin constructs should escape to the plasma membrane at similar rates.

**Figure 7.**
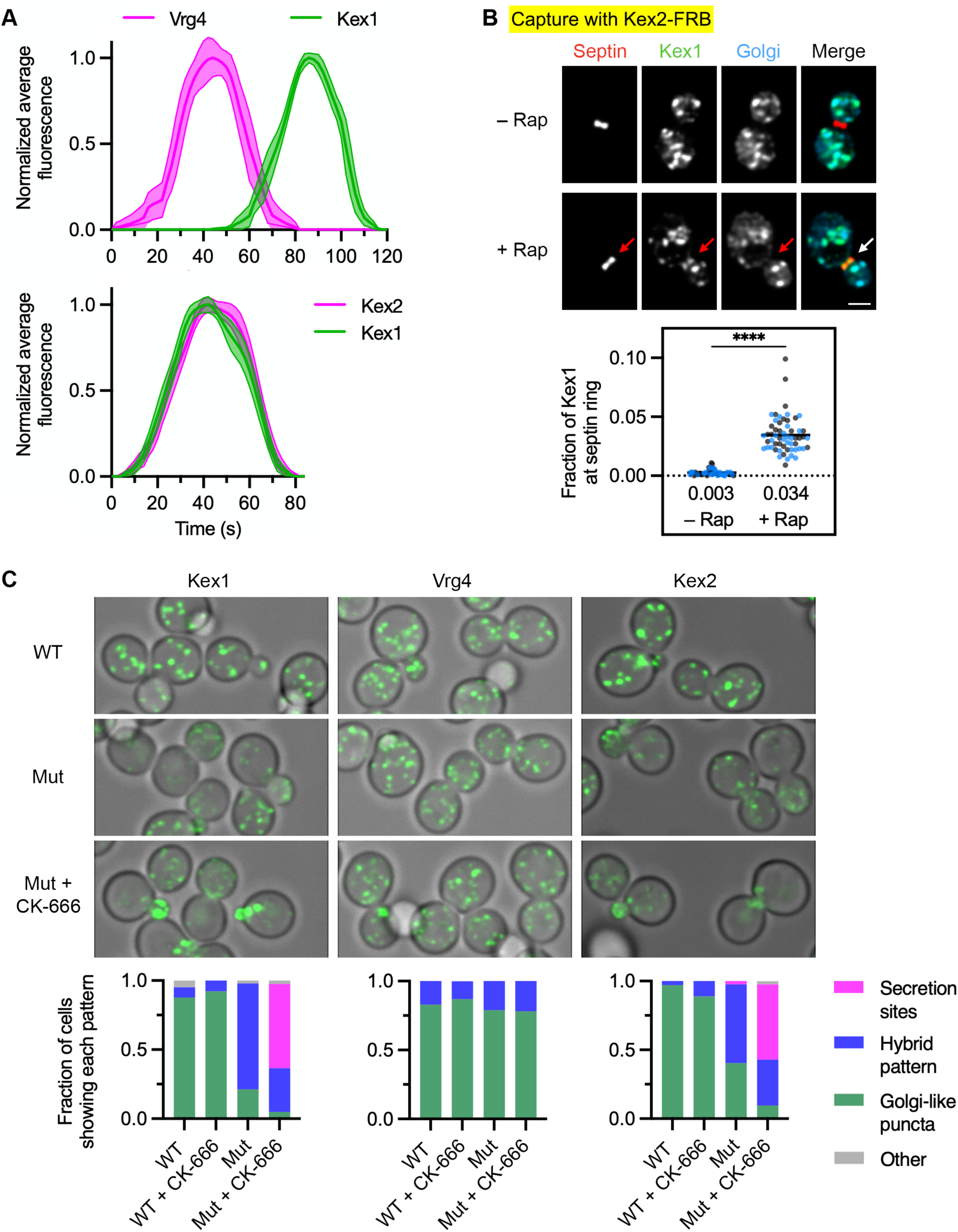
The CPD-like yeast protein Kex1 localizes to the late Golgi by recycling together with Kex2 in the AP-1/Ent5 pathway. **(A)** Golgi maturation kinetics of Kex1 relative to Vrg4 and Kex2. Green represents a GFP label and magenta represents a HaloTag label. Shown are normalized and averaged fluorescence traces derived from 10 individual traces for the Vrg4-Kex1 comparison or 16 individual traces for the Kex2-Kex1 comparison. The plots show solid lines for mean values plus shaded areas for 95% confidence intervals. **(B)** Rapamycin-dependent co-capture of Kex1 with Kex2-FRB by an FKBP-tagged septin. In the fluorescence micrographs, the Shs1 septin was tagged with FKBP plus mScarlet (red), Kex1 was tagged with GFP (green), and Golgi cisternae were marked by HaloTag-labeled Ric1 and Sec7 (blue). The arrows show a particularly clear example of non-Golgi Kex1 signal at the bud neck in cells treated for 5 min with rapamycin (“Rap”). In the graph, co-capture of GFP-tagged Kex1 with Kex2-FRB was quantified for multiple cells by measuring the fraction of the total Kex1 signal within a septin mask, which had been modified by subtraction of a Golgi mask. The black and blue dots represent two biological replicates. The total numbers of cells examined were 51 ("– Rap") and 58 ("+ Rap"). Mean capture values for the combined individual data points from the two biological replicates are displayed as thick horizontal bars and are listed numerically below the plots. ****, significant at P value <0.0001. **(C)** Perturbation of Kex1 trafficking by loss of AP-1 and Ent5. The images and plots show that in mutant cells lacking AP-1 and Ent5, inhibition of endocytosis with CK-666 affected the localizations of Kex1 and of the AP-1/Ent5 cargo Kex2 but not of the COPI cargo Vrg4. Wild-type “WT” or *apl4Δ ent5Δ* mutant (“Mut”) cells expressing the indicated GFP-tagged Golgi proteins were examined by fluorescence microscopy after either mock treatment or treatment for 15-25 min with CK-666. The fluorescence pattern of each tagged protein was quantified by assigning cells to categories based on whether most of the protein was in Golgi-like puncta, or whether most of the protein was concentrated at secretion sites, or whether the protein was distributed in a hybrid pattern between Golgi-like puncta and secretion sites. “Other” indicates that a cell could not be confidently assigned to any of the three categories. Approximately 30-45 cells were examined for each Golgi protein under each condition. Representative images are shown for mock-treated wild-type cells, mock-treated mutant cells, and CK-666-treated mutant cells. For a given GFP-tagged protein, the images were all captured and processed with the same parameters.

**Figure 8.**
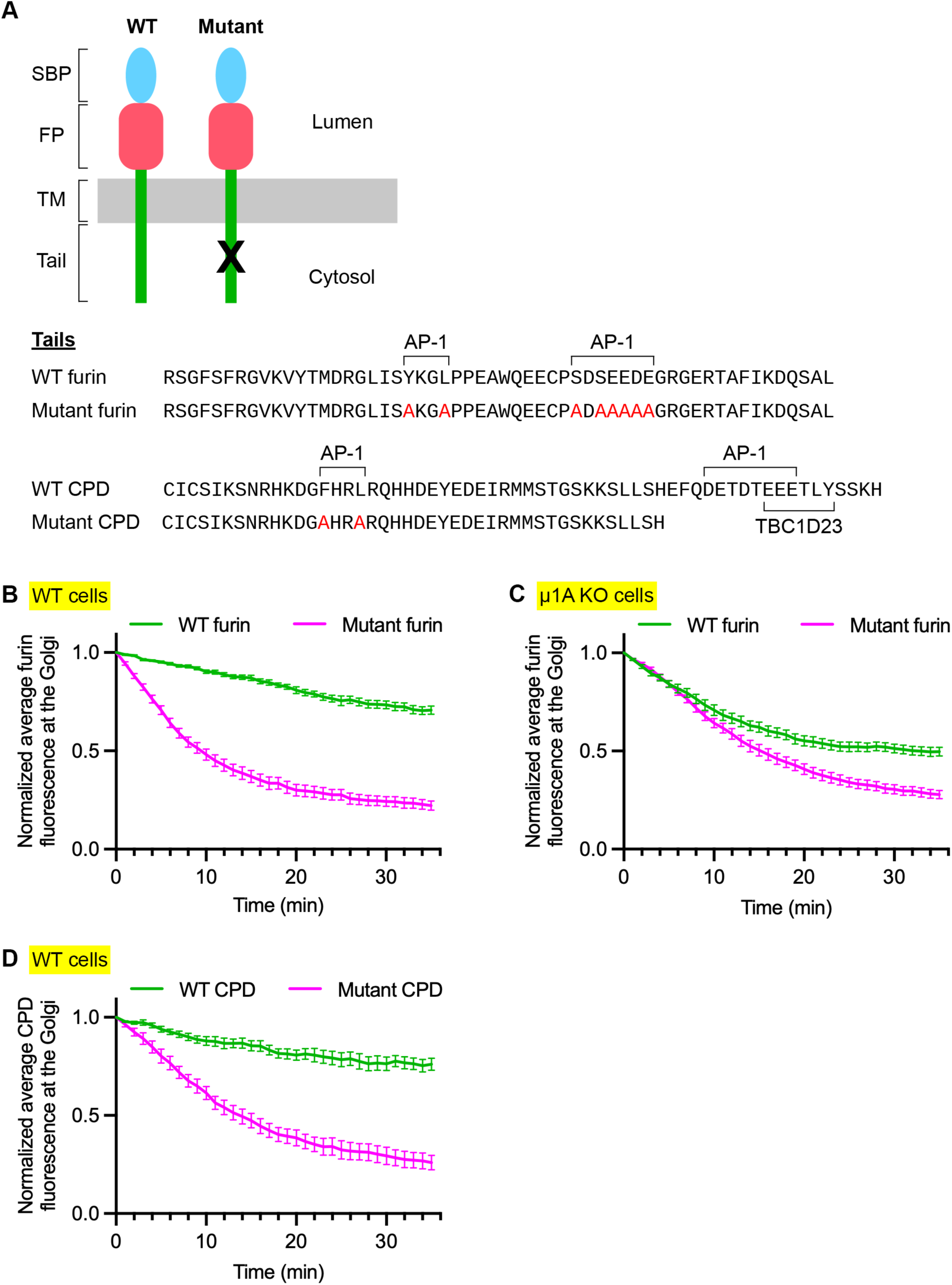
Loss of AP-1 binding allows resident TGN proteins to escape the Golgi. **(A) *Top:*** Schematic diagram of the constructs used here and in Figure 9. SBP, streptavidin-binding peptide; FP, fluorescent protein; TM, transmembrane domain; Tail, cytosolic tail. Green represents the sequences from the original furin and CPD proteins, including a short luminal sequence (6 amino acids for furin or 9 amino acids for CPD). The black "X" represents mutations in the cytosolic tails of the mutant constructs. ***Bottom:*** Sequences of the cytosolic tails of the wild-type (WT) and mutant constructs, with the AP-1 and TBC1D23 recognition sites indicated for the WT constructs. Amino acids that were changed to alanine in the mutant constructs are shown in red. **(B)** Rapid Golgi export in WT RPE-1 cells of a mutant furin construct that cannot be recognized by AP-1. To eliminate AP-1 binding, the cytosolic tail was mutated to replace YKGL with AKGA and to replace SDSEEDE with ADAAAAA. Cells were transiently transfected to express GalNAc-T2-HaloTag plus either an SBP-mScarlet-tagged wild-type (WT) furin construct or an SBP-mScarlet-tagged mutant furin construct lacking the AP-1 recognition signals. GalNAc-T2-HaloTag was labeled with JFX_650_ to mark the Golgi. The WT or mutant construct was released from the ER by addition of biotin, and the cells were incubated for 10-15 min to accumulate the construct in the Golgi. Then non-Golgi red fluorescence was photobleached, and export of the unbleached furin construct from the Golgi was tracked by measuring the remaining red fluorescence that overlapped with the Golgi. Post-photobleaching quantification of Golgi export began ∼20 min after biotin addition. This analysis was performed for 14 cells expressing the WT construct (green) or 13 cells expressing the mutant construct (magenta) from a total of 4 separate dishes (WT) or 5 separate dishes (mutant) on 2 different days. The plots for the individual cells were normalized and then averaged. Shown are mean and SEM values at each time point. Representative movies for the two constructs are shown in Video 6. **(C)** Rapid Golgi export in µ1A KO cells of both the wild-type and mutant furin constructs. The experiment was performed as in (A) except with µ1A KO cells. Post-photobleaching quantification of Golgi export began ∼26 min after biotin addition. The different timing relative to WT cells reflected the slower kinetics of Golgi export in µ1A KO cells. This analysis was performed for 20 cells expressing the WT construct (green) or 27 cells expressing the mutant construct (magenta) from a total of 6 separate dishes (WT) or 8 separate dishes (mutant) on 3 different days. Representative movies for the two constructs are shown in Video 7. **(D)** Rapid Golgi export in WT RPE-1 cells of a mutant CPD construct that cannot be recognized by AP-1. To eliminate AP-1 binding, the cytosolic tail was mutated to replace FHRL with AHRA and to remove the last 18 amino acids. The experiment was performed as in (A) except using either an SBP-mScarlet-tagged wild-type (WT) CPD construct or an SBP-mScarlet-tagged mutant CPD construct lacking the AP-1 recognition signals. Post-photobleaching quantification of Golgi export began ∼20 min after biotin addition. This analysis was performed for 13 cells expressing the WT construct (green) or 9 cells expressing the mutant construct (magenta) from a total of 4 separate dishes (WT) or 3 separate dishes (mutant) on 2 different days. Representative movies for the two constructs are shown in Video 8.

The results of this experiment are shown in Figure 8 and the accompanying videos. In WT RPE-1 cells, the wild-type furin construct largely remained in the Golgi (Figure 8B and Video 6). A slow, partial reduction in the Golgi signal was seen, perhaps because expression of this construct pushed the Golgi retention system beyond its capacity. By comparison, the mutant furin construct departed rapidly from the Golgi and was mostly depleted from the Golgi after 30 min (Figure 8B and Video 6). In µ1A KO RPE-1 cells, the wild-type fusion construct initially departed from the Golgi at the same rate as the mutant furin construct, as expected (Figure 8C and Video 7). This initial departure rate was somewhat slower than the initial departure rate of the mutant furin construct in WT cells (Figure 8B,C), consistent with the finding that Golgi export is slowed in µ1A KO cells (see Figure 1). By the end of the time course, the average level of the mutant furin construct in the Golgi was about the same in µ1A KO cells as in WT cells (Figure 8B,C). However, in µ1A KO cells, the final level of the wild-type furin construct was higher on average than the final level of the mutant furin construct (Figure 8C). The wild-type furin construct contained endocytosis signals that overlapped with the AP-1 interaction signals (Schäfer et al., 1995; Voorhees et al., 1995; Teuchert et al., 1999), so after the wild-type furin construct escaped to the plasma membrane, it was probably internalized once again to the Golgi. The key conclusion here is that Golgi retention of a furin fusion construct was compromised either by disrupting the AP-1 binding sites in the construct or by inactivating AP-1. With both treatments, the furin fusion construct exited the Golgi with kinetics reminiscent of a secretory cargo. These results favor the idea that AP-1 acts to keep furin in the Golgi.

To extend the analysis beyond furin, we created a RUSH construct in which a luminal SBP-mScarlet cassette was fused to the transmembrane domain and cytosolic tail of CPD (Figure 8A). We then created a mutant construct that lacked the last 18 amino acids of the CPD cytosolic tail, thereby removing binding sites for AP-1 and TBC1D23 (Eng et al., 1999; Navarro Negredo et al., 2017; Cattin-Ortolá et al., 2024) (Figure 8A). The mutant construct also had an additional change: an upstream FHRL tetrapeptide that is a likely binding site for clathrin adaptors (Kalinina et al., 2002; Navarro Negredo et al., 2017) was mutated to AHRA (Figure 8A). After release from the ER, wild-type mScarlet-CPD accumulated in the Golgi and then largely remained there, while mutant mScarlet-CPD accumulated in the Golgi and then departed rapidly (Figure 8D and Video 8). Thus, two different TGN proteins fail to maintain their normal Golgi localization when they cannot bind to AP-1.

A simple way to understand these results is that when cytosolic tails are mutated to remove the AP-1 recognition signals, the mutant constructs behave like secretory cargoes— i.e., the mutant constructs traverse the Golgi and reach the plasma membrane. We tested this interpretation by quantifying the appearance of the mutant furin and CPD RUSH constructs at the plasma membrane. Such an experiment would not be feasible with wild-type constructs because delivery to the plasma membrane would be followed by rapid endocytosis, but removal of the AP-1 signals also removes the AP-2 signals that promote endocytosis (Schäfer et al., 1995; Voorhees et al., 1995; Eng et al., 1999; Teuchert et al., 1999; Kalinina et al., 2002), so mutant construct molecules that arrive at the plasma membrane should largely remain there. As a positive control, we created a RUSH construct in which an SBP-msYFP cassette was fused to the transmembrane domain and cytosolic tail of sucrase-isomaltase, which traverses the Golgi to the plasma membrane and remains there, with minimal endocytosis (Liu et al., 2018; Welch et al., 2021). At various times after biotin was added to release the RUSH constructs from the ER, the cells were fixed with formaldehyde to preserve the integrity of the plasma membrane, and then surface-localized msYFP was detected by adding an anti-GFP antibody followed by a far-red fluorescent secondary antibody. Thus, for each construct, the entire population of molecules was visible in the yellow channel while the plasma membrane-localized pool was visible in the far-red channel. The sucrase-isomaltase construct was present in the ER before biotin addition, and it was concentrated in the Golgi 15 min after biotin addition (Figure 9A). At 30 min after biotin addition, the sucrase-isomaltase construct became detectable at the plasma membrane, and this plasma membrane signal continued to increase until the final time point at 90 min (Figure 9A,D). Similar kinetics of cell surface appearance were seen for the mutant furin construct (Figure 9B,D) and the mutant CPD construct (Figure 9C,D), although those signals showed less of an increase after 60 min, perhaps due to residual endocytosis. The appearance of the mutant CPD construct at the cell surface is consistent with an earlier pulse-chase analysis of a CPD construct with a truncated cytosolic tail lacking the acidic AP-1 signal (Kalinina et al., 2002). These data indicate that removal of the AP-1 signals generates proteins that traverse the Golgi like secretory cargoes, as would be expected if AP-1 normally prevents the escape of resident TGN proteins to the plasma membrane.

**Figure 9.**
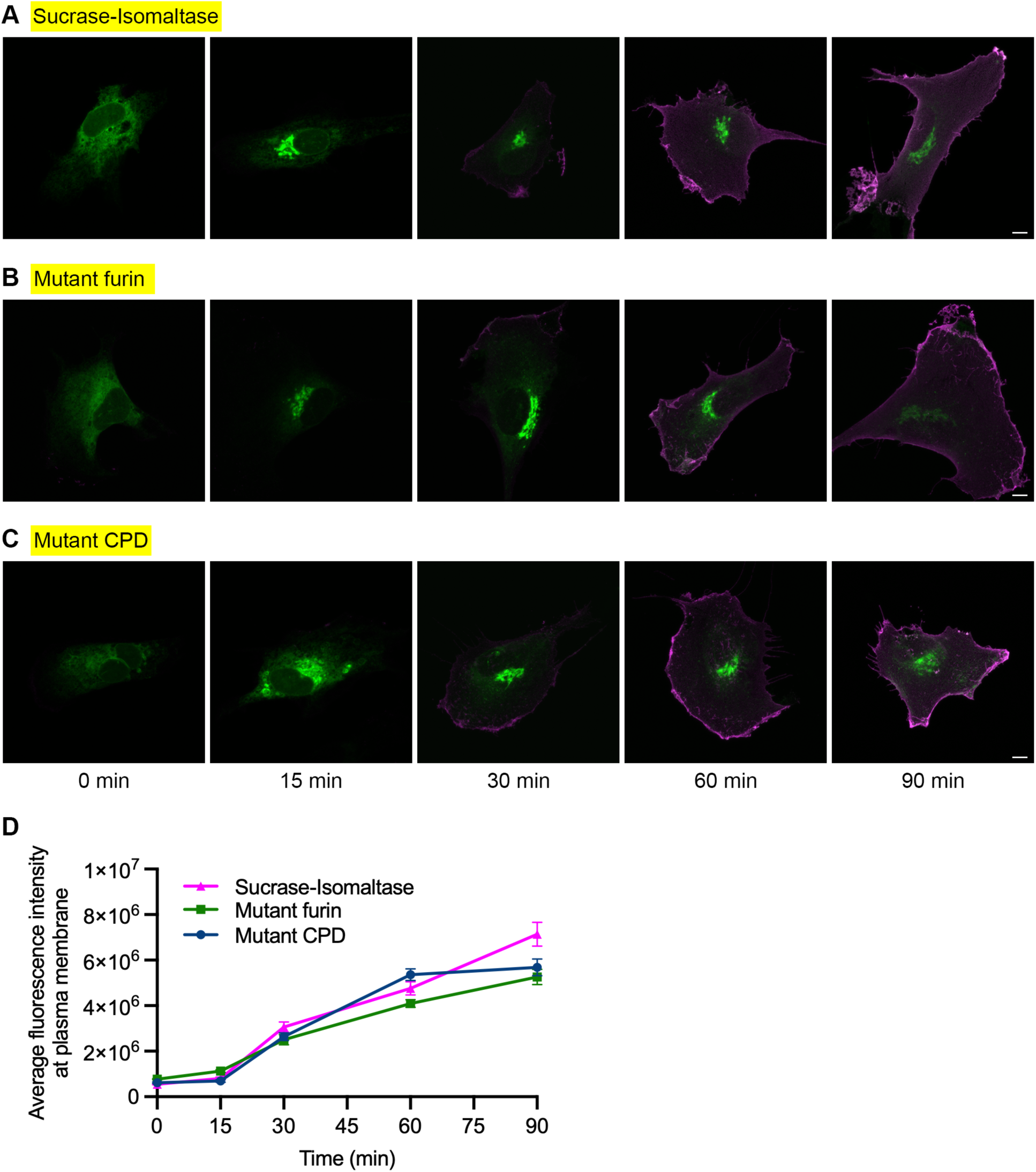
Loss of AP-1 binding allows resident TGN proteins to reach the plasma membrane. Scale bars, 5 µm. **(A)** As described in Methods, Flp-In RPE-1 cells were stably transfected with a RUSH construct for expressing a luminally msYFP-tagged sucrase-isomaltase construct. After a 2-h pretreatment with cycloheximide, biotin was added to initiate traffic through the secretory pathway. At the indicated times after biotin addition, samples were fixed with formaldehyde but not permeabilized. An anti-GFP antibody was added to bind the plasma membrane-localized pool of the construct, and this primary antibody was labeled by addition of an Alexa 647-modified secondary antibody. Then fluorescence microscopy was used to detect the total population of the construct (green) and the plasma membrane-localized pool (magenta). Representative images are shown. **(B)** Same as (A), except with a mutant msYFP-furin construct lacking the AP-1 recognition signals as described in Figure 8A. **(C)** Same as (A), except with a mutant msYFP-CPD construct lacking the AP-1 recognition signals as described in Figure 8A. **(D)** Ǫuantification of the Alexa 647 fluorescence signals from (A) - (C). For each time point, approximately 10 cells were analyzed from each of 3 separate biological replicates. Error bars represent SEM.

## Discussion

To understand what AP-1 does, we need to know where it acts. In mammalian cells, a large fraction of the membrane-associated AP-1 is at the Golgi (Meyer et al., 2000; Guo et al., 2013). This localization has been taken to indicate a role for AP-1 in anterograde traffic out of the Golgi, but it could equally well reflect a role for AP-1 in the intra-Golgi recycling of TGN proteins during cisternal maturation. In yeast, AP-1 resides at the late Golgi, and it retrieves TGN proteins from older to younger cisternae (Day et al., 2018; Casler and Glick, 2020; Casler et al., 2022; Glick, 2026; Krahn et al., 2026b). We propose that Golgi-associated mammalian AP-1 has the same function of recycling resident TGN proteins.

This hypothesis predicts that loss of AP-1 should enable TGN proteins to escape from the Golgi. To test this prediction, we knocked out the µ1A subunit of AP-1. Analysis of µ1A KO cells confirmed and extended earlier reports that inactivating AP-1 leads to selective depletion from the Golgi of various TGN proteins including furin, CIMPR, CDMPR, ATP7A, CPD, TVP23B, STX16, STX6, and Vti1a, all of which have been identified as AP-1 cargoes (Hirst et al., 2012). Depletion of TGN proteins from the Golgi in µ1A KO cells is often incomplete, probably because most TGN proteins contain endocytosis signals that provide a salvage pathway to return the escaped proteins from the plasma membrane to the Golgi (Schäfer et al., 1995; Voorhees et al., 1995; Eng et al., 1999; Teuchert et al., 1999; Kalinina et al., 2002; Ghosh et al., 2003). CIMPR and CDMPR are interesting because they are recognized not only by AP-1 but also by GGAs, which deliver mannose 6-phosphate receptors from the Golgi to endosomes (Ghosh et al., 2003). We speculate that as Golgi cisternae mature, some of the mannose 6-phosphate receptor molecules elude capture by GGAs, and then AP-1 retrieves those mannose 6-phosphate receptor molecules to younger cisternae to prevent escape from the Golgi and to give GGAs another chance at capture. If AP-1 is lost, mannose 6-phosphate receptors that elude capture by GGAs would escape to the plasma membrane. This concept is a new way of interpreting the findings that AP-1 binds to CIMPR and CDMPR and that loss of AP-1 depletes CIMPR and CDMPR from the Golgi while causing more frequent internalization from the plasma membrane (Meyer et al., 2000; Meyer et al., 2001; Doray et al., 2002; Robinson et al., 2024).

A limitation of using immunofluorescence to examine TGN proteins in µ1A KO cells is that long-term loss of AP-1 could have indirect effects (Robinson et al., 2010), so to test directly whether AP-1 keeps resident TGN proteins from escaping to the plasma membrane, we used live-cell imaging. Our analysis focused on furin, which has AP-1-dependent TGN localization signals in its cytosolic tail (Schäfer et al., 1995; Teuchert et al., 1999). By fusing the transmembrane domain and cytosolic tail of furin to a luminal fluorescent protein plus a reversible ER retention tag, we generated a furin fusion protein that could be accumulated in the ER and then released in a cargo wave for delivery to the Golgi (Boncompain et al., 2012; Tie et al., 2016). This wild-type furin fusion protein largely remained in the Golgi. By contrast, when the AP-1 signals were mutated, the mutant furin fusion protein exited the Golgi. In µ1A KO cells, both the wild-type and mutant furin fusion proteins exited the Golgi. Moreover, when cargo waves were generated in wild-type cells using a wild-type CPD fusion protein or a mutant CPD fusion protein lacking the AP-1 recognition signals, the wild-type CPD fusion protein largely remained in the Golgi whereas the mutant CPD fusion protein exited the Golgi. An assay for arrival at the plasma membrane confirmed that in wild-type cells, the mutant furin and CPD constructs reached the plasma membrane with similar kinetics as a secretory membrane protein. Thus, when AP-1 fails to bind, fusion proteins of furin and CPD behave like secretory cargoes that transit rapidly through the Golgi. AP-1 normally acts to keep those fusion proteins in the TGN. Given the well-established role of AP-1 in generating clathrin-coated vesicles (Hirst et al., 2012; Duncan, 2022; Robinson et al., 2024), a simple interpretation is that AP-1 vesicles continually recycle furin and CPD from older to younger TGN cisternae. It remains possible that AP-1 transports furin and CPD to endosomes followed by recycling to the TGN, but based on evidence that Golgi-associated tethers capture AP-1 vesicles (Brown et al., 2011; Navarro Negredo et al., 2018; Krahn et al., 2026a), we consider it more likely that AP-1 directly mediates intra-Golgi recycling.

Our analysis of the mammalian cell data aligns well with the role of AP-1 in yeast (Glick, 2026). For example, yeast Kex2 is a processing protease related to furin, and the primary mechanism that localizes Kex2 to the TGN is intra-Golgi recycling via the AP-1/Ent5 pathway (Casler et al., 2022; Glick, 2026; Krahn et al., 2026b). The parallels between mammalian and yeast cells were emphasized by harnessing the knowledge that CPD is a TGN-localized mammalian carboxypeptidase and an AP-1 cargo (Arolas et al., 2007; Hirst et al., 2012). We predicted that Kex1, a TGN-localized yeast carboxypeptidase with the same topology and enzymatic function as CPD (Cooper and Bussey, 1992; Bryant and Boyd, 1993; Lussier et al., 1995), would recycle together with Kex2 in the AP-1/Ent5 pathway. That result was indeed observed. The implication is that in diverse eukaryotes, intra-Golgi recycling pathways are similar regarding not only their mechanisms, but also their cargoes.

The finding that the mammalian Golgi has both AP-1 and GGA adaptors was initially mysterious, but it may have a simple explanation: AP-1 nucleates clathrin-coated vesicles that recycle TGN proteins within the Golgi, while GGAs nucleate clathrin-coated vesicles that carry lysosomal hydrolases from the Golgi to endosomes (Figure 10). Yeast cells have an analogous division of labor between AP-1 and GGAs (Glick, 2026). A plausible synthesis is that in diverse eukaryotes, Golgi-associated AP-1 maintains the localization of TGN proteins while Golgi-associated GGAs deliver newly synthesized acid hydrolases to the degradative system.

**Figure 10.**
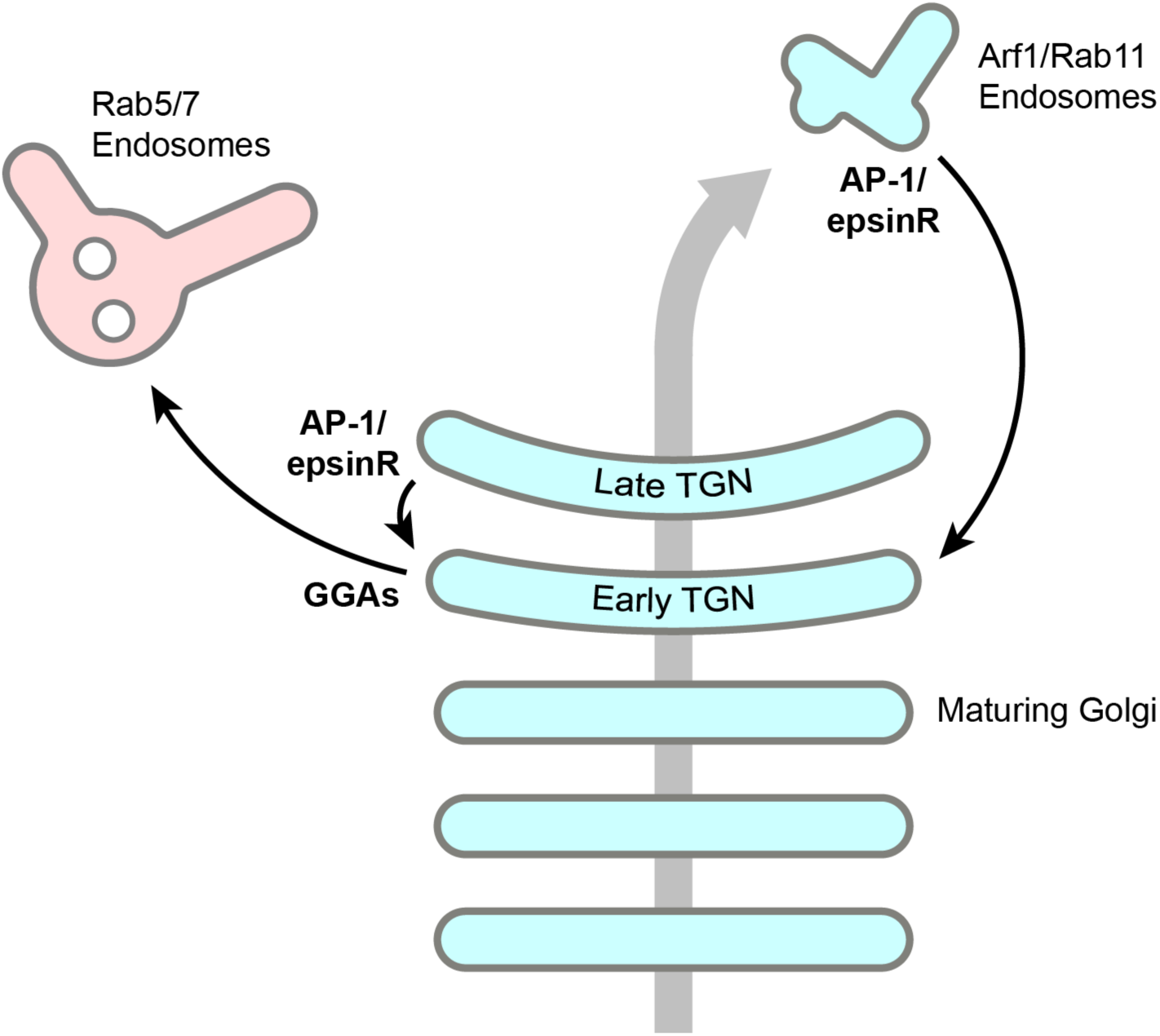
Model for the roles of mammalian AP-1 and epsinR in intracellular trafficking. The AP-1 and epsinR clathrin adaptors are present at the TGN and at Arf1/Rab11 endosomes, which are also known as recycling endosomes. The Golgi is presumed to undergo cisternal maturation as indicated by the gray arrow, and Arf1/Rab11 endosomes are presumed to arise from maturing TGN cisternae. These secretory compartments are biochemically and functionally distinct from endocytic Rab5/7 endosomes. Our data suggest that AP-1 and epsinR operate in retrograde pathways that recycle TGN proteins to keep those proteins in the Golgi. AP-1 and epsinR likely have additional functions in delivering certain internalized proteins from Arf1/Rab11 endosomes to the TGN. GGA clathrin adaptors are also present at the TGN, where they operate in an anterograde pathway to deliver newly synthesized lysosomal hydrolases to Rab5/7 endosomes.

In mammalian cells, there is more to the story because a large fraction of the membrane-associated AP-1 is on endosomes. To interrogate the additional functions of mammalian AP-1, it is crucial to clarify which types of endosomes harbor AP-1. Mammalian endosomes are termed early, late, or recycling endosomes based on differences in function and composition (Scott et al., 2014; Naslavsky and Caplan, 2018). In studies that examined AP-1, these various compartments have sometimes been grouped together under the generic term "endosomes". We propose that analyses of membrane traffic can be made more precise by classifying endosomes in both mammalian and yeast cells into two distinct categories, as follows.

The first category can be described as "Rab5/7 endosomes". In mammalian cells, these compartments label with the Rab5 and Rab7 GTPases at life cycle stages termed early endosomes and late endosomes, respectively (Wandinger-Ness and Zerial, 2014). The yeast Rab5/7 endosomes are prevacuolar (late) endosomes, which label with the Rab5 homologs Vps21 and Ypt52 and the Rab7 homolog Ypt7 (Epp et al., 2011; Day et al., 2018). Rab5/7 endosomes in mammalian and yeast cells share the following properties: they contain the lipid phosphatidylinositol 3-phosphate, they contain ESCRT proteins that generate intraluminal vesicles, they serve as intermediate destinations for acid hydrolases en route from the Golgi to lysosomes or vacuoles, and they contain sorting nexins that generate transport carriers.

The second category can be described as "Arf1/Rab11 endosomes". In mammalian cells, these compartments are termed recycling endosomes, and they label with the Arf1 and Rab11 GTPases (Wandinger-Ness and Zerial, 2014; Stockhammer et al., 2024). Recycling endosomes are closely related to the TGN—many of them are found near the TGN (Fujii et al., 2020), and they share components with the TGN including Arf1, the BIG1/2 guanine nucleotide exchange factors, subunits of the GARP/EARP vesicle tethers, and the lipid flippase ATP8A1 (Shin et al., 2004; Schindler et al., 2015; Pocognoni et al., 2024; Stockhammer et al., 2024). In yeast cells, the TGN also functions as an early and recycling endosome (Day et al., 2018) and it contains Arf1, the Rab11 homologs Ypt31/32, the BIG1/2 homolog Sec7, the GARP tether, and the ATP8A1 homolog Drs2 (Conibear et al., 2003; Losev et al., 2006; Liu et al., 2008; Kim et al., 2016; Thomas et al., 2021).

Based on this classification system, our results firmly place mammalian AP-1 on Arf1/Rab11 endosomes (Figure 10). This conclusion fits with a number of prior reports (Guo et al., 2013; Matsudaira et al., 2015; Stockhammer et al., 2024). Thus, AP-1 can be added to the list of components that are found at both the TGN and recycling (Arf1/Rab11) endosomes.

The question then becomes, what is the function of AP-1 at mammalian recycling endosomes? An appealing idea is that recycling endosomes emerge from the TGN (Gershlick and Bottanelli, 2026; Satoh, 2026). If so, then resident TGN proteins may occasionally partition into newly formed recycling endosomes, and AP-1 would retrieve those proteins to the TGN (Robinson et al., 2024). This role of recycling endosome-associated AP-1 would mirror the role of Golgi-associated AP-1. Recycling endosome-associated AP-1 likely has an additional function: certain proteins reach recycling endosomes via the endocytic pathway, and AP-1 would then deliver those internalized proteins to the TGN (Tran et al., 2007; Holloway et al., 2013; Matsudaira et al., 2015; Buser and Spang, 2023; Deng et al., 2026). In both cases, the inference is that recycling endosome-associated AP-1 mediates retrograde traffic to the TGN (Figure 10).

For proteins such as CIMPR and CDMPR, AP-1-dependent recycling is only part of a complex trafficking itinerary (Ghosh et al., 2003), but for proteins such as furin and CPD that seem to function exclusively at the TGN, AP-1-dependent recycling may be the primary localization mechanism. This concept should be evaluated in the context of previous conclusions that furin and CPD visit multiple organelles, including Rab5/7 endosomes and the plasma membrane (Varlamov and Fricker, 1998; Thomas, 2002; Silva-Ríos et al., 2025). We suspect that furin and CPD occasionally either (a) travel from the TGN to the plasma membrane and then undergo endocytosis to reach Rab5/7 endosomes, or (b) travel directly from the TGN to Rab5/7 endosomes. However, those routes are probably minor, in which case subsequent retrieval from Rab5/7 endosomes to the TGN would represent a salvage mechanism. Many Golgi proteins probably follow a primary recycling pathway plus secondary "leak" pathways (Krahn et al., 2026b). For example, yeast Kex2 recycles primarily via the intra-Golgi AP-1/Ent5 pathway, but it sometimes leaks to prevacuolar endosomes and perhaps also to the plasma membrane (Glick, 2026). With both mammalian and yeast cells, a trafficking study that imposes an artificial condition—such as overexpression or a transport block—can exaggerate a secondary sorting pathway and give a misleading impression. The distinction between primary and secondary sorting pathways for resident TGN proteins merits further attention.

Exploration of these various ideas will be aided by clarifying how AP-1 vesicles are captured at the TGN. Coiled-coil golgins with GRIP domains have been implicated in the capture of AP-1 vesicles at the mammalian TGN (Brown et al., 2011; Navarro Negredo et al., 2018), and we have evidence for a similar role of the sole GRIP domain golgin in yeast (Krahn et al., 2026a). In mammalian cells, certain GRIP domain golgins interact with the protein TBC1D23, and loss of TBC1D23 reportedly depleted CPD from the Golgi (Shin et al., 2017; Navarro Negredo et al., 2018; Cattin-Ortolá et al., 2024). It therefore seemed possible that TBC1D23 would be a key player in AP-1-dependent recycling. However, in the cell line we examined, a knockout of TBC1D23 did not noticeably deplete CPD or furin from the Golgi. A knockout of TBC1D23 did seem to enhance the accumulation of CIMPR in endosomes, consistent with a possible role for TBC1D23 in endosome-to-Golgi traffic (Shin et al., 2017; Huang et al., 2019; Cattin-Ortolá et al., 2024). On the other hand, TBC1D23 interacts with the WDR11 complex, and it has been suggested that the WDR11 complex plays a role in AP-1 vesicle targeting (Navarro Negredo et al., 2018; Deng et al., 2024). Thus, although GRIP domain golgins may help to capture AP-1 vesicles, the specific roles of GRIP domain golgins and their accessory proteins remain murky.

Another potential way to study AP-1 vesicle targeting is to characterize the SNARE machinery involved in the fusion of AP-1 vesicles with the TGN. Little information is available in this regard, but the Ǫc-SNARE GS15 was implicated in retrograde traffic from recycling endosomes to the TGN (Tai et al., 2004), and that pathway is now thought to be mediated by AP-1. Interestingly, we find that in µ1A KO cells, GS15 is depleted from the Golgi and degraded, suggesting that GS15 is an AP-1 cargo. The yeast ortholog of GS15 is Sft1, which plays a role in intra-Golgi recycling at a late stage of cisternal maturation (Wooding and Pelham, 1998; Xu et al., 2002). Further studies of GS15 and Sft1 should help to illuminate the molecular basis of AP-1-dependent retrograde traffic.

A long-standing question is the relationship between AP-1 and epsinR, which has been described as a clathrin adaptor that interacts with AP-1 (Hirst et al., 2003; Mills et al., 2003). The most thorough study of epsinR to date used acute inactivation to conclude that epsinR had a broad effect on clathrin-coated vesicle formation (Hirst et al., 2015). We became interested in epsinR after realizing that in yeast, Ent5 exhibits functional redundancy with AP-1 and can mediate intra-Golgi recycling of TGN proteins in the absence of AP-1 (Casler et al., 2022). However, the capabilities of epsinR seem to be more restricted than those of Ent5. In mammalian cells containing epsinR, loss of AP-1 depletes a wide range of TGN proteins from the Golgi, suggesting that epsinR cannot mediate intra-Golgi recycling without AP-1. Conversely, in cells containing AP-1, loss of epsinR depletes only a few TGN proteins from the Golgi, suggesting that AP-1 can mediate intra-Golgi recycling of many TGN proteins without help from epsinR. A straightforward interpretation is that epsinR assists AP-1 by playing an auxiliary role that requires the presence of AP-1.

Our structure predictions using AlphaFold indicate that epsinR and Ent5 may associate with AP-1 in a conserved manner. In the predicted complexes, two copies of Arf1 bridge the interaction of an AP-1 tetramer with two epsinR or Ent5 molecules. Although this model remains to be tested experimentally, it fits with the finding that epsinR is abundantly present in clathrin-coated vesicles, even more so than AP-1 (Hirst et al., 2015).

The relationship between epsinR and the SNAREs Vti1a and Vti1b is intriguing. epsinR has been shown to bind Vti1b (Chidambaram et al., 2004; Hirst et al., 2004; Miller et al., 2007), and that interaction is compatible with our predicted epsinR:Arf1:AP-1 interaction. But Vti1a and Vti1b have only about 30% sequence identity, and epsinR apparently does not bind Vti1a (Miller et al., 2007; Empereador-Melero et al., 2019). On the other hand, loss of epsinR strongly depletes both Vti1a and Vti1b from the Golgi, comparable to the effect seen with loss of AP-1 (this study and Hirst et al., 2004). A possible explanation is that epsinR influences Vti1a and Vti1b in different ways: epsinR would indirectly assist AP-1 in the intra-Golgi recycling of Vti1a, whereas epsinR would bind directly to Vti1b to mediate its intra-Golgi recycling with the aid of AP-1.

During budding yeast evolution, the epsinR-related proteins became functionally specialized. In addition to Ent5, which interacts with AP-1 and acts at the same stage of cisternal maturation as AP-1, yeast cells contain Ent3, which acts earlier than Ent5 to mediate delivery of Vti1 to Rab5/7 endosomes (Chidambaram et al., 2004; Daboussi et al., 2012; Casler et al., 2022). Thus, whereas mammalian Vti1a and Vti1b apparently recycle within the Golgi, yeast Vti1 apparently cycles between the Golgi and Rab5/7 endosomes. The two postulated roles of mammalian epsinR—assisting AP-1 with the intra-Golgi recycling of TGN proteins plus acting as an adaptor for Vti1b—correspond in yeast to AP-1/Ent5-mediated intra-Golgi recycling of TGN proteins plus Ent3-mediated Golgi-to-Rab5/7 endosome traffic of Vti1. Based on the available data, we propose that by analogy to the yeast AP-1/Ent5 recycling pathway, the mammalian pathways that retrieve TGN proteins from maturing TGN cisternae and from recycling endosomes can be provisionally designated "AP-1/epsinR" pathways (Figure 10).

Some authors have concluded that mammalian AP-1 mediates forward traffic of specific secretory cargoes to the plasma membrane (reviewed in Tan and Gleeson, 2019; Duncan, 2022). Those conclusions were based on phenotypes seen after inactivating AP-1. A caveat is that loss of AP-1 alters the secretory pathway by depleting multiple TGN proteins, by promoting tubulation of the TGN, and by slowing the Golgi export of certain secretory cargoes (this study and Stockhammer et al., 2024). Slow Golgi export might be a secondary consequence of altering the TGN, and it might be more severe for some secretory cargoes than for others. Alternatively, we cannot exclude the possibility that mammalian cells employ AP-1 for an evolutionarily novel role in the Golgi-to-plasma membrane traffic of certain secretory cargoes. Despite these remaining questions, the data indicate that the primary and conserved role of AP-1 is to maintain the localization of TGN proteins by mediating their retrograde intra-Golgi traffic.

## Materials and methods

### Plasmids

Plasmids were constructed by standard methods, including In-Fusion cloning (TaKaRa). Synthetic codon-optimized gene fragments were ordered from Integrated DNA Technologies or Twist Bioscience. All plasmid sequences were fully verified using Plasmidsaurus. Furin and CPD constructs were generated either by primer-directed mutagenesis or by restriction cloning of gene fragments encoding wild-type or mutant tail sequences. The plasmids developed for this study were deposited with Addgene together with annotated SnapGene files.

### Reverse transcription quantitative real-time PCR (RT-qPCR)

RNA was extracted from wild-type RPE-1 and HBE cells as follows. In a 6-well plate, 100,000 RPE-1 cells or 300,000 HBE cells were seeded in each of 3 wells two days before lysis, and the media was changed the day before lysis. Each well was used to create a separate lysate, yielding 3 biological replicates per cell line. To generate a lysate, the well was washed with phosphate-buffered saline (PBS) and then TRIzol (Thermo Fisher) was added. 1 ml of TRIzol was used for the RPE-1 cells, which were ∼80% confluent, and 0.5 ml of TRIzol was used for the HBE cells, which were ∼50% confluent. After pipetting to lyse the cells, the lysates were transferred to low nucleic acid binding microcentrifuge tubes. The TRIzol-preserved cell samples were stored at -20 °C for ∼2 weeks before RNA extraction. Then the samples were thawed, chloroform was added at a volume of 200 µl for the RPE-1 cells or 100 µl for the HBE cells, and the tubes were mixed by inversion. After 3 min at room temperature, the samples were spun for 15 min at 12,000xg at 4 °C. For each sample, the upper RNA-containing aqueous layer was carefully removed and transferred to a new low nucleic acid binding microcentrifuge tube. Then the RNA was cleaned using the RNA Clean C Concentrator Kit (Zymo Research) according to the manufacturer’s instructions. Each sample was ultimately eluted in 30 µl of H_2_O. The purified RNA samples were stored at -20 °C for ∼2 weeks, avoiding freeze-thaw cycles. The RNA concentrations ranged from 284-405 ng/µl for the RPE-1 cells or 145-187 ng/µl for the HBE cells. The A_260_/A_280_ ratios for the samples were all above 2.0, indicating high purity.

For qPCR, the RNA samples were thawed on ice and then reverse transcribed using the ReadyMade™ Random Hexamers (Integrated DNA Technologies) and the MMLV High Performance Reverse Transcriptase Kit (Biosearch Technologies) according to the manufacturers’ instructions. The resulting cDNA samples were used immediately for qPCR with the PowerUP SYBR Green Master Mix (Thermo Fisher #A25742). Two primer pairs were used for each gene, as follows. For µ1A (Ensembl reference sequence ENST00000291439.8), the primer pairs were: CTGTATCTGGTTGCCACATC plus ACCTGCACCACCTTATAGA, and TCTCAACACCCACGTCAA plus CTTGATCATGTACTCGATGCG. For µ1B (GenBank reference sequence AF020797.2), the primer pairs were: TCCCTCCCCGACTCCTAAGT plus GTGGCCACCAAGTAGAGGTT, and GTCAACCTGCTGGTCAAT plus CCTGACAGAAACACCTTGAG. The control gene was GAPDH (Ensembl reference sequence ENST00000229239.10), and the primer pairs were: ACAGTCAGCCGCATCTTCTTT plus CAATACGACCAAATCCGTTGACT, and TCATCAGCAATGCCTCCT plus GTCCTTCCACGATACCAAAG. In each case, one primer in the pair spanned an exon-exon junction. Each primer pair was used for 4 technical replicates of each of the 3 biological replicates. The qPCR (144 total reactions) was performed in a 384-well plate on a CFX384 Touch Real-Time PCR Detection System (Bio Rad) using the following parameters. The samples were first kept at 50 °C for 2 mins and then had an initial denaturation step at 95 °C for 10 mins. Samples were denatured at 95 °C for 0.15 min and then allowed to anneal and elongate at 60 °C for 1 min. This process was repeated for 39 cycles. The plate fluorescence was measured at the end of each cycle. After the cycles were complete, a melting curve was generated by reading the plate as the temperature increased from 65 °C to 95 °C in 0.5 °C increments over a period of 1.5 min.

To analyze the data, the raw Cq (Cycle of Ǫuantitation) values between technical replicates were first averaged, and GraphPad Prism’s Grubbs’ test was performed on each set of replicates to determine any statistical outliers. Four individual Cq values were removed from the technical averages through this method. The standard deviation between technical replicates was less than or equal to 0.5 for all primer pairs and biological replicates. The Cq values for each primer pair were normalized to GAPDH within biological replicates. The 2^-dCq^ approach was used to transform the raw data, where dCq is equal to the normalized Cq value. The resulting HBE values were averaged among biological replicates for each primer pair, and the fold-change difference of µ1A and µ1B expression was determined for each RPE-1 and HBE replicate. The average and standard deviation between biological replicates was calculated for each primer pair. This analysis was performed for both GAPDH primer pairs, but only the results from GAPDH primer pair #1 are shown. An unpaired t-test was performed, with the fold-change differences grouped into an RPE-1 group and an HBE group. There was no statistically significant difference between the two groups when comparing the values for both primer pairs of µ1A and µ1B or when comparing the values for just both primer pairs of µ1B. The same was true using the values normalized to GAPDH primer pair #2. GraphPad Prism was used to plot the fold-change differences in expression for each primer pair.

To confirm specificity of the qPCR reaction, the 384-well plate was sealed and stored at 4 °C. The qPCR reactions were run on an agarose gel, and both primer pairs for µ1A and µ1B showed only one band of the correct expected size for both RPE-1 and HBE cell types. The qPCR products were sequenced by Plasmidsaurus to confirm that the correct transcripts were amplified.

### Mammalian cell lines

RPE-1 cells, COS-7 cells, U2OS cells, and BJ-5ta cells were obtained from the American Type Culture Collection (ATCC). HBE cells (16HBE14o^−^ strain, originally from Dieter Gruenert) were a kind gift of Heike Fölsch. The growth media were as follows. RPE-1, COS-7, and BJ-5ta cells: DMEM/F-12 (Gibco) + 10% fetal bovine serum (FBS; Gibco); U2OS cells: McCoy’s 5A Medium (Gibco) + 10% FBS; HBE cells: MEM (Gibco) + 10% FBS + 2 mM L-glutamine (Invitrogen) for use in fibronectin-coated flasks (Nokes et al., 2008). For live-cell microscopy, cells were shifted to imaging medium: FluoroBrite DMEM (Gibco) + 10% FBS + 1X GlutaMAX Supplement (Gibco) the day before imaging. Transfections were performed by electroporation using a Thermo Fisher Neon system (Voltage = 1350, Width = 20, Pulses = 3) or Neon NxT system (Voltage = 1260, Width = 20, Pulses = 3). After transfection, the FBS concentration was raised to 20% for 1-2 days to enhance recovery.

The RPE-1 Flp-In cell line was generated as follows. 1x10^6^ low-passage RPE-1 cells were transfected with 1 µg of the PvuI-linearized plasmid pFRT-PuroR-GFP, which was derived from Thermo Fisher’s pFRT/lacZeo vector by replacing the lacZ-Zeocin gene with the EGFP-puro fusion gene from pEGFP-puro (Addgene plasmid #45561; Abbate et al., 2001). After 2 days in growth medium, puromycin was added at a concentration of 10 µg/ml. The cells were maintained in this selection medium for 10 days. Surviving clones were then expanded, trypsinized, and pooled in a single 10-cm dish. The resulting mixed population was expanded until it reached 90% confluence. Single GFP-positive cells were sorted by flow cytometry into 96-well microplates. Clones were expanded and checked for GFP fluorescence and puromycin resistance. One clone was subjected to whole-genome sequencing by Plasmidsaurus. A fragment of pFRT-PuroR-GFP containing the SV40 promoter, FRT cassette, EGFP gene, PuroR gene, and SV40 poly(A) signal was inserted as a single copy in the first intron of the HTR2B gene, which is nested within an intron of the PSMD1 gene on chromosome 2.

The knockout cell lines were generated using TrueCut HiFi Cas9 Protein (Invitrogen) by following the manufacturer’s protocol, as follows. For the µ1A knockout ("µ1A KO") cell line, the original RPE-1 cell line was transfected to introduce the purified HiFi Cas9, a pair of single guide RNAs (sgRNAs; 5’-CTCATCTGCCGGAACTACCGT-3 and 5’-TTCCGAGTACTTCAAGGAGC-3’) targeting exons 2 and 4 of the AP1M1 gene encoding µ1A, and the plasmid pEGFP-puro. The cells were allowed to recover for 2 days and then were placed under selection for 3 days by adding puromycin at a concentration of 10 µg/ml. Surviving cells were expanded and subsequently sorted by flow cytometry into 96-well microplates. Clones were expanded and checked for the absence of Golgi-localized AP-1 by immunofluorescence using an antibody against the γ1 subunit. For the TBC1D23 knockout cell line ("TBC1D23 KO"), which was derived from the original RPE-1 cell line, the procedure was similar except that the sgRNA was 5’-TCAGTAGATGTATCCAGCTA-3’ targeting exon #4 of the TBC1D23 gene, and the clones were checked by immunofluorescence using an antibody against TBC1D23. For the epsinR KO and AP-1/epsinR double knockout cell lines, the procedure was similar except the sgRNA was 5’-GCAACGAACGATGATCCTTG-3’ targeting exon 2 of the CLINT1 gene encoding epsinR, and the clones were checked by immunofluorescence using an antibody against epsinR. The knockout clones used here were further analyzed by sequencing of PCR products obtained from genomic DNA. For this purpose, genomic DNA was extracted using the PureLink Genomic DNA Mini Kit (Invitrogen). The following PCR primers were used. µ1A: 5’-CTGGGTCTGTTTCAATTAAAGCTGCT-3’ and 5’-ATGTCATCGTGCATTGTCTGCCTG’ (Exon 2) or 5’-GGAAAGCTTTCATCCCTCACTGATGC-3’ and 5’-CACGGTTATGTGGGGGCGTAGAG-3’(Exon 4); TBC1D23: 5’-TTTGACTGTGTGTTTTCCTTGGGTTTTGG-3’ and 5’-AAGGTAACACCCCAGAAATTTTACTTTTCAC-3’ (Exon 4); epsinR: 5’-CACCCAGTAGTGTCATTATTAAGTTTTCATGTG-3’ and 5’-TTTCTTTTTTGAGCCAAATTTTTAAGGGGAATGG-3’ (Exon 2). PCR amplicons were checked for the correct sizes by agarose gel electrophoresis, and they were sequenced by the University of Chicago DNA Sequencing Core or by Plasmidsaurus. Whole-genome sequencing by Plasmidsaurus confirmed that compared to the WT cell line, the µ1A KO cell line had mutations in Exons 2 and 4 of the µ1A (AP1M1) gene but no mutations in exons of the µ1B (AP1M2) gene.

CRISPR-mediated tagging of the endogenous RAB11A and AP1M1 (µ1A) genes in the original RPE-1 cell line was performed as previously described (Stockhammer et al., 2024). After transfection, selection, and expansion, the tags were rendered fluorescent by incubating with JFX_650_ HaloTag ligand for 10 min or JFX_554_ SNAP-tag ligand for 1 h followed by an overnight wash. Then the brightest 1% (RAB11A) or 3% (AP1M1) of the cells were enriched using flow cytometry to select cells that were homozygous for the CRISPR edit, and that enriched population was used for analysis.

Stable derivatives of the RPE-1 WT Flp-In cell line were generated as follows. 1x10^6^ cells were electroporated with 2 µg of the RUSH integration plasmid plus 9 µg of a FLP-recombinase-encoding plasmid: either plasmid pOG44 (Invitrogen), which encodes the original FLP recombinase, or plasmid pOG-FLPe, which encodes the more thermostable FLPe variant (Buchholz et al., 1998). The cells were allowed to recover in growth medium for 3-4 days. Then they were split into two to three 10-cm dishes or T75 flasks (Thermo Fisher) and selected with 5 µg/ml blasticidin (Sigma) for 7-10 days. After further growth in medium lacking antibiotic, the brightest 1-3% of the cells were enriched using flow cytometry, and that enriched population was used for analysis.

Transient transfections were performed by electroporating cells with 1-5 µg of plasmid DNA. After 2 days in growth medium at 37 °C, the cells were used for microscopy. All plasmids used for transfection were either midi- or maxi-prepped using commercial kits from Zymo Research or Invitrogen.

### Immunoblotting

For each cell line to be analyzed, cells grown to 80-90% confluency in a T75 flask were dissociated using TrypLE (Thermo Fisher). The cell suspension was centrifuged for 2 min at 100xg in a tabletop centrifuge. After removal of the supernatant, the cells were resuspended in 1 ml PBS and counted using a Cellometer automated cell counter (Nexcelom). A volume corresponding to 500,000 cells was spun at 10,000xg in a microcentrifuge, and the supernatant was discarded. The pellet was resuspended in SDS sample buffer in PBS containing a protease inhibitor cocktail (1 mM ABESF, 0.8 μM aprotinin, 20 μM leupeptin, 15 μM pepstatin A, 40 μM bestatin, 15 μM E-64, 600 μM benzamidine). Then the sample was heated for 5 min at 95 °C followed by centrifugation for 15 min at 19,000xg at 4 °C to remove insoluble material. The supernatant was collected and stored at -20 °C.

For SDS-PAGE and immunoblotting, 10-20 µl of the cell lysates were loaded into lanes of a Mini-PROTEAN TGX 4-20% gradient SDS gel (Bio-Rad). The molecular weight marker was 5 µl of the Chameleon 700 pre-stained protein ladder (LICORbio). After electrophoresis at 135-150 V for 35-50 min, the proteins were transferred to a nitrocellulose membrane using the semi-dry Trans-Blot Turbo Transfer System (Bio-Rad) with the parameters 25 V, 2.5 A, 3-5 min. The membrane was incubated with blocking buffer (Tris-buffered saline (TBS) with Tween 20 (TBST) + 5% dried milk (Biotium)) for 1 h at 23 °C, then washed 3 times by shaking for 5 min with TBST at 23 °C. If desired, the membrane was cut to separate the loading control from the test protein, and the two separated membrane pieces were processed in parallel.

The membrane was incubated with shaking overnight at 4 °C with the appropriate primary antibody diluted in blocking buffer. Additional incubations the following day were performed at 23 °C. The membrane was washed 3 times with TBST and then incubated with the appropriate HRP-conjugated secondary antibody diluted in blocking buffer for 1 h. Then the membrane was washed 3 times with TBST and placed in TBS. To detect HRP, the SignalBright Max Chemiluminescent Substrate (Proteintech) was mixed and allowed to reach room temperature in the dark. Substrate was added dropwise to the membrane until the entire surface was covered, and then the membrane was immediately imaged using an Amersham ImageǪuant 800 system. The antibodies used are listed in Table S1.

### Immunofluorescence microscopy of mammalian cells

Immunofluorescence was performed as previously described (Bhattacharyya et al., 2010) and as summarized briefly below. For each well of a CultureWell MultiWell chambered glass coverslip (Grace Bio-Labs), 1000-1500 cells were seeded the day before staining. When wild-type and mutant cell lines were being compared, wild-type cells were added to the wells in the top row of a coverslip and mutant cells were added to the wells in the bottom row. In some cases, wild-type and mutant cells (500 cells of each type) were mixed in a single well. The cells were incubated in growth medium overnight at 37 °C. The coverslip was immersed in 40 ml of ice-cold acetone, rehydrated in the presence of a chemical crosslinker, and incubated with primary and secondary antibodies in a humidified chamber constructed from a Petri dish. The antibodies used are listed in Table S1. Each well of the coverslip could hold 40 µl of liquid, and the volumes of the antibody and wash solutions were determined accordingly. After antibody labeling was complete, ∼3.5 µl of SlowFade mounting solution with DAPI (Invitrogen) was applied to each well, and the coverslip was flipped over onto a labeled microscope slide.

The knockout phenotypes for µ1A, epsinR, and TBC1D23 as well as the immunofluorescence images showing AP-1 and epsinR colocalization were confirmed with a Zeiss Axio Observer widefield microscope using a Plan-Apo 1.4-NA 63x oil objective and a Lumencor light engine. Single-plane images were captured in three channels: blue (370-410 nm excitation, 430-470 nm emission), red (533-558 nm excitation, 570-640 nm emission), and far-red (625-655 nm excitation, 665-715 nm emission).

Immunofluorescence images for Figures 2 and S2 were collected with a Zeiss LSM 980 confocal microscope using a 63x oil objective. 1024 x 1024 px confocal z-stacks of 18 optical sections spaced 170 nm apart were captured in three channels configured for Alexa Fluor 647 (639 nm excitation, 641-694 nm emission), Alexa Fluor 555 (561 nm excitation, 552-632 nm emission), and DAPI (405 nm excitation, 407-551 nm emission). For each test protein, the laser power was set to maximize the detection of peripheral structures. The zoom factor was 2.5, the pixel size was 52.6 nm, the pixel dwell time was 0.768 µs, the line step was 1, the pinhole was 45.32 um, line averaging was 2, and the effective NA was 1.4. For analysis of the confocal images, the z-stacks were deconvolved using Huygens, average projected, and analyzed in ImageJ using the custom "Ǫuantify Overlap" plug-in to quantify signal overlap between the red and far-red channels. In brief, the deconvolved, average projected channels were merged using the following scheme. DAPI was in blue, the endosomal marker was in green, and AP-1 was in red. An ROI was then hand-drawn around the nucleus and Golgi to remove the Golgi-localized pool of protein from the image, leaving only the peripheral signal. Then the AP-1 channel was thresholded to preserve the brightest 2% of the signal histogram and to remove diffuse cytoplasmic AP-1 signal. The resulting image was processed with the plug-in to measure the fraction of total peripheral AP-1 signal overlapping with the endosomal marker signal. An arbitrary threshold of "Higher” for the endosomal marker channel was used to filter background prior to quantification. Values from two independent experiments encompassing a total of 38-40 cells were used to generate the final averages.

Immunofluorescence images for Figures 3-5 and Figure S3A-K were collected with a Zeiss LSM 980 confocal microscope using a 63x oil objective. 1024 x 1024 px confocal z-stacks of 20 optical sections spaced 210 nm apart were captured in three channels configured for Alexa Fluor 647 (639 nm excitation, 641-694 nm emission), Alexa Fluor 555 (561 nm excitation, 561-623 nm emission), and DAPI (405 nm excitation, 416-482 nm emission). For each test protein, the laser power was set to obtain the maximal possible signal without saturation, and this value was kept constant between replicates and between cell lines. The zoom factor was 2, the pixel size was 70 nm, the pixel dwell time was 1.02 µs, the line step was 1, the Alexa Fluor 647 channel pinhole was 1 AU (64 μm), the DAPI/Alexa Fluor 555 channel pinhole was 1 AU (43 μm), and the effective NA was 1.4. For analysis of confocal images, the z-stacks were average projected using Fiji. For each test protein, the brightness and contrast were set to be the same between replicates and between cell lines. To quantify the test protein signal at the Golgi using Fiji, a freehand ROI was drawn around the Golgi ribbon using the GM130 channel as a reference, and the integrated density of the test protein signal was measured. Then the same ROI was moved to an area outside the Golgi, and the integrated density was measured to quantify the cytosolic background signal. To make sure this second ROI was still within the cell, a high-contrast view of the test protein channel was used. The cytosolic background signal was then subtracted from the Golgi signal. For each knockout cell, the background-subtracted Golgi signal was divided by the average background-subtracted Golgi signal from the corresponding wild-type cell line. These mutant-to-wild-type ratios were then averaged and plotted. For each test protein and cell line, the plotted values represent the average of 3 biological replicates of about 10-15 cells each.

### Ǫuantification of AP-1 distribution

To quantify the fractions of membrane-associated AP-1 present at the Golgi versus peripheral puncta, RPE-1 cells expressing endogenously tagged µ1A-SNAP-tag and transiently overexpressing GalNAc-T2-msGFP2 were labeled with 1 µM JFX_650_ SNAP-tag ligand (courtesy of Luke Lavis’ lab) for 1 h at 37 °C and then placed in fresh medium (Fluorobrite DMEM + 10% FBS + 1x GlutaMax) for 2 h at 37 °C. Imaging of live cells was performed at 37 °C with a Zeiss LSM 980 confocal microscope using a 63x oil immersion objective. The resulting z-stacks were deconvolved, averaged projected, and converted to 8-bit.

Cytosolic AP-1 signal was removed by thresholding in ImageJ using the following procedure. (1) Split the channels to isolate the AP-1 channel. (2) Subtract background (a value of 10 was empirically determined and was used for all micrographs). (3) Use the Auto Local Threshold command with the Phansalkar method (Radius = 25, Parameter 1 = 0.01). (4) Combine the resulting binary image with the original projected image by an AND operation to remove the diffuse cytosolic signal. (5) Merge the resulting processed image with the GalNAc-T2 channel. (6) Measure the total AP-1 signal in the processed image. (7) Use the GalNAc-T2 channel to draw a freehand ROI around the Golgi ribbon, and measure the Golgi-associated AP-1 signal in the processed image. (8) Determine the percentage of the total AP-1 signal that was Golgi-associated. (9) Calculate the average of this percentage by analyzing 8-10 cells in each of two independent experiments.

### Time-lapse imaging of mammalian cells

For 4D microscopy, about 150,000 to 200,000 cells were seeded on 35-mm coverslip-bottom dishes (MatTek P35G-0.170-14-C), either without any manipulation or after transfection, and were kept at 37°C. For RUSH experiments, 2 µg/ml avidin was added to the media to reduce cargo leakage caused by endogenous biotin present in FBS. The day before imaging, cells were changed to imaging media (Fluorobrite DMEM, 10% FBS, 1x GlutaMax) supplemented with 2 µg/ml avidin. The day of imaging, the cells were incubated with JFX_650_ HaloTag ligand for 10 min and then placed in imaging media without avidin. For the cell line expressing SNAP-tag-labeled µ1A, there was an additional 1 h labeling with JFX_550_ SNAP-tag ligand followed by a 1 h wash in fresh growth medium and then a transfer to imaging medium. Cells were visualized with a Leica SP5 or Leica Stellaris confocal microscope using a 63x oil objective inside an incubator that maintained the temperature at 37 °C.

To measure Golgi accumulation and Golgi export kinetics, time-lapse imaging was performed with WT cells transfected to overexpress the indicated secretory cargo proteins and Golgi marker. Previously described protocols were used for RUSH (Boncompain et al., 2012) and ESCargo (Casler et al., 2020) cargo waves and for iFRAP (Patterson et al., 2008). In brief, cells were incubated with 100 µM cycloheximide (RPI) for 3 h to block protein synthesis and allow the cargo synthesized before addition of the drug to become fluorescent. To reduce phototoxicity, 500 µM of freshly made L-ascorbic acid (Sigma) was added to the medium 3 h before imaging (Harada et al., 2023). A RUSH cargo wave was then initiated by addition of 100 µm biotin, and an ESCargo cargo wave was initiated by addition of 50 µM SLF. Cells were then either immediately imaged or incubated for 15 min to allow cargo accumulation in the Golgi. Cells expressing ESCargo were incubated with SLF for 3 min before time-lapse acquisition to allow cargo disaggregation. To measure Golgi export by iFRAP, the Golgi pool of cargo was isolated 10-15 min after initiation of the cargo wave by identifying the Golgi ribbon using HaloTag-labeled GalNAc-T2 and by photobleaching all non-Golgi cargo signal in the cell. Photobleaching was performed on an SP5 confocal microscope using maximum laser power for ∼45-55 s with the 561-nm laser to bleach ESCargo or mScarlet. Imaging of post-photobleaching Golgi export commenced ∼20 min after initiation of the cargo wave in WT cells or ∼26 min after initiation of the cargo wave in µ1A KO cells, with the different timing reflecting the slower Golgi export kinetics in µ1A KO cells. Cell volumes encompassing the entire Golgi ribbon were collected every minute for 30-40 minutes using 2-5% of the 561-nm line and 5-14% of the 633-nm line. The resulting z-stacks were deconvolved using Huygens, assembled as 4D hyperstacks, and average projected with ImageJ. For the experiments of Figure 8, the images were bleach corrected with Fiji using exponential fitting. Then the amount of cargo signal overlapping with the Golgi marker at each time point was quantified using the following ImageJ macro:

t=getTitle();

run("Split Channels");

//work on entire green (Golgi marker) stack

selectWindow("C2-"+t);

run("Gaussian Blur…", "sigma=1 stack");

//run("Threshold…"); setAutoThreshold("Li dark");

//work on entire red (protein of interest) stack

selectWindow("C1-"+t);

run("Grays");

run("Gaussian Blur…", "sigma=1 stack");

//run("Threshold…");

//loop through slices creating C restoring selection and measuring

//commented out bits are internal checks to make sure Ch01 and Ch02 slices match

//can remove // to run these checks, but leave // in place when running analysis

for (n=1; n<=nSlices; n++) {

setSlice(n);

selectWindow("C2-"+t);

setSlice(n);

run("Create Selection");

//run("Measure");

selectWindow("C1-"+t);

run("Restore Selection");

//run("Set Measurements…", "area mean min integrated display redirect=None decimal=0");

//run("Measure");

run("Set Measurements…", "area mean min integrated limit display redirect=None

decimal=0");

run("Measure");

selectWindow("C2-"+t);

run("Select None");

selectWindow("C1-"+t);

run("Select None");

}

The resulting values for each time point were background-subtracted and averaged across two to three independent experiments. A total of 9-27 cells were pooled to generate the final averages.

### Ǫuantification of secretory cargo arrival at the plasma membrane

Stable cell lines expressing SBP-msYFP-tagged sucrase-isomaltase, furin, or CPD were generated using the WT Flp-In RPE-1 cell line, and the 2% brightest cells were enriched by fluorescence-activated cell sorting. The day before the experiment, 80,000 cells were seeded onto each coverslip, with one coverslip per time point per cell line. The cells were allowed to grow overnight. On the day of the experiment, the cells were treated with cycloheximide for 2 h and then were incubated with 10 µM biotin, all at 37 °C. Biotin addition was staggered according to time point—e.g., biotin was first added to the t=90 min sample, then 30 min later to the t=60 min sample, and so on. Samples were collected after 0, 15, 30, 60, or 90 min of biotin treatment. Following the biotin treatment, the medium was aspirated and immediately replaced with 4% formaldehyde in PBS (Cell Signaling Technology). The cells were fixed for 15 min at room temperature and then washed three times with PBS. There was no permeabilization step. The coverslips were then blocked and stained according to our immunofluorescence protocol (see above). msYFP-tagged proteins were detected at the cell surface by indirect immunofluorescence, where the primary antibody was rabbit polyclonal anti-GFP (1:1000) and the secondary antibody was Alexa 647-conjugated goat anti-rabbit IgG (1:1000).

The Zeiss LSM 980 confocal microscope with a 63x oil objective was used to collect 1024 x 1024 px z-stacks of 20 optical sections spaced 210 nm apart. Channels were configured for msYFP (514 nm excitation, 517-641 nm emission) and Alexa Fluor 647 (653 nm excitation, 641-694 nm emission). For each test protein, the laser power was set to obtain the maximal possible signal without saturation at the latest time point, and this value was kept constant between replicates and between cell lines. The zoom factor was 1.5, the pixel size was 90 nm, the pixel dwell time was 2.05 μs, the line step was 1, the Alexa Fluor 647 channel pinhole was 1 AU (64 μm), the msYFP channel pinhole was 1 AU (55 μm), and the effective NA was 1.4. Three biological replicates of approximately 10 cells were collected per cell line per time point. For analysis of the confocal images, the z-stacks were average projected using Fiji. The brightness and contrast were set to be the same between time points, cell lines, and replicates. The Alexa 647 signal of the whole frame was measured using ImageJ and averaged between three biological replicates. This number was plotted for each time point and cell line.

### Yeast methods

The parental haploid *S. cerevisiae* strain was JK9-3da (*leu2-3,112 ura3-52 rme1 trp1 his4*) (Kunz et al., 1993) carrying *pdr1Δ* and *pdr3Δ* mutations to facilitate HaloTag labeling (Barrero et al., 2016). For imaging, yeast cells were grown with shaking in baffled flasks at 23 °C in nonfluorescent minimal glucose dropout medium (NSD) (Bevis et al., 2002).

A construct to tag Kex1 with GFP was designed with the aid of the Saccharomyces Genome Database (Wong et al., 2023). In-Fusion cloning was used to insert a synthetic DNA fragment encoding a 3’ fragment of the *KEX1* gene fused to two copies of yeast codon-optimized msGFP2 (Valbuena et al., 2020) into the integrating vector YIplac211 (Gietz and Sugino, 1988). The resulting plasmid was linearized with XhoI for integration at the *KEX1* locus, and gene replacement was performed by the pop-in/pop-out method (Rothstein, 1991; Rossanese et al., 1999).

Additional reagents and methods used in this study were described in our previous publication (Krahn et al., 2026b), including yeast protein tagging by chromosomal gene replacement, yeast cell growth, labeling of proteins with a fluorescent HaloTag ligand (Grimm et al., 2021), preparation and use of rapamycin and CK-666, live-cell 4D confocal imaging of yeast cells attached to coverglass-bottom dishes, processing and averaging of fluorescence kinetic traces, detection and quantification of rapamycin-dependent vesicle capture at the yeast bud neck, quantification of CK-666-induced trafficking defects in *apl4Δ ent5Δ* cells, and statistical analysis.

### Structure modeling

AlphaFold3 predictions (Abramson et al., 2024) were generated using the public server (www.alphafoldserver.com). Structural models were prepared with the PyMOL Molecular Graphics System, Version 3.1.8 (Schrödinger, LLC).

### Software

Plasmids were designed using SnapGene (Dotmatics). Images were analyzed with ImageJ and assembled using Adobe Photoshop. Plotting and statistical analysis were performed using Prism (Dotmatics). Processing of 4D yeast confocal data was performed with a custom ImageJ plugin termed “Ǫuantify Overlap” (Krahn et al., 2026a), available with source code from https://github.com/bsglicker/4D-Image-Analysis.

## Acknowledgments

This work in the BSG laboratory was supported by NIH/NIGMS award R35 GM144050. FMV, ML, AHK, and CFL were supported by NIH training grant T32 GM007183, and FMV was supported by an HHMI Gilliam Fellowship. JCF was supported by NIH/NIGMS award R35 GM136258. PA and FB were supported by the Deutsche Forschungsgemeinschaft (German Research Foundation) grant TRR186 (project A20 to FB). Thanks for assistance with fluorescence microscopy to Christine Labno and Lorraine Horwitz at the Integrated Microscopy Core Facility (RRID:SCR_019197), which is supported by the NIH-funded Cancer Center Support Grant P30 CA014599. Thanks for assistance with cell sorting to Cezary Ciszewski at the Human Disease and Immune Discovery Core Facility (RRID:SCR_022936). Thanks to Lloyd Fricker for providing the CPD antibody, to Heike Fölsch for providing HBE cells and advice about qRT-PCR primers, and to Gabriela Haddad in Alex Ruthenburg’s lab for assistance with RT-qPCR. We thank the Lavis lab and Open Chemistry team (Janelia) for the gift of JFX dyes. The authors declare no competing financial interests.

## Author Contributions

Conceptualization, FMV, ML, AHK, AP, CFL, JCF, and BSG; Methodology, FMV, ML, AHK, AP, CFL, JCF, PA, FB, and BSG; Software, BSG; Investigation, FMV, ML, AHK, AP, CFL, JCF, and BSG; Resources, FMV, ML, AHK, AP, CFL, PA, FB, and BSG; Writing – Original Draft, FMV, ML, JCF, and BSG; Writing – Review and Editing, FMV, ML, AP, and BSG; Visualization, FMV, ML, AHK, AP, CFL, and JCF; Supervision, FB and BSG; Funding Acquisition, FB and JCF and BSG

FMV led the project, optimized protocols for mammalian tissue culture, immunofluorescence microscopy, iFRAP and 4D live cell imaging, genome editing, and Western blotting, performed microscopy data acquisition, processing, and quantification, and revised the manuscript and figures. ML optimized immunofluorescence protocols, collected and quantified immunofluorescence microscopy data, developed a cell surface labeling assay to assess TGN protein leakage to the plasma membrane, performed qRT-PCR to assess transcript levels in different cell lines, generated and validated stable overexpression and knockout lines, and revised the manuscript and figures. AHK identified Kex1 as a putative AP-1 cargo, generated yeast strains, and performed ectopic vesicle capture and kinetic mapping experiments. AP performed experiments with CK-666 to assess Kex1 loss from the Golgi in wild-type and AP-1/Ent5 mutant yeast strains. CFL generated yeast strains and performed kinetic mapping experiments. JCF built and analyzed the structure models in consultation with BSG, and generated Figure 6. PA and FB provided technical advice and resources to generate a double knock-in RPE-1 line expressing endogenously tagged AP-1 and Rab11. BSG procured funding and guided the project, provided conceptual and technical feedback, and generated the original manuscript and the figures other than Figure 6.

## Abbreviations

CDMPR: cation-dependent mannose 6-phosphate receptor
CIMPR: cation-independent mannose 6-phosphate receptor
CPD: carboxypeptidase D
FKBP: FK506-binding protein
FRB: FK506-rapamycin binding domain
iFRAP: inverse fluorescence recovery after photobleaching
ipTM: interface predicted template modeling
PAE: predicted aligned error
PI4P: phosphatidylinositol 4-phosphate
RT-qPCR: quantitative reverse transcription PCR
RUSH: retention using selective hooks
TGN: *trans*-Golgi network

---

**Figure S1.**
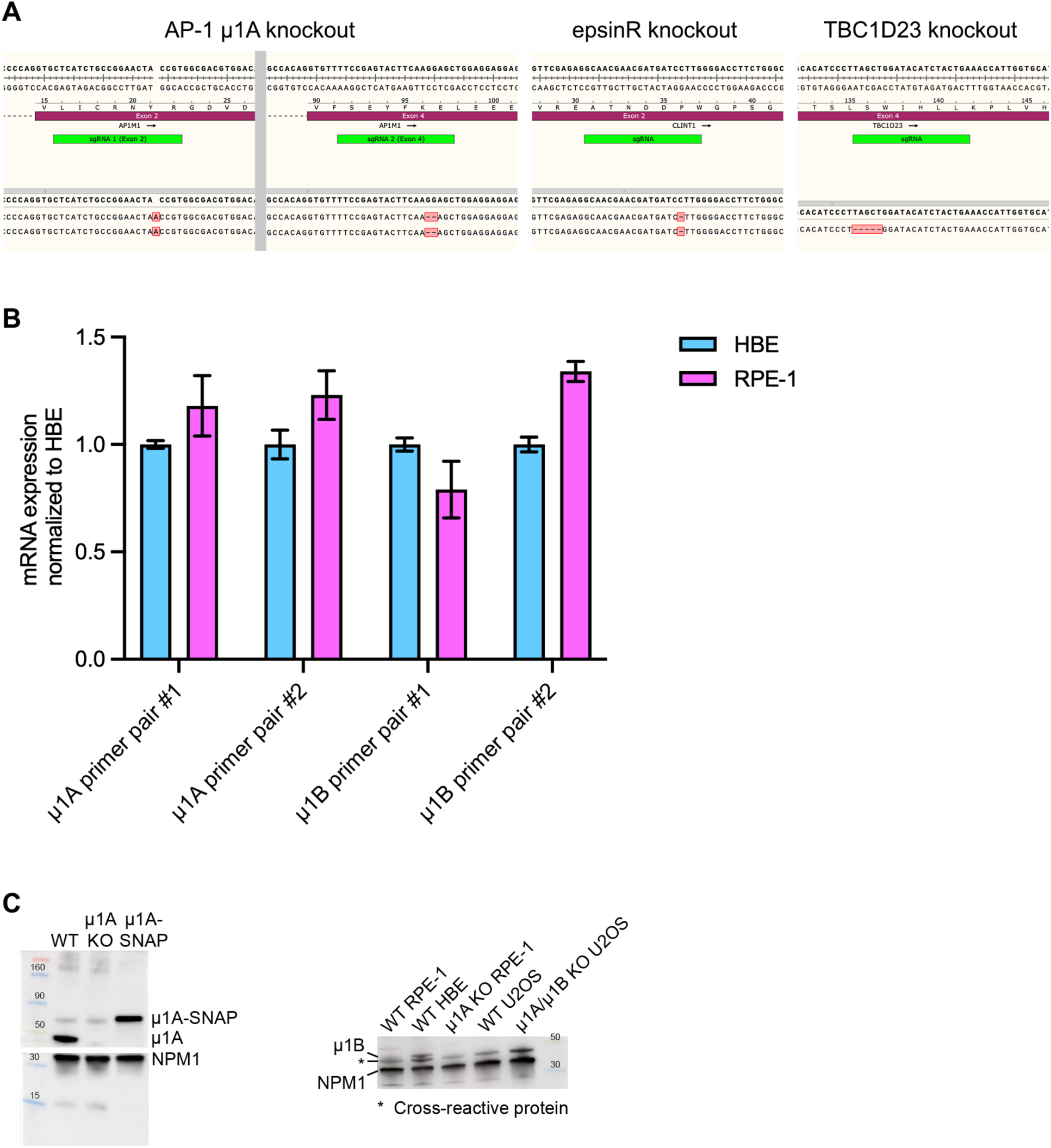
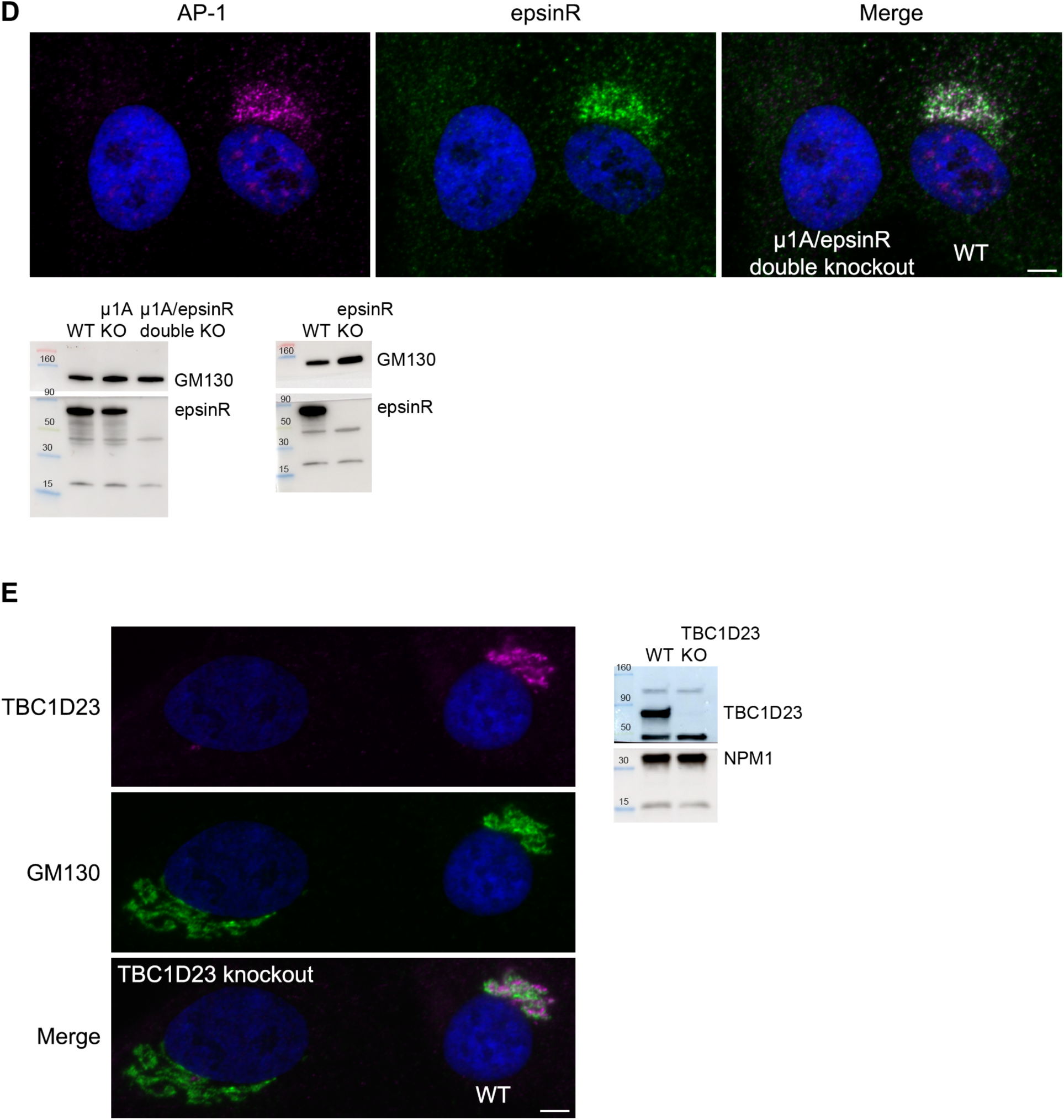
Supplementary data for the knockout cell lines. **(A) *Left:*** Confirmation of the mutations in the µ1A gene. Shown at the top are portions of the µ1A (AP1M1) gene sequence, together with the binding sites for two sgRNAs that were used for CRISPR/Cas9-mediated gene disruption. Shown in the lower section, below the parental sequence, are sequencing results for PCR products obtained from genomic DNA of the µ1A knockout (µ1A KO) derivative of the original RPE-1 cell line. Exon 2 contains a 1-base insertion and Exon 4 contains a 2-base deletion, both of which cause frameshifts. ***Middle:*** Confirmation of the mutation in the epsinR gene. The procedure was the same as for µ1A, except that the disrupted gene contains just a single mutation, in this case a 1-base insertion. When epsinR was knocked out in the µ1A KO background, the mutation was a 1-base deletion. ***Right:*** Confirmation of the mutation in the TBC1D23 gene. The procedure was the same as for µ1A, except that the disrupted gene contains just a single mutation, in this case a 5-base deletion. **(B)** RT-qPCR analysis of µ1A and µ1B expression in RPE-1 cells. Two primer pairs were used for each gene. Average Cq values were normalized to GAPDH for each primer pair and biological replicate. 2^-dCq^ was calculated for each primer pair, where dCq is the normalized Cq value. The expression of µ1A and µ1B in RPE-1 cells was calculated relative to the average values for HBE cells. Average values and standard deviations are plotted for four technical replicates from each of three biological replicates. An unpaired t-test indicated that there was no statistically significant difference in µ1A and µ1B expression between RPE-1 and HBE cells. The correct sizes and sequences of the qPCR products from all primer pairs in both cell types were confirmed by agarose gel electrophoreses and DNA sequencing, respectively. **(C)** Immunoblotting of µ1A and µ1B in various cell lines. The numbers in the first lanes represent the sizes in kDa of molecular weight marker proteins. NPM1 was used as a loading control. ***Left:*** Immunoblotting of µ1A in WT cells, µ1A KO cells, and cells expressing endogenously SNAP-tagged µ1A. ***Right:*** Immunoblotting of µ1B in WT RPE-1 cells, WT HBE cells, µ1A KO RPE-1 cells, WT U2OS cells, and µ1A/µ1B double-knockout U2OS cells. For this experiment, the anti-µ1B and anti-NPM1 antibodies were mixed while labeling the blot; the NPM1 band was absent in control experiments that omitted that antibody. The band corresponding to µ1B is visible only in WT RPE-1 cells and WT HBE cells. As marked by the asterisk ("*"), an unidentified protein is visible directly below µ1B, presumably due to cross-reactivity of the antibody even in the non-epithelial U2OS cells that do not express µ1B. **(D)** Confirmation by immunofluorescence and immunoblotting of the cell line with the double knockout of µ1A and epsinR. ***Top:*** Equal numbers of WT and µ1A/epsinR double knockout cells were seeded onto a coverslip and grown overnight. The following day, the cells were processed for immunofluorescence to detect the γ1 subunit of AP-1 (magenta) and epsinR (green). Nuclei were stained with Hoechst dye (blue). In a separate experiment that examined coverslips seeded purely with either WT or µ1A/epsinR double knockout cells, essentially all of the WT cells and none of the mutant cells showed Golgi signals for γ1 and epsinR. That difference is illustrated here with an example of a WT cell next to a µ1A/epsinR double knockout cell. Scale bar, 5 µm. A similar immunofluorescence experiment confirmed the loss of epsinR in the cell line with the single knockout of epsinR. ***Bottom:*** Immunoblotting of epsinR in WT, µ1A KO, and µ1A/epsinR double-knockout cells (*left*) or in WT and epsinR KO cells (*right*). The numbers in the first lanes represent the sizes in kDa of molecular weight marker proteins. GM130 was used as a loading control. **(E)** Confirmation by immunofluorescence and immunoblotting of the knockout of TBC1D23. ***Left:*** Equal numbers of WT and TBC1D23 KO cells were seeded onto a coverslip and grown overnight. The following day, the cells were processed for immunofluorescence to detect TBC1D23 (magenta) and GM130 (green). Nuclei were stained with Hoechst dye (blue). In a separate experiment that examined coverslips seeded purely with either WT or TBC1D23 KO cells, essentially all of the WT cells and none of the mutant cells showed Golgi signals for TBC1D23. That difference is illustrated here with an example of a WT cell next to a TBC1D23 KO cell. Scale bar, 5 µm. ***Right:*** Immunoblotting of TBC1D23 in WT and TBC1D23 KO cells. The numbers in the first lanes represent the sizes in kDa of molecular weight marker proteins. NPM1 was used as a loading control.

**Figure S2.**
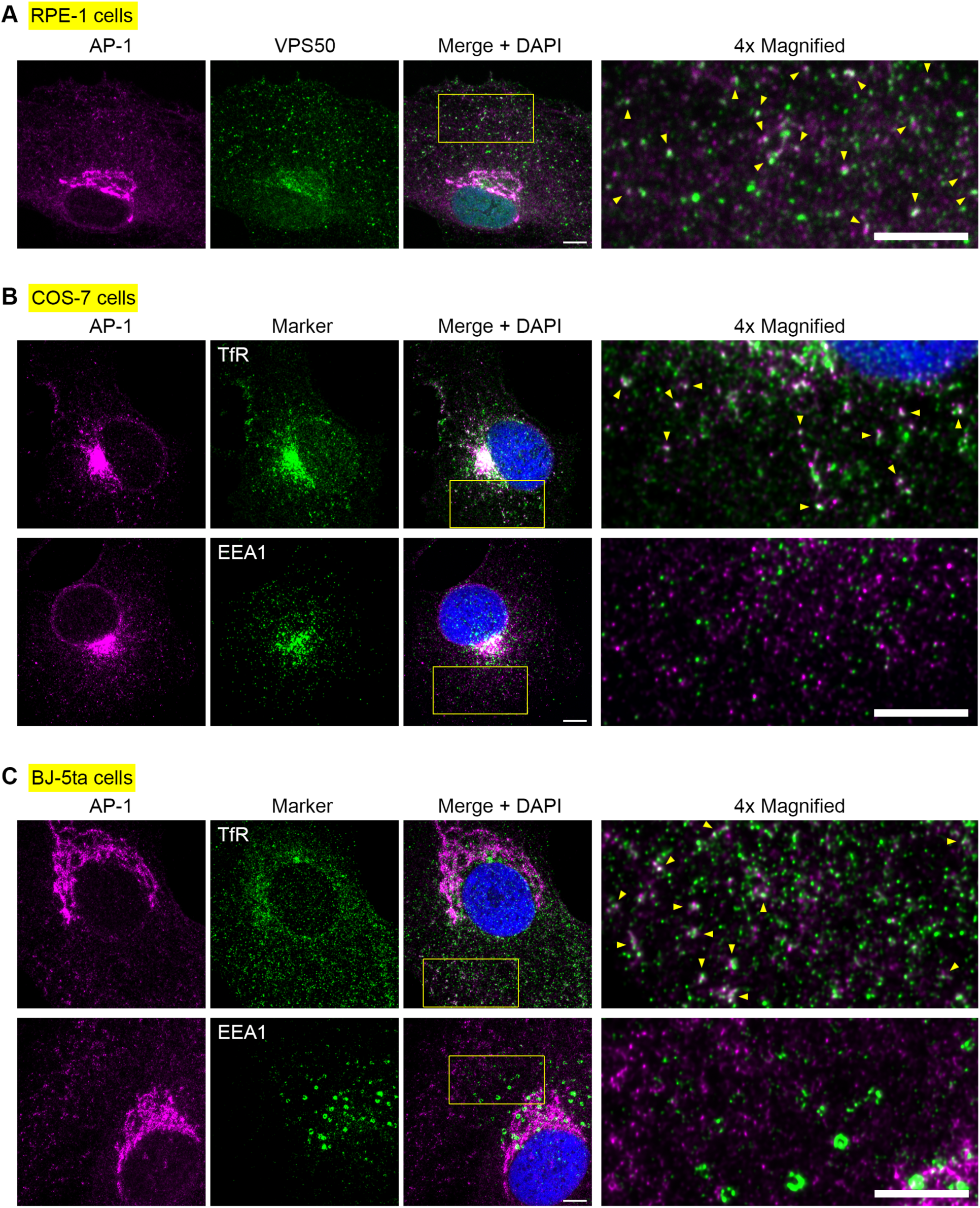
Supplementary data for AP-1 localization. Scale bars are 5 µm. **(A)** Immunofluorescence of AP-1 together with the recycling endosome marker VPS50 in RPE-1 cells. Shown are representative images from an experiment in which the antibody labeling worked well. The γ1 subunit of AP-1 is shown in magenta, VPS50 is shown in green, and DAPI-labeled nuclei are shown in blue. The area in the yellow box is magnified 4x in the right column, where arrowheads in the magnified image indicate examples of structures that contain both γ1 and VPS50. **(B)** Representative immunofluorescence images of AP-1 together with the recycling endosome marker TfR and the early endosome EEA1 in COS-7 cells. The γ1 subunit of AP-1 is shown in magenta, the indicated organelle markers are shown in green, and Hoechst dye-labeled nuclei are shown in blue. The areas in the yellow boxes are magnified 4x in the right column, where arrowheads in the first magnified image indicate examples of structures that contain both γ1 and TfR. **(C)** Representative immunofluorescence images of AP-1 together with the recycling endosome marker TfR and the early endosome EEA1 in BJ-5ta cells. The procedure was as in (B).

**Figure S3.**
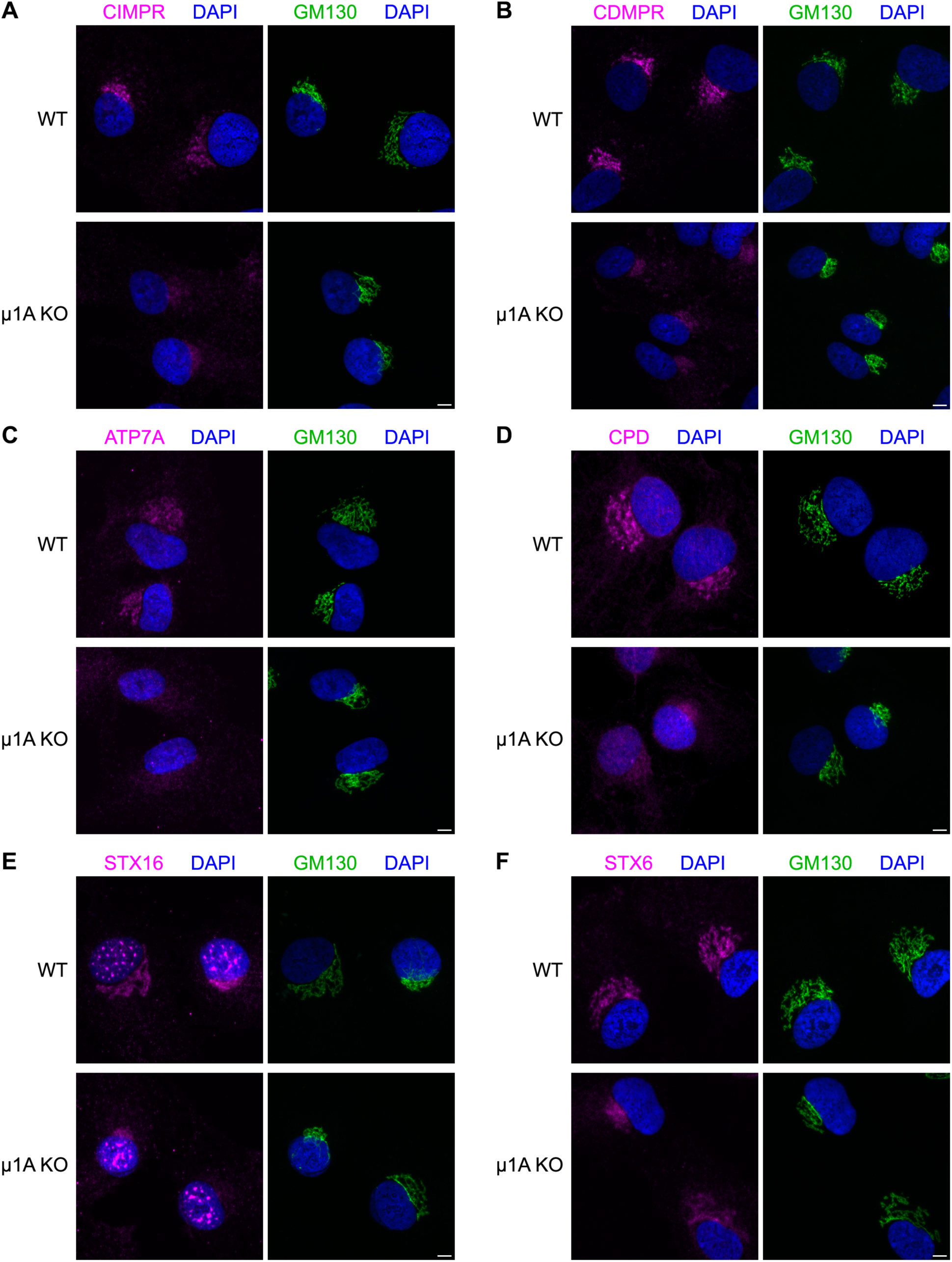
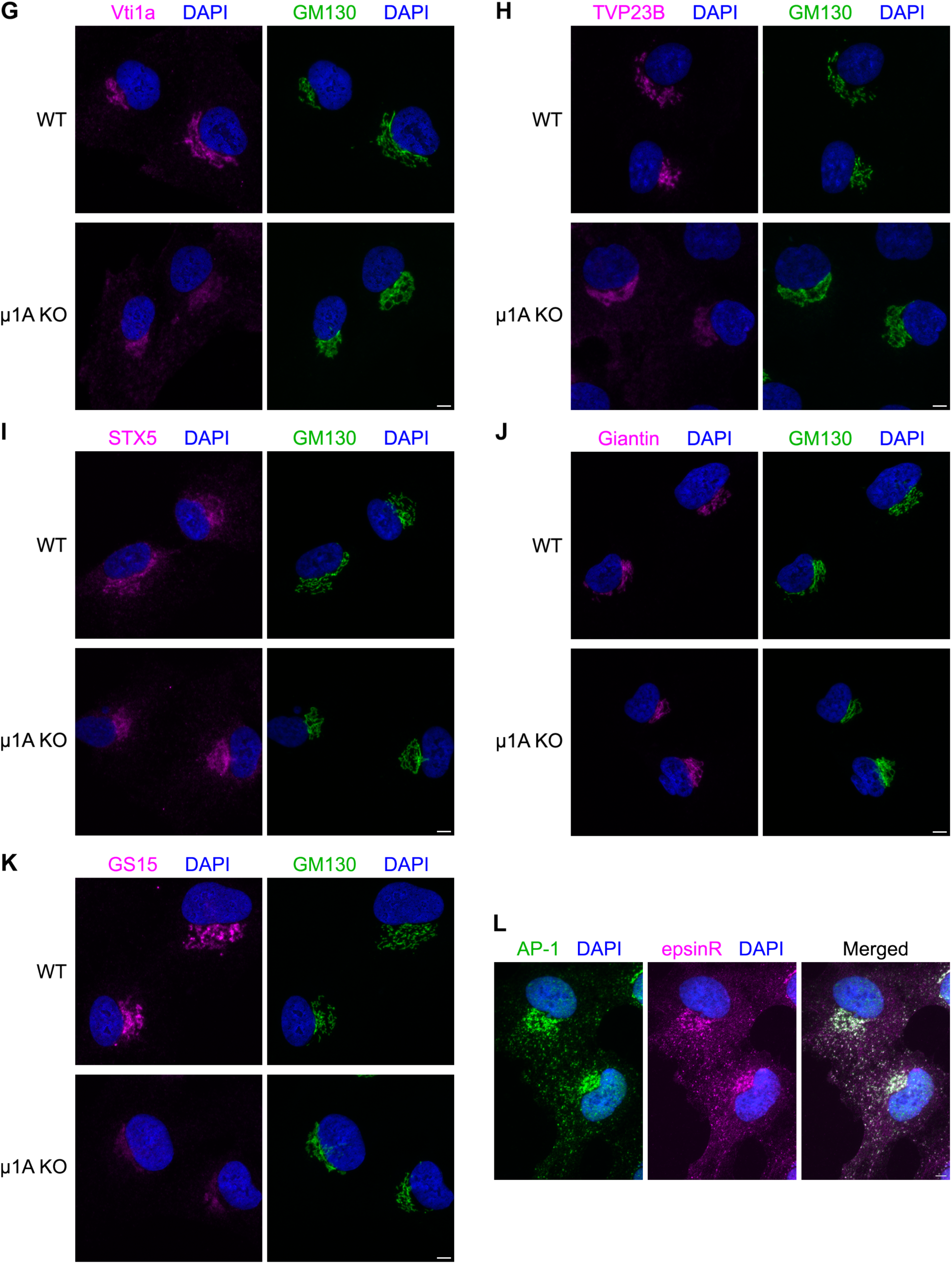
Supplementary data for the effects of knocking out µ1A and for the colocalization of epsinR with µ1A. DAPI-labeled nuclei are shown in blue. Scale bars, 5 µm. Parts (A) - (L) show representative immunofluorescence images of various Golgi proteins (magenta) together with GM130 (green) in WT and µ1A KO cells. Each comparison shows cells imaged in the same experiment using the same parameters. The Golgi proteins examined were (A) CIMPR, (B) CDMPR, (C) ATP7A, (D) CPD, (E) STX16 (for which the antibody cross-reacted with an unknown nuclear component), (F) STX6, (G) Vti1a, (H) TVP23B, (I) STX5, (J) giantin, and (K) GS15. Part (L) shows representative widefield immunofluorescence images of the γ1 subunit of AP-1 compared to epsinR in RPE-1 cells.

**Figure S4.**
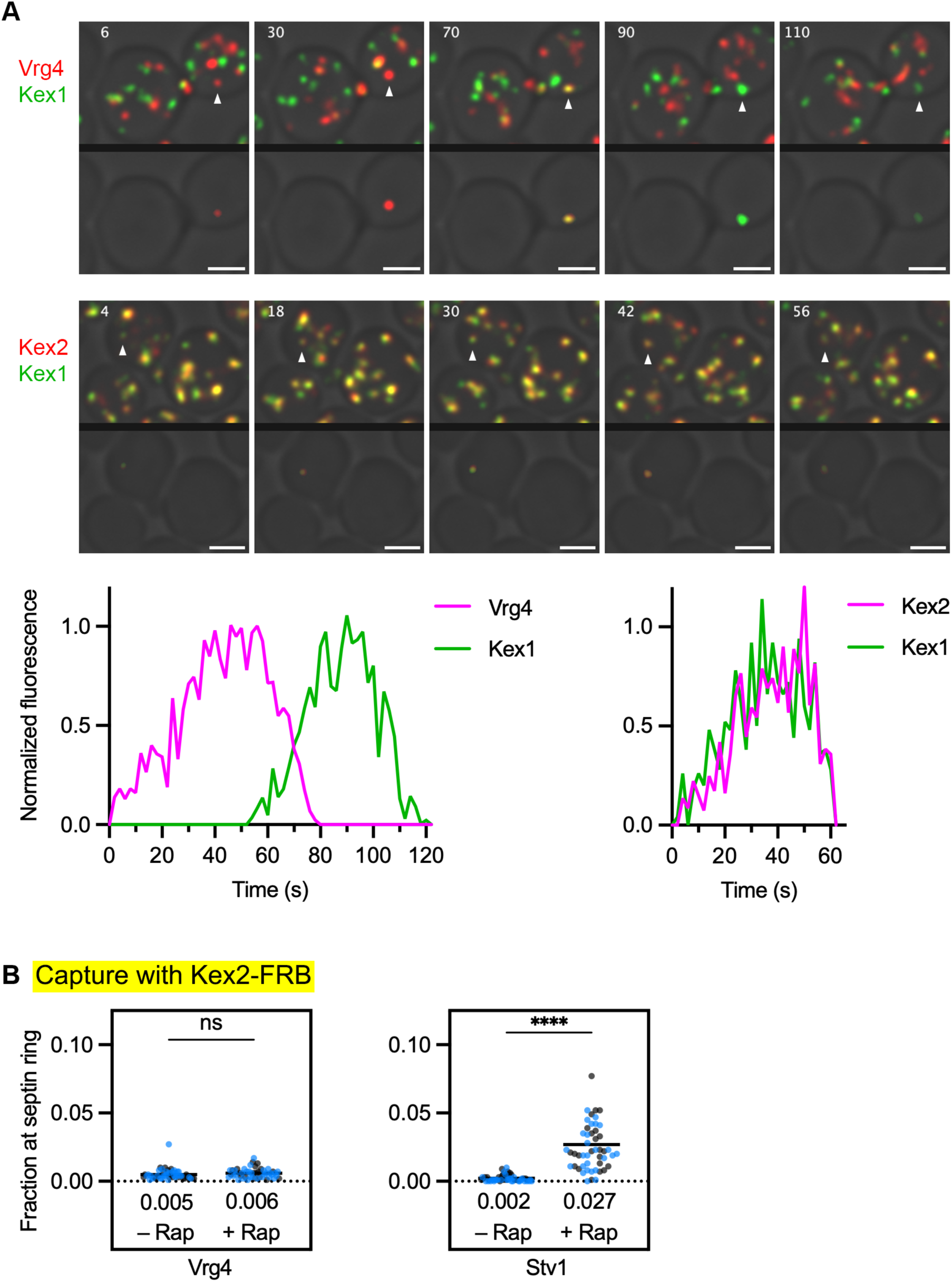
Supplementary data for yeast Kex1. **(A)** Frames from representative 4D confocal movies of HaloTag-Vrg4 plus Kex1-GFP (top) and Kex2-HaloTag plus Kex1-GFP (bottom), and kinetic traces from individual cisternae in the two movies. Depicted are average projected z-stacks at the indicated time points from the two movies contained in Video 5. In each case, the upper row shows the complete projections with arrowheads added to mark the cisterna that was tracked, while the lower row shows edited projections including only the cisterna that was tracked. Scale bars, 2 µm. Plotted below the movie frames are normalized fluorescence intensities for the cisternae tracked in the two movies. **(B)** Control experiments showing that in cells expressing an FKBP-tagged septin, tagging Kex2 with FRB results in no detectable rapamycin-dependent capture of GFP-tagged Vrg4 but strong rapamycin-dependent capture of GFP-tagged Stv1. ns, not significant; ****, significant at P value <0.0001. This analysis was performed in parallel with the one in Figure 7B, and the results are reproduced from Krahn et al. (2026b), originally published in the Journal of Cell Biology (https://doi.org/10.1083/jcb.202509174).

**Video 1.**
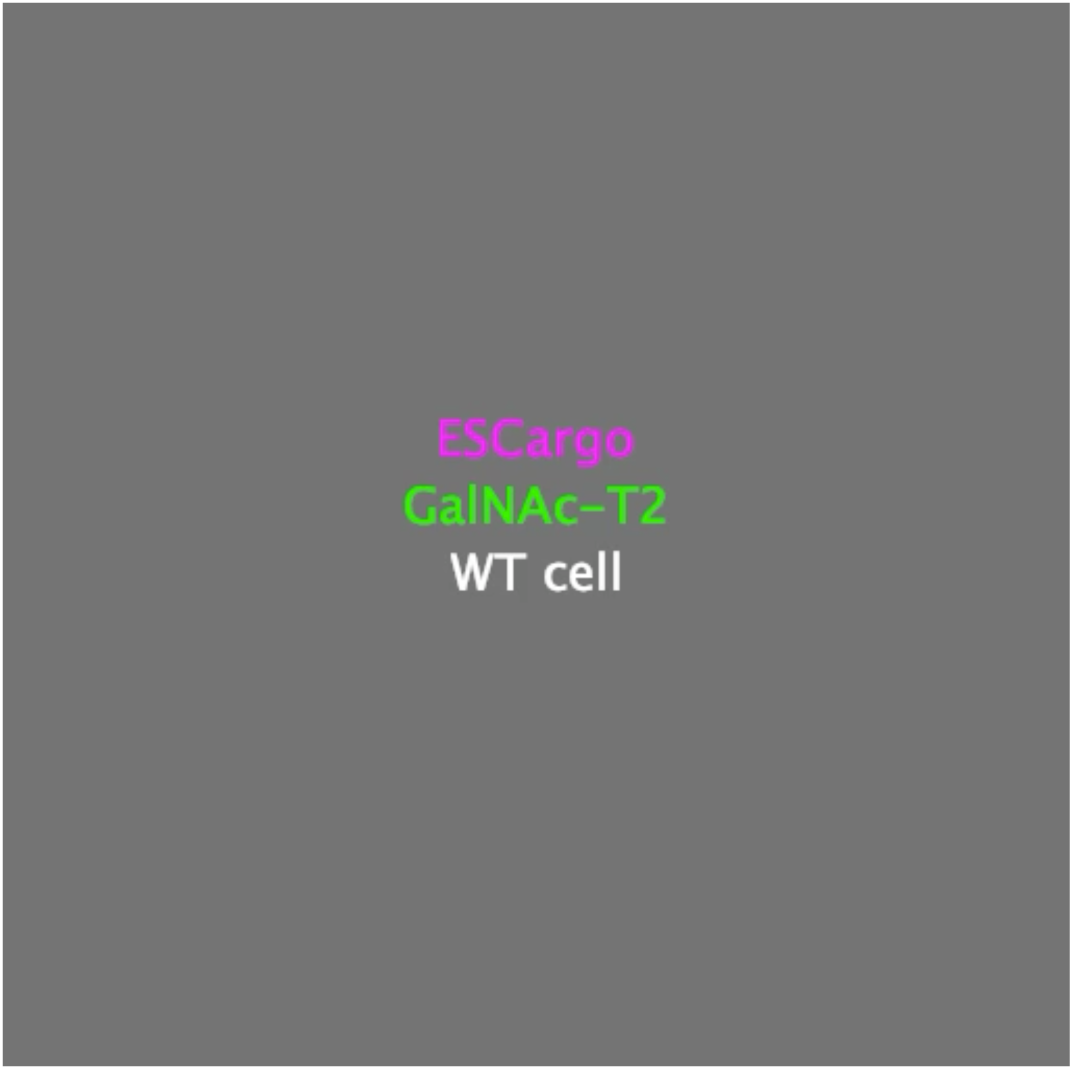
Representative 4D confocal movies of WT and µ1A KO cells showing ESCargo during ER-to-Golgi transport. The first part of the video shows ESCargo transport from the ER to the Golgi in a representative WT cell after the ER-localized aggregates were dissolved by adding SLF for 3 min, and the second part of the video shows the same process in a representative µ1A KO cell. 3D z-stacks for the individual time points were average projected. Intervals between frames are 60 s. Scale bars, 5 µm.

**Video 2.**
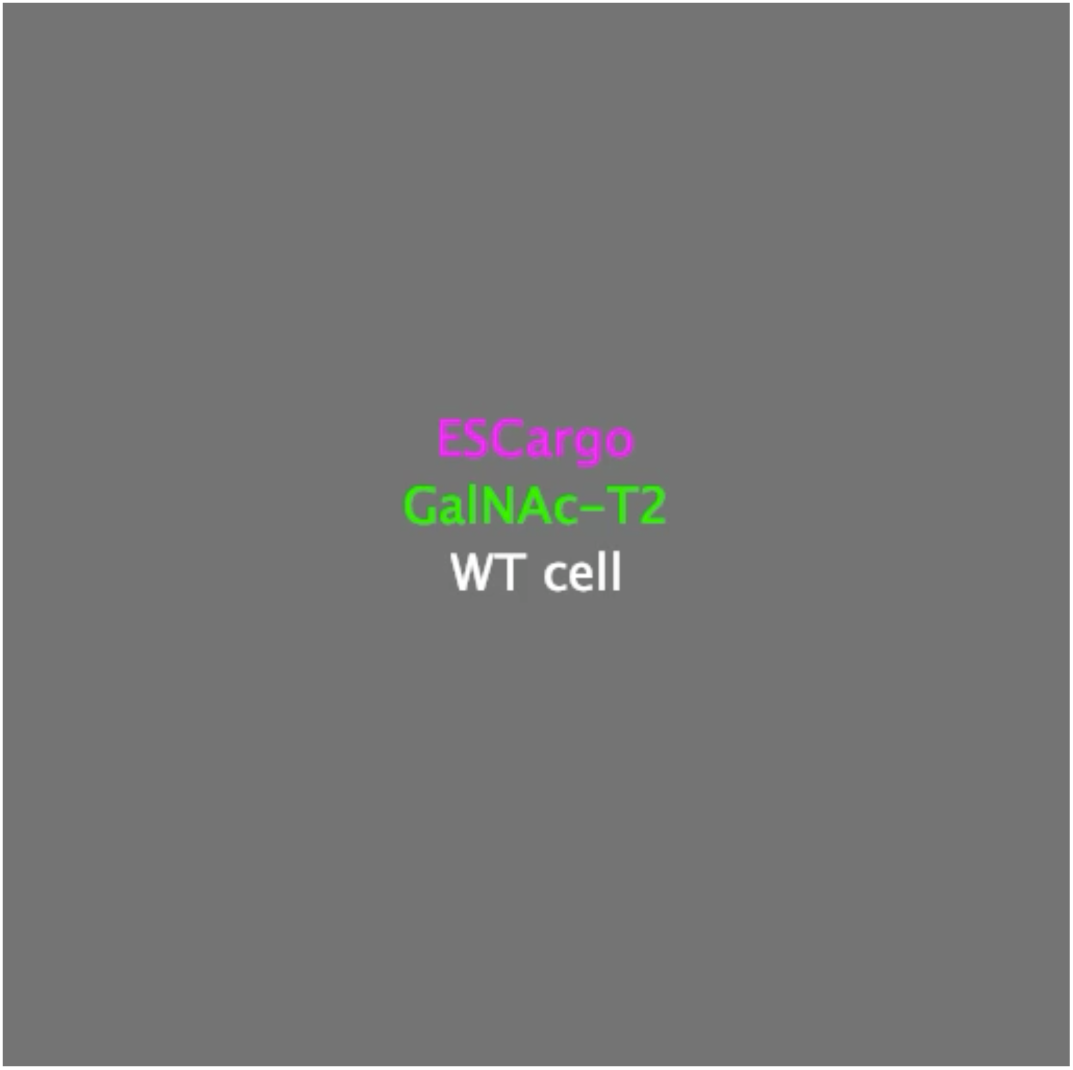
Representative 4D confocal movies of WT and µ1A KO cells showing ESCargo during Golgi export. The first part of the video shows ESCargo export from the Golgi in a representative WT cell after photobleaching the non-Golgi pool, and the second part of the video shows the same process in a representative µ1A KO cell. 3D z-stacks for the individual time points were average projected. Intervals between frames are 60 s. Scale bars, 5 µm.

**Video 3.**
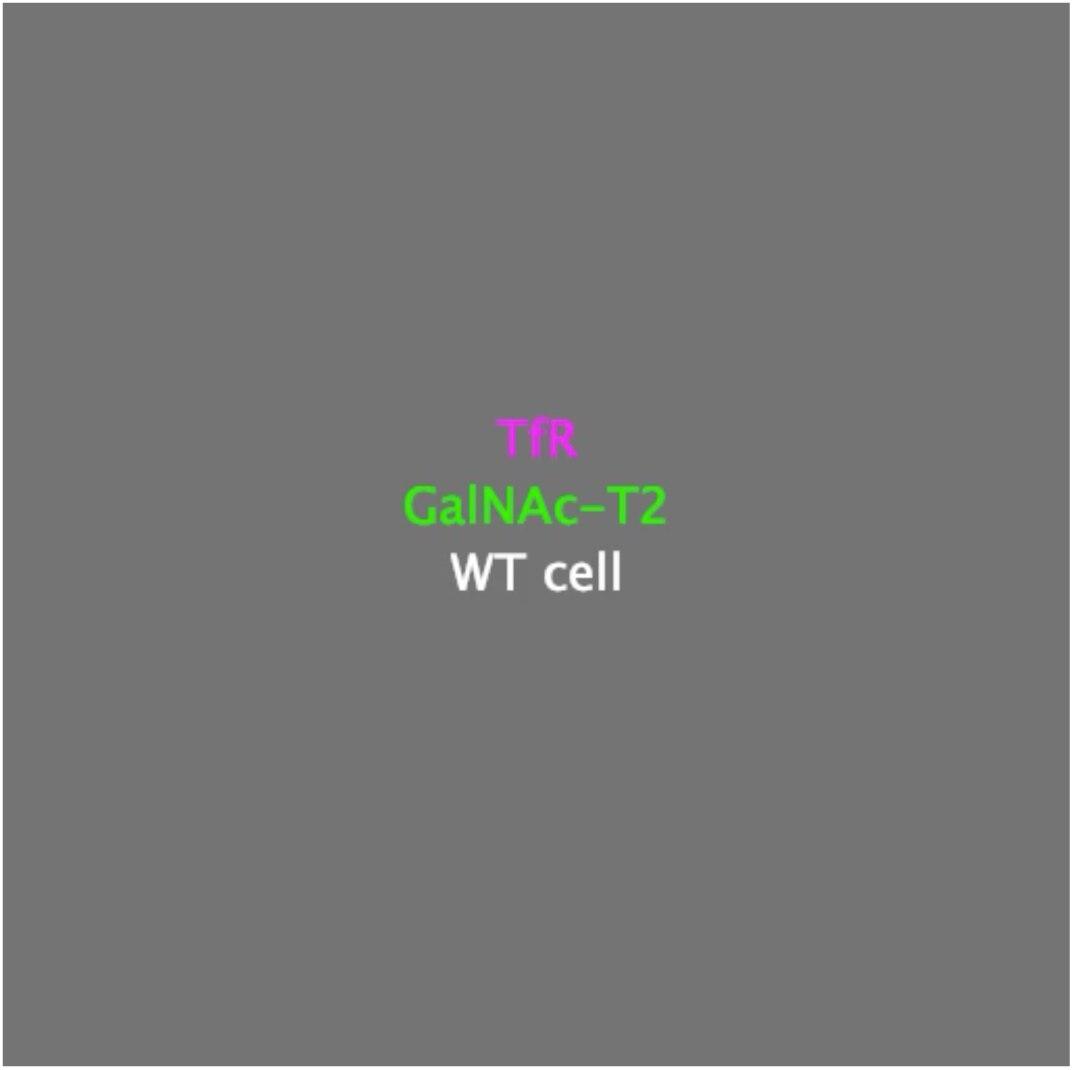
Representative 4D confocal movies of WT and µ1A KO cells showing TfR during ER-to-Golgi transport. The first part of the video shows SBP-mScarlet-TfR transport from the ER to the Golgi in a representative WT cell after the construct was released from the ER by adding biotin, and the second part of the video shows the same process in a representative µ1A KO cell. 3D z-stacks for the individual time points were average projected. Intervals between frames are 60 s. Scale bars, 5 µm.

**Video 4.**
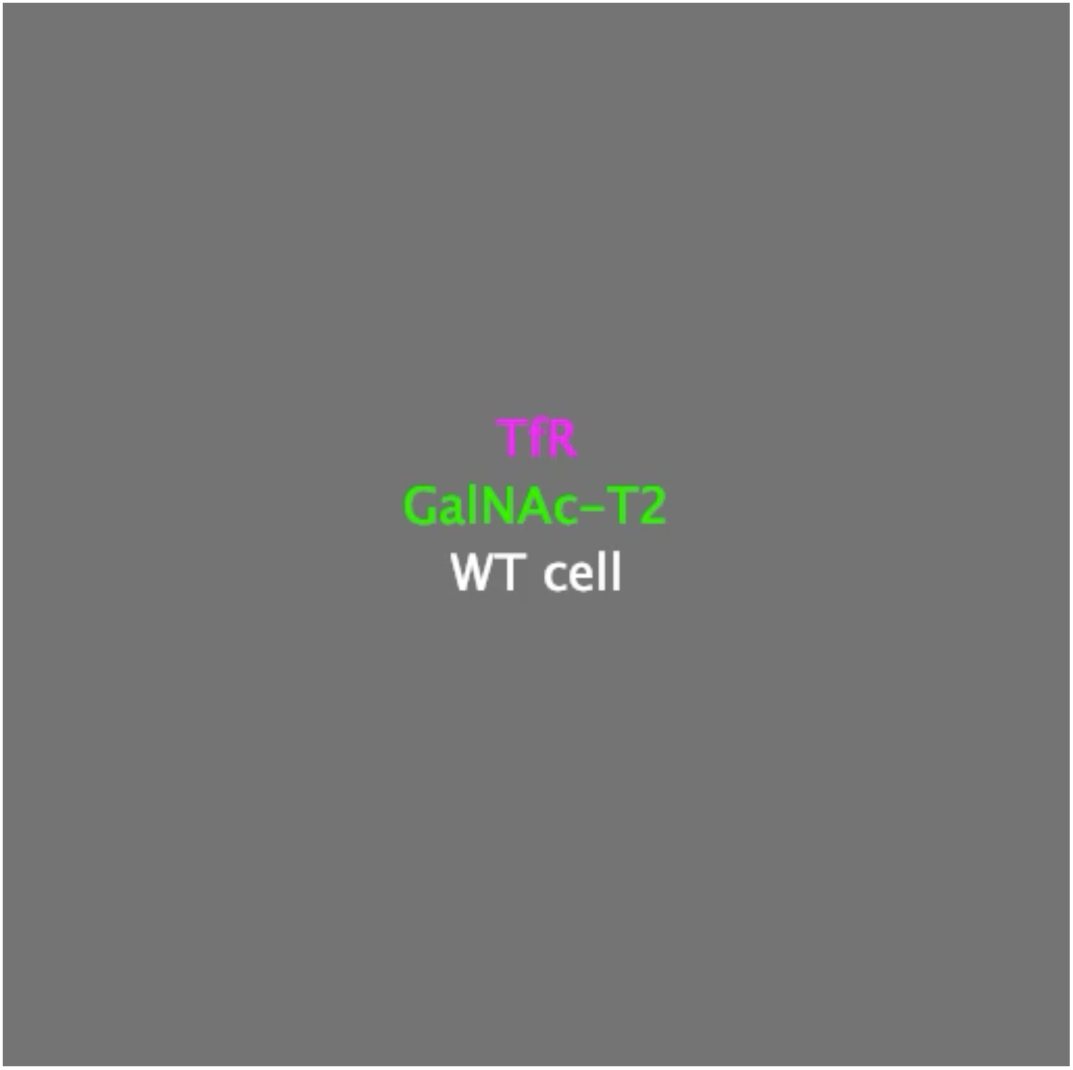
Representative 4D confocal movies of WT and µ1A KO cells showing TfR during Golgi export. The first part of the video shows SBP-mScarlet-TfR export from the Golgi in a representative WT cell after photobleaching the non-Golgi pool, and the second part of the video shows the same process in a representative µ1A KO cell. 3D z-stacks for the individual time points were average projected. Intervals between frames are 60 s. Scale bars, 5 µm.

**Video 5.**
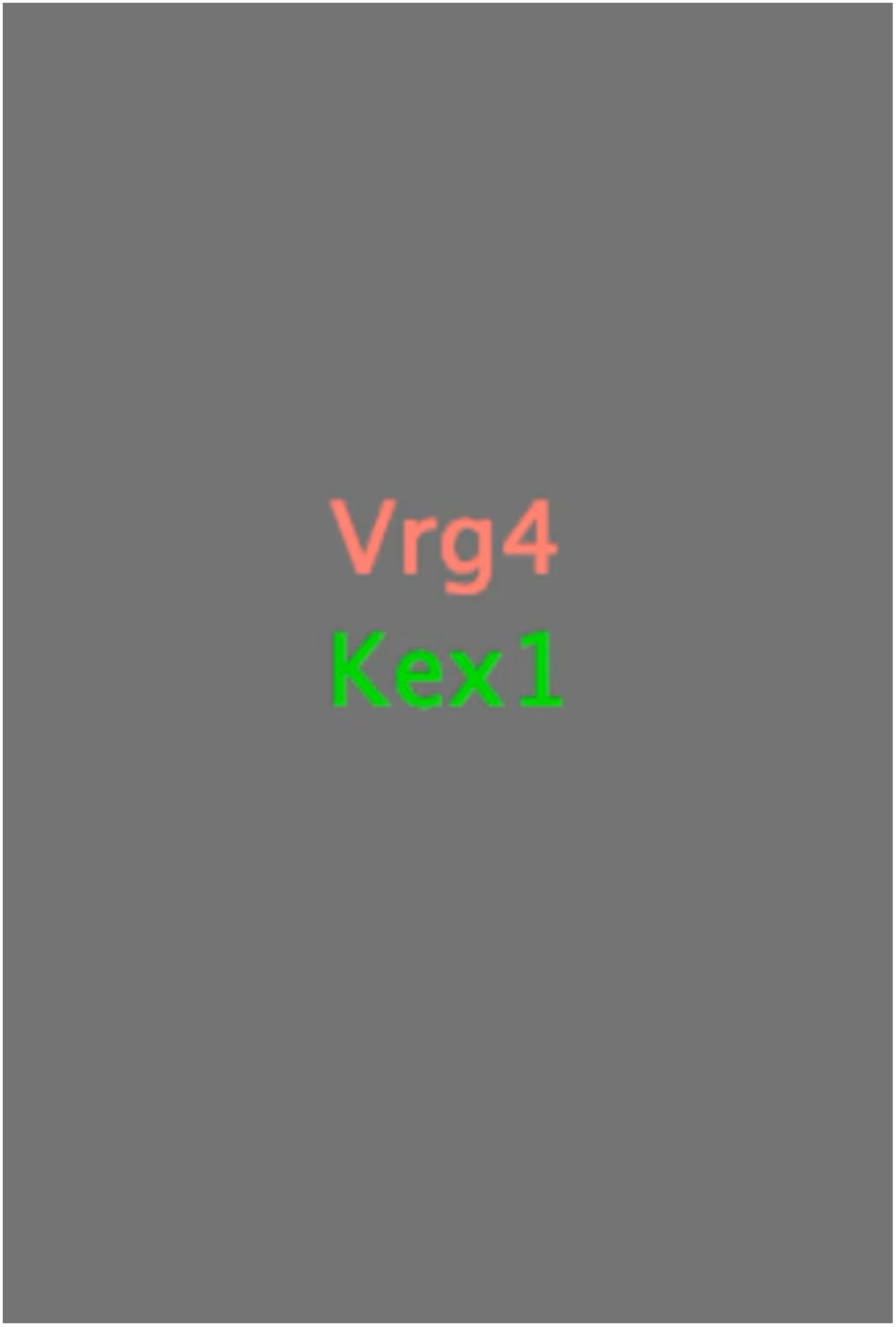
Representative 4D confocal movies of yeast cells showing Kex1 together with an early Golgi marker or a late Golgi marker. The first part of the video shows Kex1-GFP together with HaloTag-Vrg4, and the second part of the video shows Kex1-GFP together with Kex2-HaloTag. 3D z-stacks for the individual time points were average projected. The upper row is the complete projections, and the lower row is edited projections that include only the cisterna that was tracked. Intervals between frames are 2 s. The overlaid numbers represent the time in seconds after the cisterna that was tracked first became detectable. Scale bars, 2 µm. See Figure S4A for further details.

**Video 6.**
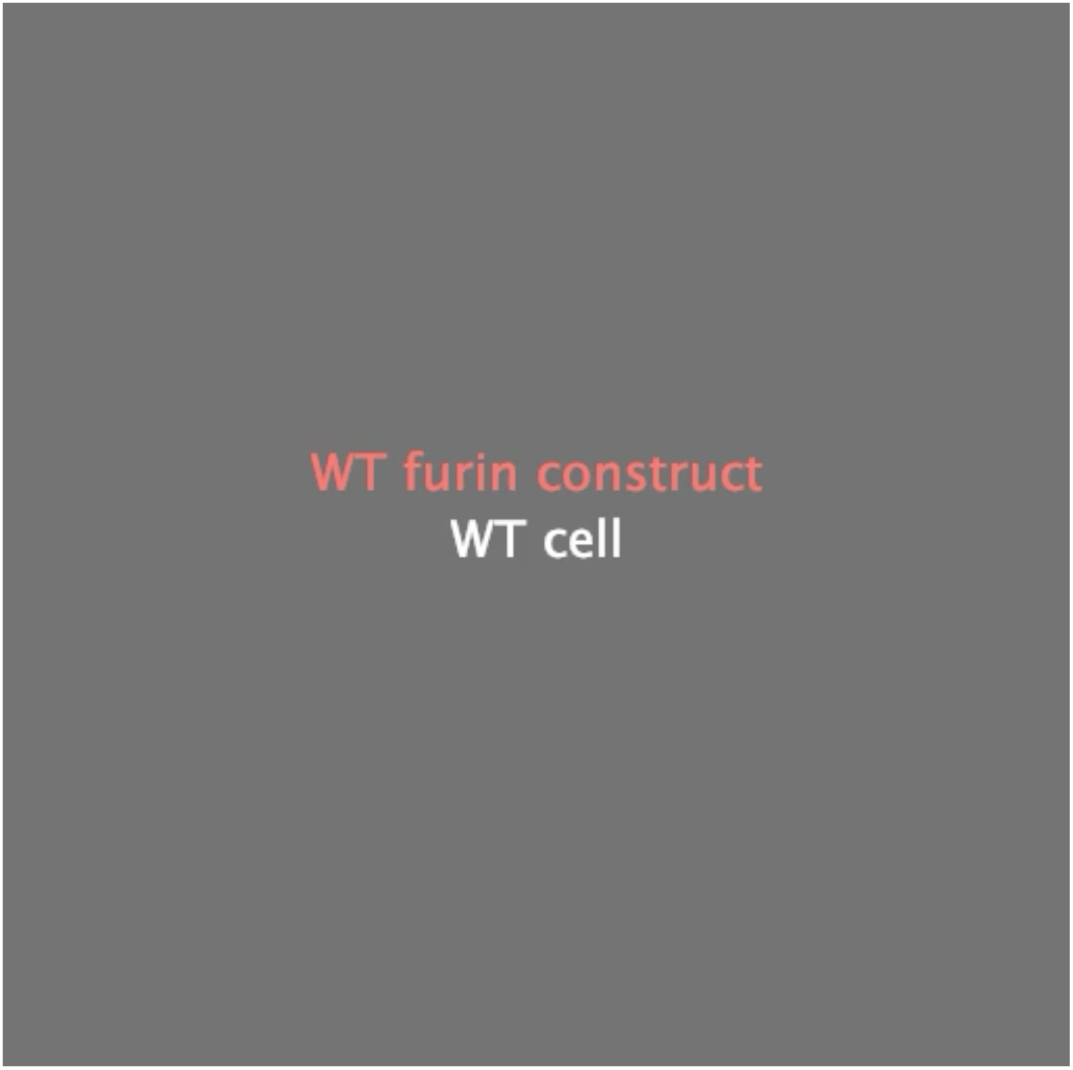
Representative 4D confocal iFRAP movies of WT cells showing wild-type furin or mutant furin lacking the AP-1 recognition signals. The first part of the video shows wild-type SBP-mScarlet-furin remaining at the Golgi in a representative cell after photobleaching the non-Golgi pool, and the second part of the video shows mutant SBP-mScarlet-furin departing from the Golgi in a representative cell after photobleaching the non-Golgi pool. 3D z-stacks for the individual time points were average projected. Intervals between frames are 60 s. Scale bars, 5 µm.

**Video 7.**
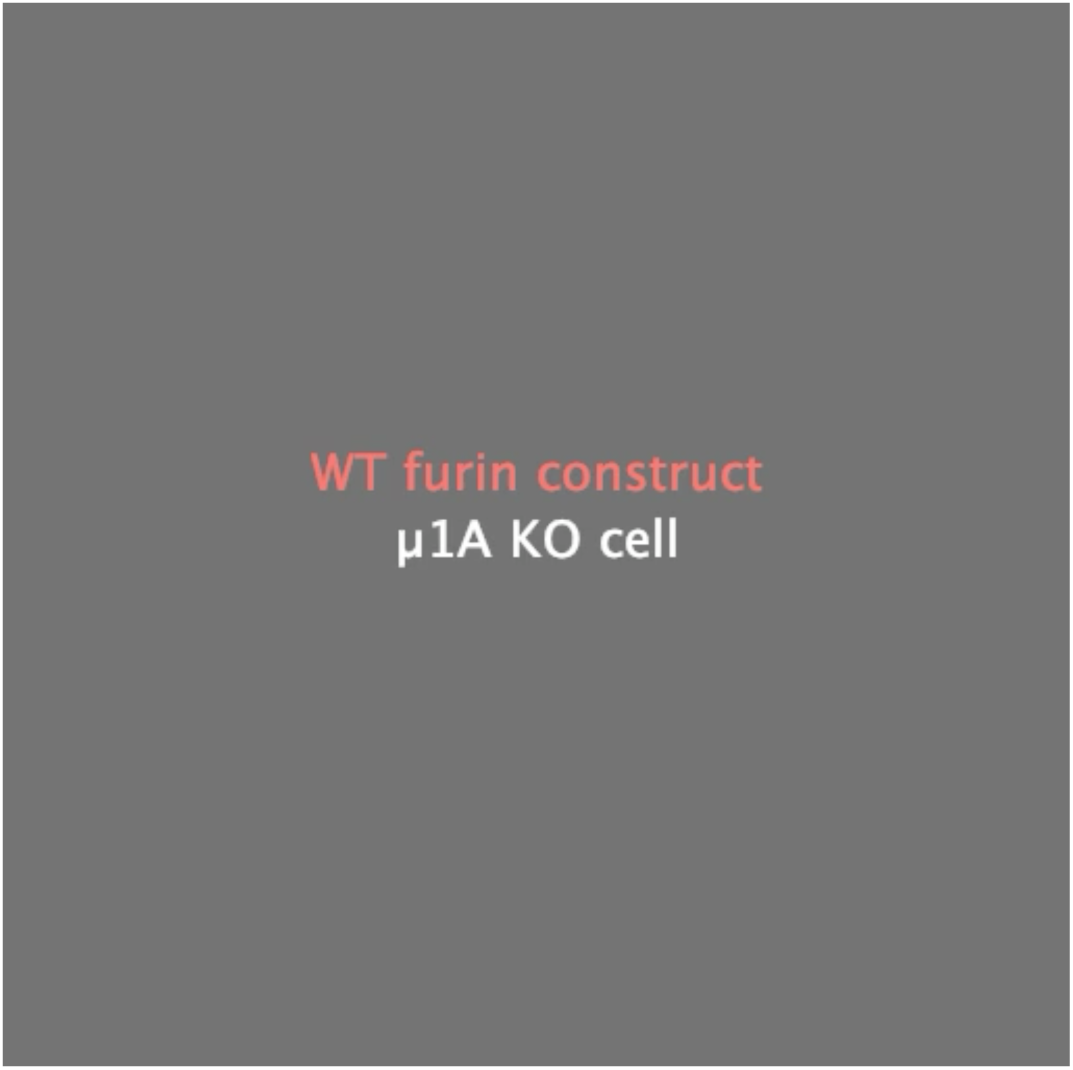
Representative 4D confocal iFRAP movies of µ1A KO cells showing wild-type furin or mutant furin lacking the AP-1 recognition signals. The first part of the video shows wild-type SBP-mScarlet-furin departing from the Golgi in a representative cell after photobleaching the non-Golgi pool, and the second part of the video shows mutant SBP-mScarlet-furin departing from the Golgi in a representative cell after photobleaching the non-Golgi pool. 3D z-stacks for the individual time points were average projected. Intervals between frames are 60 s. Scale bars, 5 µm.

**Video 8.**
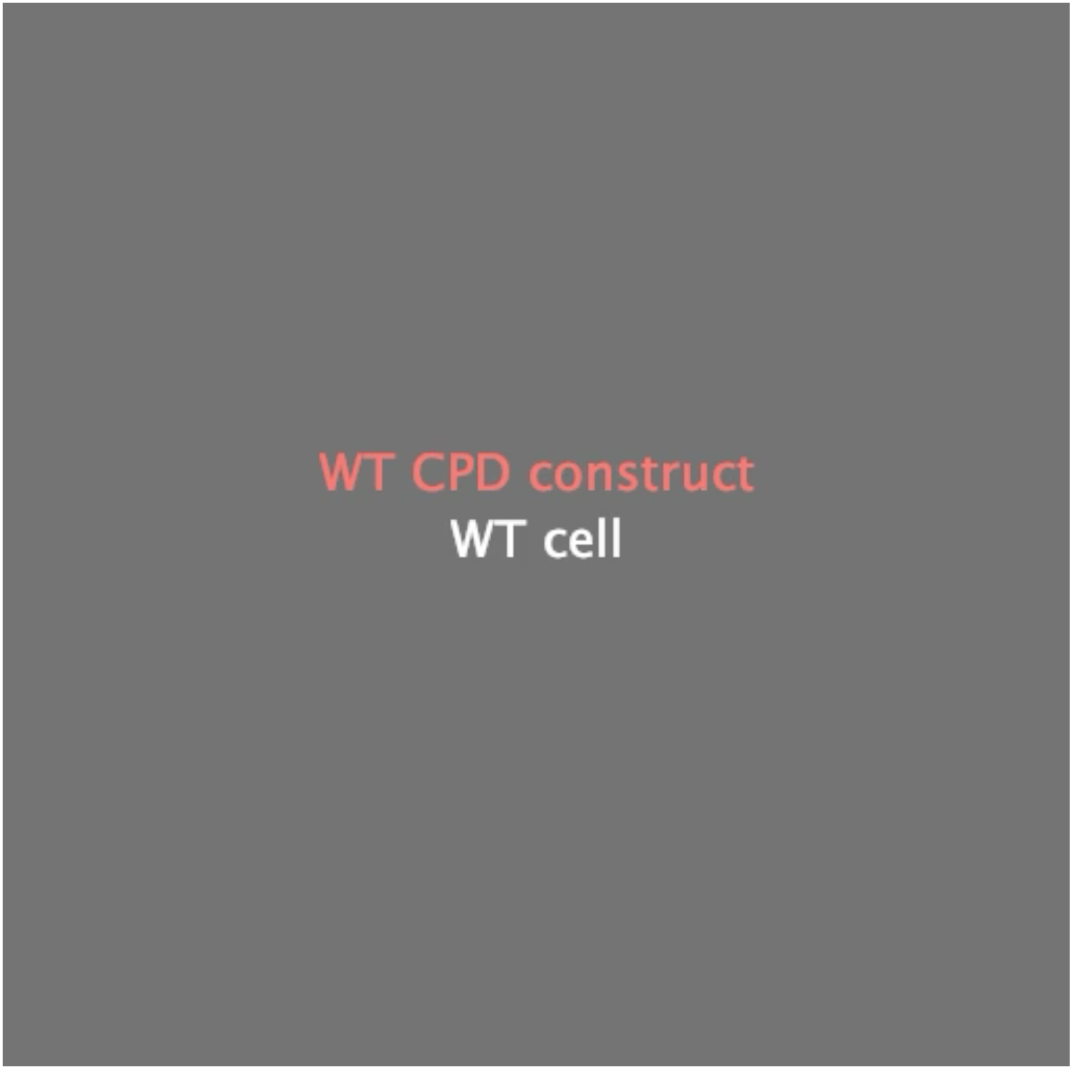
Representative 4D confocal iFRAP movies of WT cells showing wild-type CPD or mutant CPD lacking the AP-1 recognition signals. The first part of the video shows wild-type SBP-mScarlet-CPD remaining at the Golgi in a representative cell after photobleaching the non-Golgi pool, and the second part of the video shows mutant SBP-mScarlet-CPD departing from the Golgi in a representative cell after photobleaching the non-Golgi pool. 3D z-stacks for the individual time points were average projected. Intervals between frames are 60 s. Scale bars, 5 µm.

